# Accurate proteome-wide measurement of methionine oxidation in aging mouse brains

**DOI:** 10.1101/2022.02.19.481142

**Authors:** John Q. Bettinger, Matthew Simon, Anatoly Korotkov, Kevin A. Welle, Jennifer R. Hryhorenko, Andrei Seluanov, Vera Gorbunova, Sina Ghaemmaghami

## Abstract

The oxidation of methionine side chains has emerged as an important posttranslational modification of proteins. A diverse array of low-throughput and targeted studies have suggested that the oxidation of methionine residues in select proteins can have diverse impacts on cell physiology, ranging from detrimental effects on protein structure and stability to functional roles in cell signaling. Despite its importance, the large-scale investigation of methionine oxidation in a complex matrix, such as the cellular proteome, has been historically hampered by technical limitations. Herein we report a methodology, Methionine Oxidation by Blocking (MobB), that allows for accurate and precise quantification of low levels of methionine oxidation typically observed *in vivo*. To demonstrate the utility of this methodology, we applied MobB to the brain tissues of young (6 m.o.) and old (20 m.o.) mice and identified over 280 novel sites for *in vivo* methionine oxidation. We further demonstrated that oxidation stoichiometries for specific methionine residues are highly consistent between individual animals and methionine sulfoxides are enriched in clusters of functionally related gene products including membrane and extracellular proteins. However, we did not detect significant changes in methionine oxidation in brains of old mice. Our results suggest that under normal in vivo conditions methionine oxidation is a biologically regulated process rather than a result of stochastic chemical damage.

## Introduction

Methionine is a normally hydrophobic amino acid with an oxidatively labile thioether group. When oxidized by reactive oxygen species (ROS), methionine forms the hydrophilic amino acid methionine sulfoxide.^1^ For many protein-bound methionines, this reversal of hydrophobicity is believed to have negative consequences for protein structure and stability.^2–4^ For example, it has been shown that the oxidation of surface exposed methionines in GAPDH is sufficient to induce it’s *in vivo* aggregation.^3^ Additionally, it has been suggested that the oxidation of methionine residues in amyloidogenic proteins, such as the prion protein and α-synuclein, may contribute to their misfolding and cytotoxic aggregation.^5, 6^

Although *in vivo* oxidation of methionines has historically been thought of as a stochastic chemical reaction, recent studies have shown that it can also be enzymatically regulated. For example, the molecule interacting with CasL (MICAL) family of monooxygenases (MO) can enzymatically oxidize protein-bound methionines using molecular oxygen as a substrate.^7–9^ MICAL-catalyzed oxidation of methionine residues on F-actin stimulates its depolymerization and regulates cytoskeletal dynamics, establishment of cell shape, vesicle/membrane trafficking and cytokinesis.^10–19^ In addition to the direct enzymatic oxidation of methionine residues, MICAL proteins stimulate the local production of hydrogen peroxide that can indirectly lead to the oxidization of proximal proteins.^20^ Thus, the functional roles of enzymatic methionine oxidation likely extend well beyond F-actin depolymerization.

The biological importance of methionine oxidation is further suggested by the presence of a highly conserved cellular pathway for its chemical reversal. Indeed, although over a dozen different forms of oxidative protein modifications have been described, the oxidation of methionine is one of only a limited number known to have a well-characterized reversal pathway.^21^ Methionine sulfoxides are reduced by the action of a family of enzymes known as methionine sulfoxide reductases (MSRs), acting in concert with thioredoxin and thioredoxin reductases.^22–25^ Thus, the *in vivo* methionine redox cycle is typically described as having and an enzymatic reverse reaction, and a forward reaction that can be either stochastic or enzymatically regulated.

Currently, it is unclear what fraction of methionine sulfoxides observed *in vivo* are a result of regulated enzymatic oxidation, and what fraction are formed by stochastic oxidation events due to random collisions between ROS and methionine side chains. In the former scenario, it would be expected that methionine oxidation would be maintained at specific levels, and perhaps regulated in response to distinct cellular stimuli. In the latter scenario, it would be predicted that methionine oxidation accumulates gradually and sporadically over time, limited by the frequency of random collisions between methionine side chains and ROS. There is significant experimental support for both of these concepts. For example, consistent with the idea of sporadic oxidation, numerous labs have demonstrated that solvent accessibility is a strong predictor of methionine oxidation.^26–29^ Conversely, in support of the regulated oxidation model, a number of studies have presented evidence of specialized cell signaling roles for methionine oxidation.^30, 31^ ^32–38^

The view of methionine oxidation as a form of stochastic protein damage is also supported by the fact that the MSR system is critical for tolerance to oxidative stress in diverse organisms.^39, 40^ These observations, coupled with established associations between the pathways for oxidative stress tolerance and the molecular mechanisms of aging, have led many researchers to hypothesize that methionine oxidation and the MSR pathway play a critical role in aging and the regulation of lifespan.^41, 42^ However, the evidence for the association between MSRs and lifespan in mammalian model systems has been inconsistent.^43, 44^ Furthermore, although some studies have highlighted an increase in the oxidation of specific proteins as a function of aging in certain mammalian model systems^45–49^, the global content of methionine sulfoxides in mouse tissues does not appear to increase as a function of age.^50^ Thus, although bulk levels of methionine sulfoxides may not increase significantly during aging, potentially critical subsets of the proteome may be accumulating oxidative modifications in a site-specific manner. For example, it has been suggested that long-lived proteins with slow turnover rates may be uniquely prone to the accumulation of oxidative modifications during aging.^51–54^

Attempts to clarify the role of methionine oxidation in aging by conducting direct proteome-wide surveys have been historically hampered by technical limitations. Methionine oxidation has been shown to spuriously accumulate, *in vitro,* during the upstream stages of a typical bottom-up proteomics workflows, making it difficult to distinguish methionines that are oxidized *in vivo* from those that are spuriously oxidized *in vitro*.^55, 56^ Although proteome-wide screens for *in vivo* methionine oxidation under conditions of oxidative stress have been successful in characterizing highly oxidized methionines, the quantification of *in vivo* methionine oxidation in the absence of oxidative stress has been mired by high technical variability.^30, 31, 57–59^

In order to address this issue, we and others have previously developed strategies for the proteomic quantification of methionine oxidation that relies on the isotopic labeling of unoxidized methionine residues with H_2_^18^O_2_ during the early stages of sample preparation and prior to LC-MS/MS analysis.^60, 61^ This strategy results in the conversion of all unoxidized methionines to an ^18^O labeled version of the oxidized peptide. Conversely, peptides that are already oxidized *in vivo* retain their ^16^O modifications. The 2 Da mass difference between the ^16^O and ^18^O labeled methionine containing peptides is then used to distinguish between peptides that were unoxidized from those that were oxidized *in vivo*. Although previous proof-of-concept applications of this strategy to proteome-wide analyses showed promise, it lacked the precision needed to quantify the low abundance of methionine oxidation (<5%) typically observed in unstressed cells.^57^

Herein we report an updated methodology, Methionine Oxidation by Blocking (MobB), that allows for the accurate and precise quantification of methionine oxidation in unstressed mammalian tissues. MobB does not require enrichment protocols and is sufficient for the unbiased, large-scale quantification of *in vivo* methionine oxidation stoichiometries. Applying MobB to the brain cortices of young (6 m.o.) and old (20 m.o.) mice, we identify over 280 potentially novel sites for *in vivo* methionine oxidation. Furthermore, we demonstrate that methionine oxidation stoichiometries are biologically reproducible between individual animals, remain stable during aging and are enriched for clusters of functionally related gene ontology terms. Taken together, our results indicate that for a significant subset of the methionine-containing proteome, methionine sulfoxides are tightly regulated and maintained at specific steady-state levels *in vivo* during the course of murine aging.

## Materials and Methods

### Animal handling

All mouse experiments were performed in accordance with guidelines established by University of Rochester Committee on Animal Resources. Male C57BL/6 mice were used in this study and both age groups, young (6 m.o.) and old (20 m.o.), were from University of Rochester colonies where they were housed in a 1-way facility in microisolator housing. Three animals from each group were used. Animals were sacrificed by treatment with isoflurane and perfused with saline prior to organ harvesting. Tissues were flash frozen in liquid nitrogen upon extraction.

### Sample preparation and labeling (oxidation)

Brain cortices were ground in the frozen state using a chilled mortar and pestle. Ground tissues were then resuspended in a denaturing lysis buffer; 50 mM triethylammonium bicarbonate (TEAB) (Fischer Scientific) and 10% sodium dodecyl sulfate (SDS). Homogenization and genomic DNA shredding were achieved by high-energy sonication (Qsonica, amplitude 30, 10s on/60s off), on ice. Samples were clarified of cell debris by centrifugation at 16,000×*g* for 10 min. Protein concentration was quantified by bicinchoninic acid (BCA) assay and immediately diluted (1:1) to a final protein concentration of 0.5 mg/mL with either ^18^O heavy (Sigma) or ^16^O light (Fisher) H_2_O_2_ to a final H_2_O_2_ concentration of approximately 1.25%. Oxidation reactions were allowed to continue for 2 hours at 25°C. Disulfide bonds were reduced by adding 2 mM dithiothreitol (DTT) (Fisher) and alkylated with 10 mM iodoacetamide (IAA) (Sigma). Samples were acidified by adding phosphoric acid to a final concentration of 1.2% and subsequently diluted 7-fold with 90% methanol in 100 mM TEAB. The samples were added to an S-trap column (Protofi), and the column was washed twice with 90% methanol in 100 mM TEAB. Trypsin (Pierce) was added to the S-trap column at a ratio of 1:25 (trypsin/protein), and the digest reaction was allowed to continue overnight at 37°C. Peptides were eluted in 80 *μ*L of 50 mM TEAB followed by 80 *μ*L of 0.1% trifluoroacetic acid (TFA) (Pierce) in water and 80 *μ*L of 50/50 acetonitrile/water in 0.1% TFA. Samples were dried by lyophilization and resuspended in 10 mM ammonium hydroxide to a final concentration of 1ug/uL.

Titration samples were prepared by mixing ^16^O (light)-labeled peptides with ^18^O (heavy)-labeled peptides at prespecified ratios to a final amount of 25ug. In order to increase coverage, titration samples were pre-fractionated on homemade C18 spin columns. Sixteen different elution buffers were made in 100 mM ammonium formate (pH 10) with 2.0, 3.5, 5.0, 6.5, 8.0, 9.5, 11.0, 12.5, 14.0, 15.0, 16.5, 18.0, 19.5, 21.0, 27.0 and 50% acetonitrile added. To reduce the number of samples, fractions were combined in the following ways: 1-9, 2-10, 3-11, 4-12, 5-13, 6-14, 7-15, 8-16. All fractions were then lyophilized and resuspended in 12.5 *μ*L of 0.1% TFA.

### LC-MS/MS analysis

Fractionated peptides were injected onto a homemade 30 cm C18 column with 1.8 *μ*m beads (Sepax), with an Easy nLC-1200 HPLC (Thermo Fisher), connected to a Fusion Lumos Tribrid mass spectrometer (Thermo Fisher). Solvent A was 0.1% formic acid in water, while solvent B was 0.1% formic acid in 80% acetonitrile. Ions were introduced to the mass spectrometer using a Nanospray Flex source operating at 2 kV. The gradient began at 3% B and held for 2 min, increased to 10% B over 5 min, increased to 38% B over 68 min, then ramped up to 90% B in 3 min and was held for 3 min, before returning to starting conditions in 2 min and re-equilibrating for 7 min, for a total run time of 90 min. The Fusion Lumos was operated in data-dependent mode, with MS1 scans acquired in the Orbitrap and MS2 scans acquired in the ion trap. The cycle time was set to 1.0 s to ensure that there were enough scans across the peak. Monoisotopic precursor selection (MIPS) was set to Peptide. The full scan was collected over a range of 375–1400 *m*/*z*, with a resolution of 120 000 at an *m*/*z* of 200, an AGC target of 4e5, and a maximum injection time of 50 ms. Peptides with charge states between 2 and 5 were selected for fragmentation. Precursor ions were fragmented by collision-induced dissociation (CID) using a collision energy of 30% with an isolation width of 1.1 *m*/*z*. The ion trap scan rate was set to rapid, with a maximum injection time of 35 ms and an AGC target of 1e4. Dynamic exclusion was set to 20 s.

### MaxQuant analysis and data conversion

Raw files for all samples were searched against the *Mus musculus* UniProt database (downloaded 5/3/2021) using the integrated Andromeda search engine with the MaxQuant software. Peptide and protein identifications were performed with MaxQuant using the default parameter settings. ^18^O Methionine sulfoxide, ^16^O methionine sulfoxide, and N-terminal acetylation were set as variable modifications, and carbamidomethyl cysteine was set as a fixed modification. Raw files were converted to the mzXML format with the ProteoWizard’s MSConvert software using the vendor-supplied peak picking algorithm, no additional filters were used. The MaxQuant-supplied evidence file and the mzXML file were used as the input into a custom algorithm described below. All raw and processed data are available at ProteomeX-change Consortium via the PRIDE database (accession number PXD014770).

### Custom quantification of ^16^O/^18^O labeled methionine ratios

Quantification of ^16^O/^18^O labeled methionine ratios (MOS values) was done using an in-house algorithm, described in detail in *Supplementary Materials and Methods* and available for download as a supplementary file. Briefly, ^16^O/^18^O labeled methionine pairs are subset from the MS1 spectra using predicted retention times and mass to charge ratios. A gaussian, or mixture of gaussians, model is then fit to the RAW data and consecutive isotopologues are connected based on cosine similarities. Estimated ^16^O/^18^O labeled methionine ratios are measured as the slope of a linear regression between the summed intensities of light and heavy labeled isotopologues. Finally, estimated ^16^O/^18^O labeled methionine ratios are corrected based on the theoretical overlap between the isotopic envelopes of light and heavy labeled peptides.

### Statistical analysis

One sample analysis was used to identify methionines that were methionine significantly more oxidized than the global median oxidation of methionine residues, we have named these methionines oxidation-prone methionines. Significance was assigned using an adaptive shrinkage model as described in the *ashr* package of R.^62^

Effect sizes, or inter-age MOS values, were calculated using all available data on each unique peptide sequence. Peptide-specific titration response curves were generated as described above. The inter-age MOS value for each peptide (effect size) and associated standard error are estimated parameters returned by the models used to fit titration response curves. Prior to analysis by an adaptive shrinkage model inter-age MOS values were reformatted as distances from the global median. A half-uniform model with only positive effects was used. Significance was assigned on the criteria of q-value <= 0.01.

Two sample analysis was used in order to identify age-specific differences in the proteomic distribution of methionine oxidation. Age-specific titration response curves were generated as described above, using all available data collected on either young (6 m.o.) animals or old (20 m.o.) animals.

Effect sizes were measured as the difference in estimated *in vivo* MOS values between old and young age groups. Pooled variances were calculated using the standard error in estimated in vivo MOS values for each age group. Effect sizes and pooled variances were used to calculate a t-test statistic and p-value for each peptide. Corrections for multiple hypothesis testing were done using a Holm-Bonferroni method.

### Comparisons between methionine oxidation and intrinsic properties of proteins and methionines

Methionine solvent accessible surface areas (SASA) were calculated in pymol using an in-house python script and databank of publicly available alphafold structures. Protein turnover rates were taken from publicly available datasets.^63, 64^ All turnover rates are for proteins measured in the brain cortices of wildtype mice. Protein abundances were taken from a publicly available dataset on the brain cortex of wildtype mice.^63^ Methionine solvent accessible surface area, protein turnover rates and protein abundances were grouped into their respective categories using ggplot2 in R.

Significance was tested by two different methods. First, a pairwise t-test was used to compare the global means of in-vivo MOS values between the three categories of each bioinformatic parameter (SASA, protein turnover rates and abundance). Second a chi-squared test was used to test for a significant association between oxidation-prone methionines and the three categories of each bioinformatic parameter.

Sequence analysis was done using icelogo. The three amino acids N-terminal to methionine and the three amino acids C-terminal to methionine were taken from protein sequences in the *Mus musculus* UniProt database (downloaded 5/3/2021).

### Geno ontology (GO) analysis

Proteins represented by at least one oxidation-prone methionine (target) were compared by to all other proteins quantified in the assay (background). In order for a protein to be considered quantified in the assay it must have been identified in at least seven out of a possible twelve samples for both age groups. The list of background proteins included proteins identified by non-methionine containing proteins. Peptides were annotated for GOterms and tested for significance using Perseus software and databases. A cutoff of 0.05 for the Bonferroni-corrected *p*-values was used to determine significantly enriched terms. Enriched terms were clustered based on semantic and functional similarity.

## Results

### Proteomic quantification of methionine oxidation stoichiometries by MobB

MobB is an improved proteomic workflow, based on our previously published methods.^57^ An overview of the experimental strategy is illustrated in Figure 1. Briefly, the accumulation of *in vitro* methionine oxidation events during sample preparation may result in an overestimation of *in vivo* oxidation stoichiometries. We circumvent this problem by forced oxidation of methionines with ^18^O-labeled hydrogen peroxide (H_2_^18^O_2_) shortly after extraction (Figure 1A). The 2 Da mass difference between the ^16^O and ^18^O-labeled methionine containing peptides is then used to distinguish between peptides that were oxidized (^16^O) from those that were unoxidized (^18^O) *in vivo*. Spike-in carrier proteomes of fully ^16^O-labeled peptides are used to measure titration response curves and extrapolate *in vivo* methionine oxidation stoichiometries (MOS) values, discussed further in Supplementary Materials.

**Figure 1.**
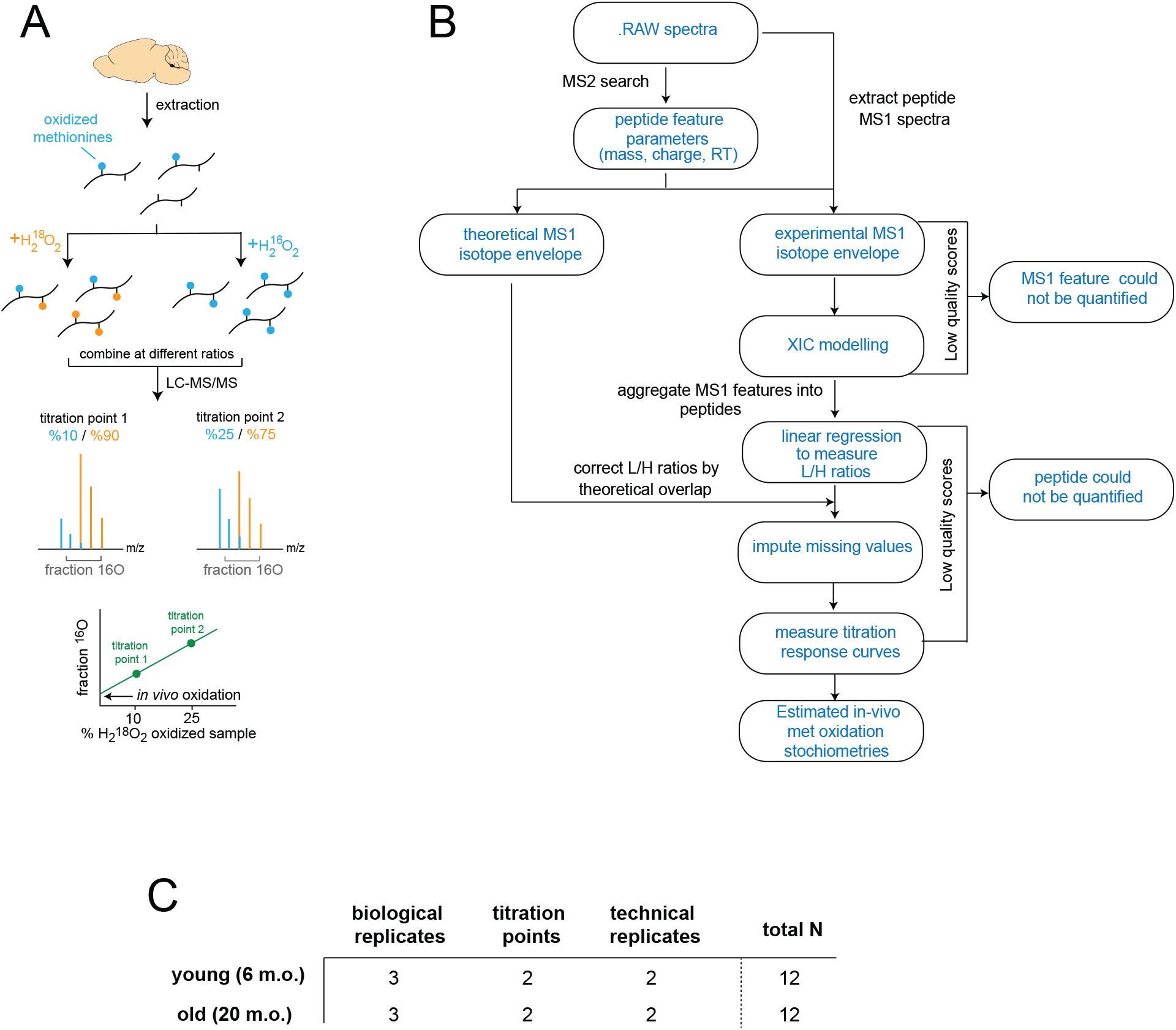
Experimental design and MobB workflow. (A) Schematic overview of ^18^O blocking and titration methodology. Biological extracts that contain variable levels of ^16^O oxidized methionines (blue) are fully oxidized with ^18^O and ^16^O containing hydrogen peroxide. ^18^O-labeled methionines are shown in orange. The two oxidized samples are mixed at predefined ratios and analyzed in a bottom-up LC-MS/MS workflow. Relative levels of ^16^O and ^18^O oxidized methionine-containing peptides are measured by quantifying the MS1 spectra. The resulting titration response (bottom) can be used to estimate *in vivo* methionine oxidation stoichiometries. (B) A general overview of the computational strategy used to quantify the relative ratios of ^16^O to ^18^O labeled peptides and estimate *in vivo* methionine oxidation stoichiometries. (C) Experimental design of this study. Brain cortices from three biological replicates of young and old mice were assayed for methionine oxidation stoichiometries using the MobB workflow. Two titration points, 10% (low) and 25% (high) were used to quantify titration response curves. Two technical replicates of each titration point were used for a total of 12 samples per age group.

The 2 Da mass difference between ^16^O and ^18^O labeled peptides is not sufficient to fully separate the isotopic envelopes of the labelling pair and MobB makes use of a custom algorithm to quantify the resulting irregular isotopic envelopes of ^16^O and ^18^O labeled methionine containing peptide pairs (Figure 1B). Previous attempts at applying a similar strategy to the proteome-wide quantification of MOS in unstressed cultured cells reported a generally low abundance of ^16^O-labeled peptides that could only be quantified with low precision.^57^ MobB improves upon this strategy with novel additions to the computational workflow, discussed individually in Supplementary Materials, aimed at improving (i) precision, (ii) coverage and (iii) false discover rate (FDR).

In this study, we applied the MobB workflow to quantify the proteome-wide distribution of *in vivo* MOS values in the brain cortices of young (6 m.o.) and old (20 m.o.) mice. Our experimental design included two isotope titration points; one prepared with a fully ^16^O labeled carrier proteome at 10% and another at 25%, two technical replicates of each titration point and three biological replicates of each age group, young and old (Figure 1C).

In order for a peptide to be considered quantified in the current study, we required a minimum of seven (out of a possible twelve) valid values for each age group. In total, after quality control filtering, 1232 methionine-containing peptides mapped to 629 distinct proteins were quantified in the final dataset. Compared to previous versions of this methodology, which utilized a computationally expensive target-decoy strategy for FDR control, MobB has a simplified and more user-friendly approach to FDR control. As can be seen in Figure 1B, MobB includes user defined quality cutoffs at multiple stages, allowing for the filtering of both low quality MS1 peptide features and peptide sequences that are not quantifiable. A complete discussion of the quality cutoff scores used in this study can be found in Supplementary Materials.

### MObB measurements are precise

As can be seen in Figure 2A, MobB estimates of *in vivo* MOS values in brain cortices of both young and old mice are highly precise, with an average peptide-level standard error (SEM) of +/- 1%. In addition, MOS values in both titration points could be measured with high precision. In the case of the former, SEM values were estimated by fitting a peptide-specific titration response curve to the data (N=12) and in the case of the latter, SEM values were estimated from the distribution of measurements made on technical and biological replicates (N=6). The quantitative precision of MobB is a significant improvement upon previous versions of this methodology, allowing for meaningful quantification of evenly lowly oxidized methionines.

**Figure 2.**
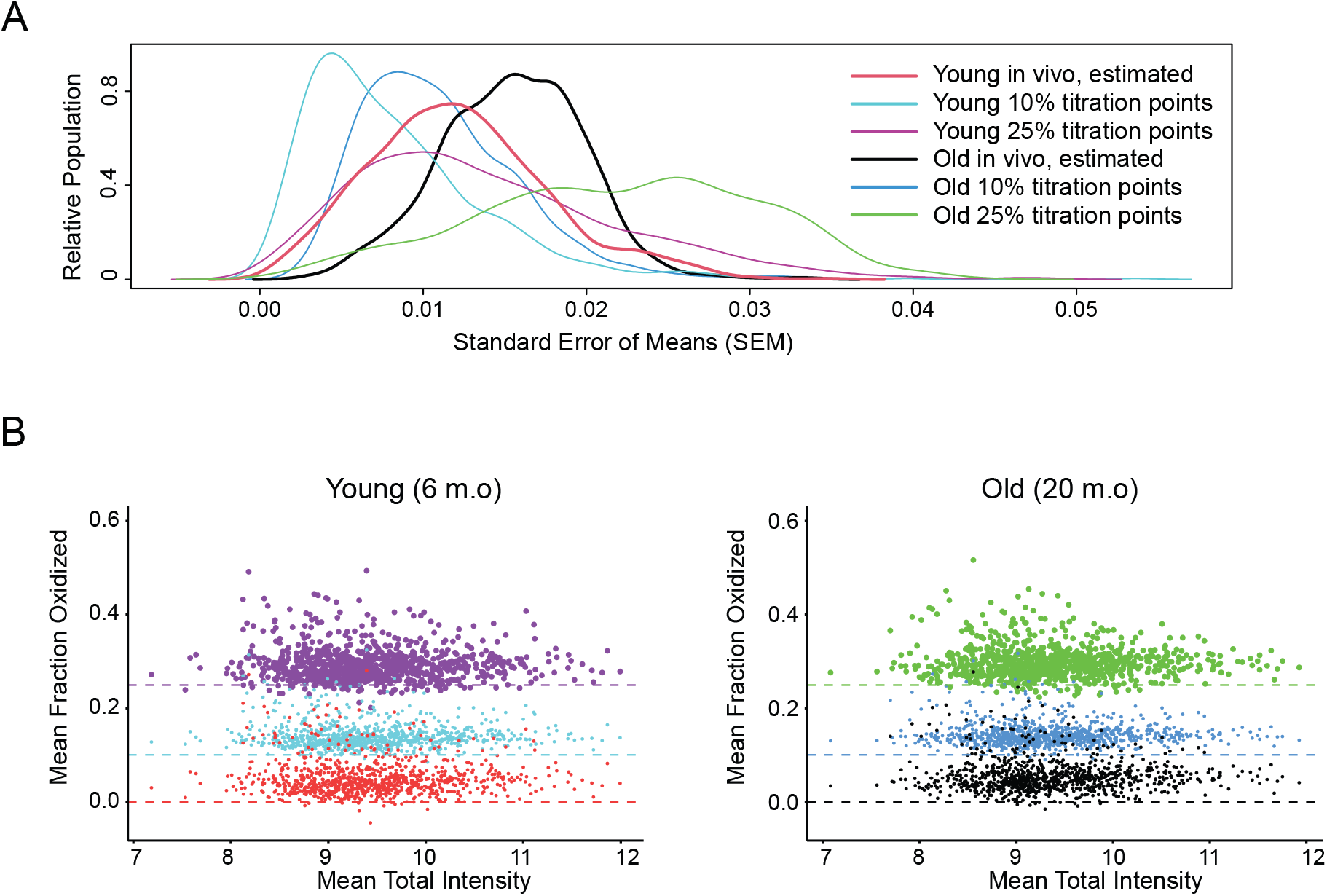
The proteome-wide precision of MobB measurements. (A) Density plots showing the distribution of standard errors associated with each experimental grouping. Distributions associated with *in vivo* estimates (black, red) are the SEMs associated with parameter estimates when fitting experimental data to a titration response curve. All other distributions are measured as the SEM associated with distributions of experimentally observed values. (B) Scatter plots comparing the average total MS1 intensity of a methionine sulfoxide containing peptide to its average measured light to heavy ratio (methionine oxidation stoichiometries) in young (left) and old (right) mouse brain cortices. Colors are as in (A).

Furthermore, as can be seen in Figure 2B, for each titration point we observe tight clustering about a global mean and no intensity bias in our measurements. Taken together these results suggest that MobB measurements are precise and sufficient for the unbiased estimation of *in vivo* MOS values (Figure 2b; red, black).

### Methionine oxidation stoichiometries are generally low in the brain cortices of both young and old animals

Previous studies on enriched or purified single-proteins and synthetic peptides utilizing an identical labelling protocol suggested that isotopic impurity of the ^18^O blocking reagent is negligible and therefore baseline oxidation measurements for blocked peptides are approximately zero.^36^ Therefore, our results indicate that in the brain cortex the average protein-bound methionine is marginally oxidized (∼4.5%) in both age groups (Figure 3).

**Figure 3.**
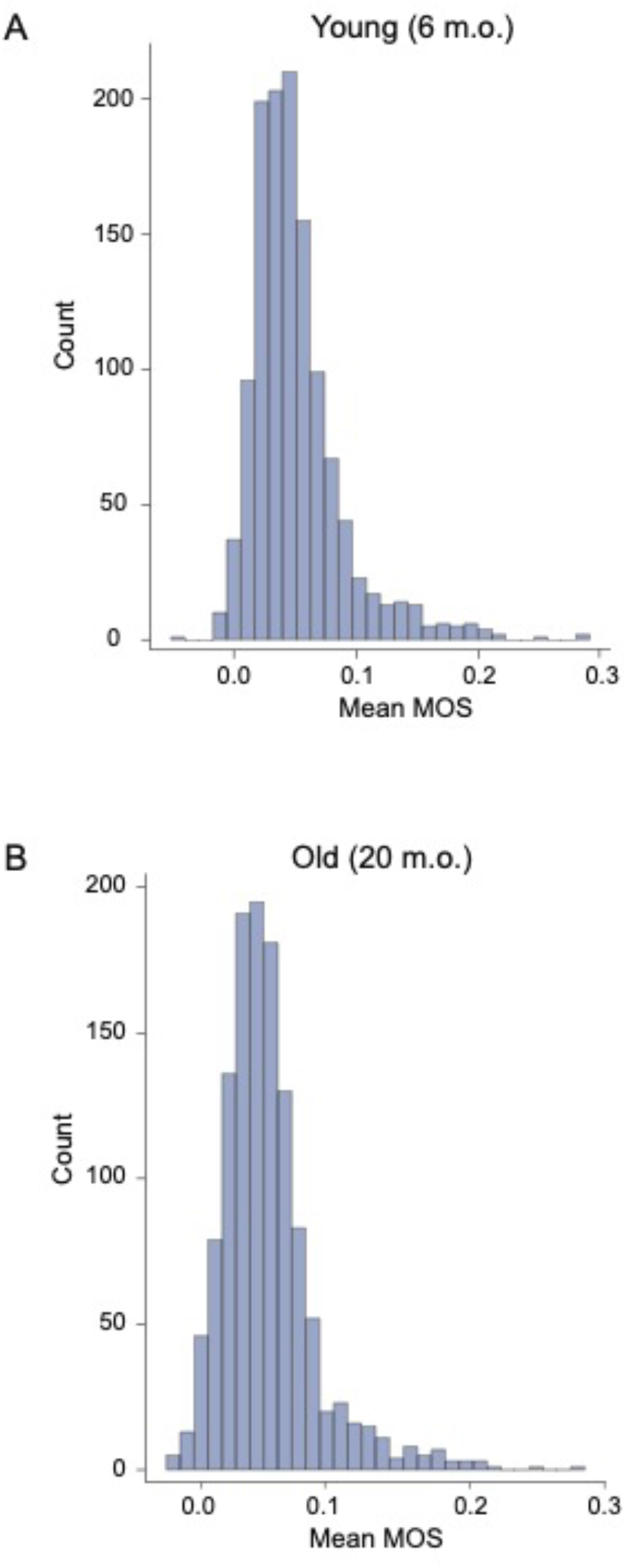
The proteome-wide distribution of measured *in vivo* methionine oxidation stoichiometries (MOS) in the brain cortices of young and old mice. Histograms showing the proteome-wide distribution of *in vivo* methionine oxidation stoichiometries (MOS) in the brain cortices of young (A) and old (B) mice. The measured average MOS values for young and old mice were 0.051 and 0.050, respectively. For both distributions, 1232 shared methionine containing peptides representing 629 proteins were quantified.

The results of a two-sample t-test comparing the proteomic distribution of *in vivo* MOS values estimated for young and old animals suggested that *in vivo* methionine oxidation does not globally increase in the brain cortices of mice during aging (p-value = 0.40). The global titration response curves used to normalize MObB data did not significantly differ between the two age groups prior to normalization, suggesting that the observed effect is not an artifact of normalization protocols (Supplementary Figure 1). Together, these results indicate that in mouse brains methionine oxidation levels are generally low and do not significantly increase as a function of age.

### MOS measurements are reproducible across biological replicates

Stochastic models of *in vivo* methionine oxidation suggest that methionine oxidation is a rare and transient event and not a regulated process like most other posttranslational modifications (PTMs). Such models suggest that “snapshot” measurements of *in vivo* methionine oxidation stoichiometries would be randomly distributed across the proteome and not biologically reproducible. However, our results demonstrate that the proteomic distribution of *in vivo* methionine oxidation stoichiometries in the brain cortices of both young and old animals are highly reproducible across biological replicates (Figure 4). Pearson correlation coefficients between different titration points of the same animal (technical variation) and Pearson correlation coefficients between the same titration point of different animals (biological variation) are approximately equal, suggesting that the highly significant correlations between biological replicates observed in this study are conservative relative to the remaining technical variation in our measurements. Furthermore, as can be seen in Supplementary Figure 2, the correlation in MOS values between biological replicates remains highly significant even when excluding imputed missing values, suggesting that the reproducibility of our measurements is not an artifact of missing value imputations.

**Figure 4.**
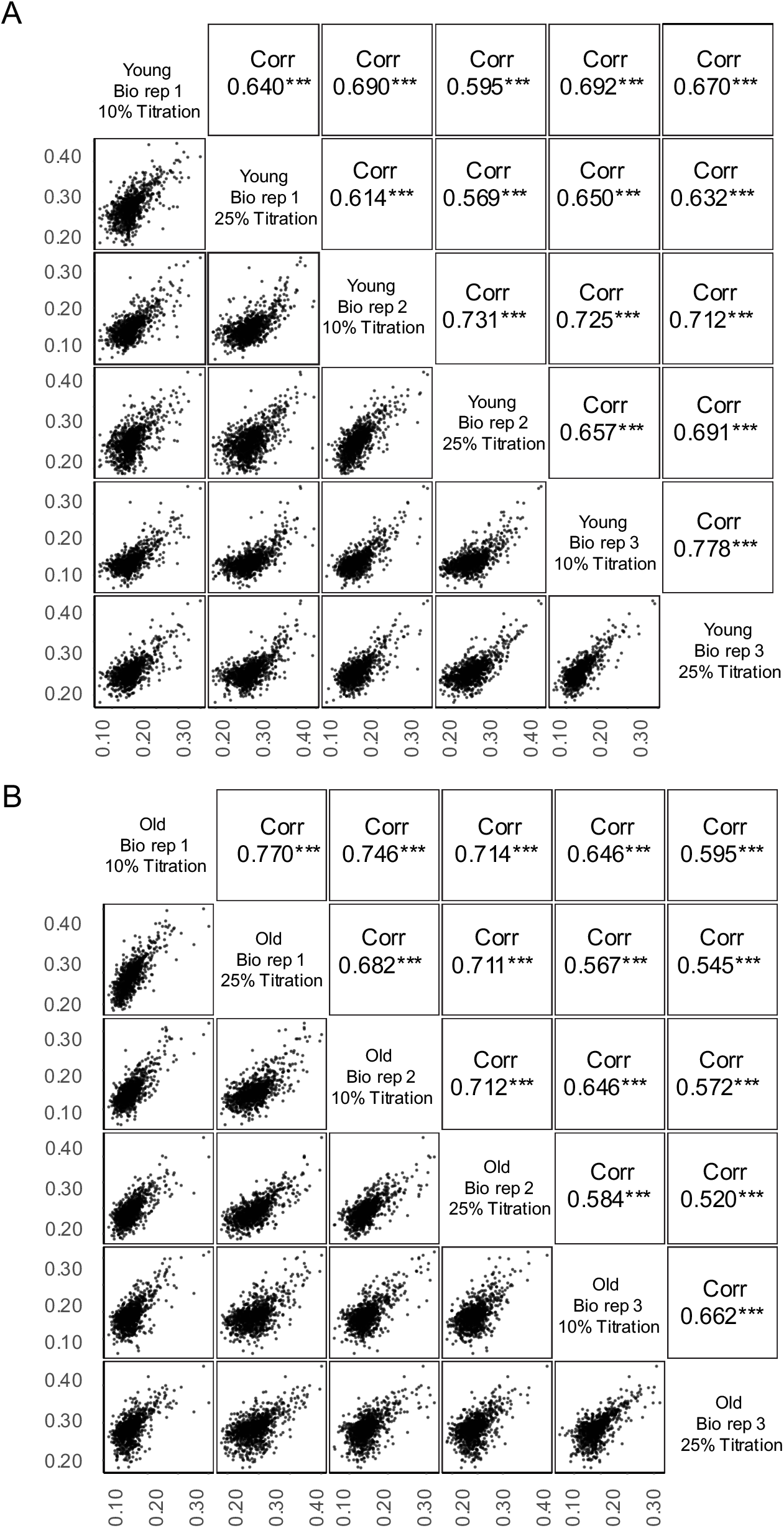
Technical and biological reproducibility of MObB measurements. Multi-scatter plot representing a complete series of pairwise comparisons of methionine oxidation stoichiometries measured between two samples from the young (A) and old (B) age groups. Pearson correlation coefficients, along with significance levels are shown in the upper panels and the data visualized in the lower panels. Each point represents a unique methionine sulfoxide containing peptide.

In addition, the Pearson correlation coefficients between biological replicates in young and old animals are approximately equal, suggesting that the proteomic distribution of *in vivo* methionine oxidation does not become more variable as mice age. These observations indicate that (i) methionine oxidation levels within biological replicates are not random, and (ii) the proteomic distribution of methionine oxidation does not become more variable as mice age. Therefore, our data suggest that oxidation levels are maintained at specific steady-state levels in both young and old animals for most methionines.

### Proteomic distributions of methionine oxidation stoichiometries remain stable during murine aging

Previously, it had been reported that the bulk tissue contents of methionine sulfoxides do not significantly change in any tissues during aging in mice.^50^ However, because these were not proteome-wide measurements, it remained unclear whether or not specific subsets of the proteome may be more vulnerable to the accumulation of methionine oxidation than others. In particular it has been suggested that proteins with slow turnover rates are the most likely to accumulate oxidation. ^26, 51–54^ In our experiments, the improved precision of MobB and the fact that *in vivo* MOS values are biologically reproducible made it possible to compare the proteomic distribution of methionine oxidation between young and old mice on a global scale.

Similar to previous bulk measurements of methionine oxidation^50^, our proteome-wide measurements also indicate that *in vivo* methionine oxidation remains stable during aging in mice. A series of two-sample t-test comparing MOS values of young and old mice suggested that there were no statistically significant differences between the two age groups, after correcting for multiple hypotheses testing (Figure 5D-E). Additionally, we did not observe a correlation between age-associated changes in MOS values and turnover rates for individual proteins (Supplementary Figure 3). Thus, neither the global content nor the proteomic distribution of methionine sulfoxides change during murine aging regardless of basal protein turnover rates.

**Figure 5.**
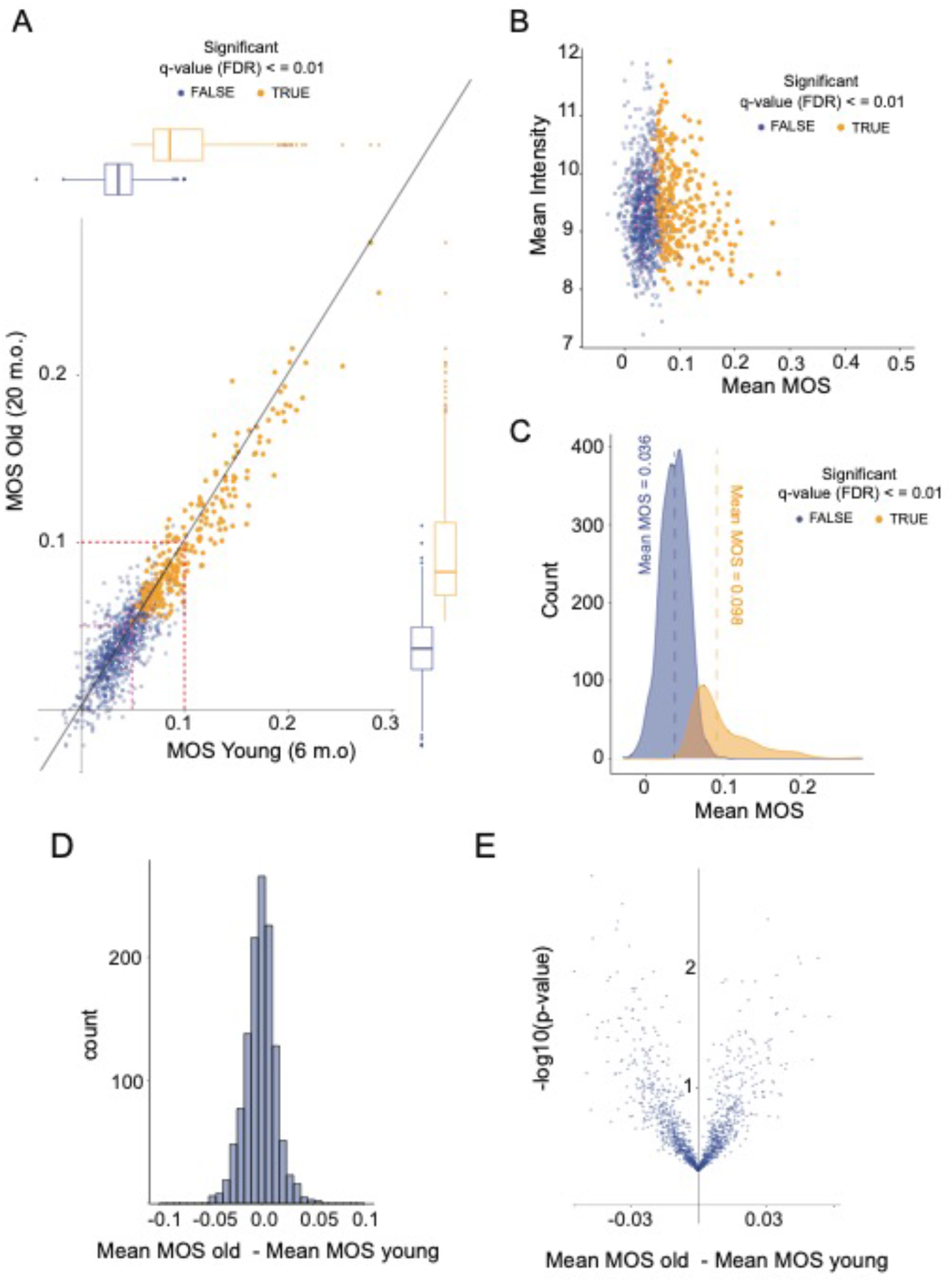
The proteomic distribution of methionine oxidation in the brain cortices of young and old mice. (A) Scatter plot comparing methionine oxidation stoichiometries measured in brain cortices of young and old mice. Each point represents a unique methionine containing peptide. Pink and red dotted lines represent the sample medians (∼5.0%) and the maximum limit of isotopic impurity in the labeling reagent (10%), respectively. (B) A scatter plot comparing the inter-age mean MOS value (x-axis) to the inter-age mean MS1 intensities (y-axis, log-scale). Each point represents a unique methionine containing peptide. (C) Density plots illustrating the global distribution of inter-age mean methionine oxidation stoichiometries. In A-C, peptides identified as having methionine oxidation stoichiometries significantly higher than the global mean are shown in orange, all other (N.S.) peptides are shown in blue. (E) Histogram showing the proteome-wide distribution of inter-age differences in mean methionine oxidation stoichiometries. (E) Volcano plot comparing the proteome-wide distribution of inter-age differences (x-axis) and their associated p-values (y-axis, -log- scale). No peptides are identified as being significantly different between age groups, after correcting for multiple hypotheses testing.

### A subset of the proteome is significantly oxidized in the brain cortexes of both young and old mice

While most methionines have MOS values that are clustered closely about a low global average (∼4.5%), there appears to be a subpopulation of the methionine-proteome that is highly and reproducibly oxidized *in vivo*, regardless of age group (Figure 5A-C). In order to identify the subset of the methionine-containing proteome that is significantly oxidized *in vivo,* we performed a statistical analysis of peptide-specific response curves for each methionine containing peptide that was quantified in our assay. Data from both age groups were analyzed simultaneously, resulting in an inter-age mean (effect size) and an associated standard error. Statistical significance was assigned by an adaptive shrinkage model (q-value <= 0.01). The resulting list of significantly oxidized methionines are highly oxidized in both age groups (Figure 5A), identified without any intensity bias (Figure 5A) and have an inter-age mean MOS value of 9.8% (Figure 5C). A complete list of methionines that are significantly oxidized *in vivo*, referred to hereafter as oxidation-prone methionines, can be found in Supplementary Table 2.

These results demonstrate that MobB is sufficient for the unbiased identification of potentially novel sites for *in vivo* methionine oxidation on a proteome-wide scale. In total, we identify over 280 potential oxidation-prone methionines in the brain corteices of young and old mice (Supplementary Table 2). Furthermore, many of the oxidation-prone methionines identified in this study have mean MOS values <30% and would have likely been inaccessible by previous proteome-wide methods used to quantify methionine oxidation, demonstrating that MobB is sufficient for the statistical evaluation of even lowly to moderately oxidized methionines.^58^

### Intrinsic protein factors do not strongly correlate with in vivo methionine oxidation stoichiometries (MOS)

We next attempted to identify specific physiochemical and biological factors that correlate with the propensity of methionines to be oxidized *in vivo* in both young and old mice. Damage-centric models of *in vivo* methionine oxidation, that describe it as a primarily nonenzymatic process, suggest that the *in vivo* stoichiometries of methionine oxidation should be strongly influenced by each individual methionine’s unique chemical environment. In particular, it has been demonstrated that there is a strong positive correlation between methionine solvent accessibility and *in vitro* oxidation propensities.^26^ Interestingly, we observe a weak but significant negative correlation between methionine solvent accessibility and *in vivo* MOS values, the opposite of what is observed in *in vitro* oxidation experiments (Figure 6A). Although an explanation for this observed trend remains to be determined, one possibility is that solvent accessibility may influence the enzymatic reduction of methionine sulfoxide residues by MSRs. Furthermore, there is no categorical association between oxidation-prone methionines (significantly oxidized methionines), and solvent accessibility (Figure 6A). Taken together, the results suggest that factors that determine *in vivo* methionine oxidation levels may be distinct from those that determine *in vitro* oxidation levels.

**Figure 6.**
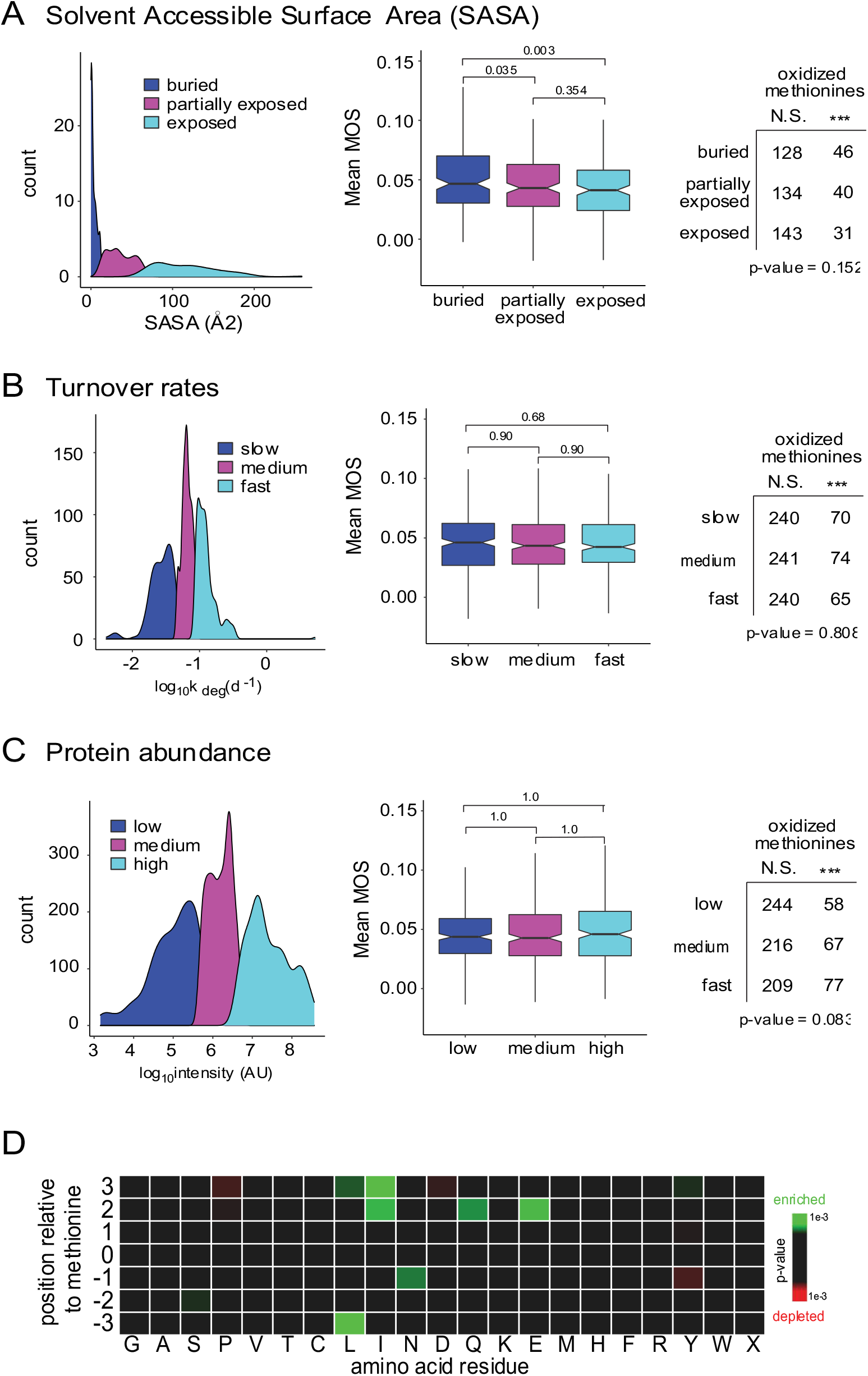
The association between *in vivo* methionine oxidation stoichiometries intrinsic protein properties. (A) Correlation between methionine solvent accessible surface areas (SASA) and oxidation stoichiometries (MOS). Density plot (left) illustrates the groupings of SASA into three categories of approximately equal sizes, buried (blue), partially exposed (magenta) and exposed (cyan). Boxplot (middle) compares methionine solvent exposure to inter-age mean methionine oxidation stoichiometries. Pairwise Wilcoxon signed-rank tests were performed, and p-values associated with the difference in means between groups are shown as bars above each plot. Table (right) compares methionine solvent exposure to the number of methionine oxidation sites identified as being highly oxidized *in vivo*. The result of a chi-squared analysis testing for an association between highly oxidized methionines and solvent exposure is shown below the table. (B) Correlation between protein turnover rates (as measured by Price et. al and Fornasiero et. al) and oxidation stoichiometries (MOS).^63, 64^ Density plot, boxplot and table are as described in (A). (C) Correlation between protein abundance (Fornasiero et. al) and oxidation stoichiometries (MOS).^63^ Density plot, boxplot and table are as described in (A). (D) Heat map illustrating positional sequence enrichments for amino acids surrounding highly oxidized methionines (target) to average methionines (background). Results are colored by p-value and only significant enrichments (p-value <= 0.01) are shown. Heat map was generated using icelogo.^68^

In addition to methionine solvent accessibility, it has been previously suggested that protein turnover may play a dominant role in the clearance of oxidized proteins.^53, 54^ The continual process of protein turnover is believed to minimize the accumulation of protein damage, including methionine oxidation. We observe, however, no evidence for a correlation between the rates of protein turnover and *in vivo* MOS values (Figure 6B). Protein turnover rates used in this study were harvested from two independent datasets, Fornasiero et. al and Price et. al, on the rates of protein turnover in mouse brain tissues.^63, 64^ Furthermore, there is no categorical association between oxidation-prone methionines and protein turnover (Figure 6B). As discussed above, there is also no correlation between the rates of protein turnover and the accumulation of *in vivo* methionine oxidation over time (Supplementary Figure 3a). Taken together, these results suggest that the basal turnover of proteins does not play a dominant role in establishing the *in vivo* steady-state between methionine and methionine sulfoxide for most proteins. It should be noted that the rates of protein turnover used in this study were quantified using ensemble methods that do not distinguish between oxidized and unoxidized proteoforms.^63, 64^ We therefore cannot rule out the possibility that degradation pathways dedicated to the clearance of oxidized proteins, such as the 20S proteosome, may play an important role in establishing the *in vivo* steady-state levels of methionine sulfoxide.

Protein abundance and amino acid sequence were also analyzed for possible correlations with *in vivo* MOS values. As can be seen in Figure 6C, we were not able to identify any significant correlations between *in vivo* MOS values and protein abundance, as measured by Fornasiero et. al.^63^ A heatmap map comparing the surrounding sequence of oxidation-prone methionines to the sequence surrounding average methionines suggests that there are some weak enrichments for titratable (N, Q, E) and aliphatic (I, L) amino acids surrounding oxidation-prone methionines (Figure 6D). However, there is no clear consensus sequence that is predictive of methionine oxidation.

### In vivo methionine oxidation stoichiometries (MOS) are enriched for clusters of functionally related gene ontology (GO) terms

Taken together, the above results suggest that the intrinsic physiochemical properties of proteins and methionines are not strong determinants of *in vivo* methionine oxidation. We next investigated the influence of external biological factors on the proteomic distribution of *in vivo* methionine oxidation. To this end, we conducted a gene ontology (GO) enrichment analysis on oxidation-prone methionines. As can be seen in Figure 7 highly oxidized methionines are enriched for clusters of functionally related GO terms. Biological pathways enriched for oxidized methionines include those related to (i) small molecule metabolism and ATP generation, (ii) homeostasis and the regulation of biological quality, and (iii) chemical and vesicle mediated synaptic transmission (Figure 7A, Supplementary Table 3). However, the strongest enrichments we observe are those for subcellular localization including terms related to (i) the extracellular space, (ii) the plasma membrane/cell periphery and (iii) membrane-bounded vesicles (Figure 7B, Supplementary Table 3).

**Figure 7.**
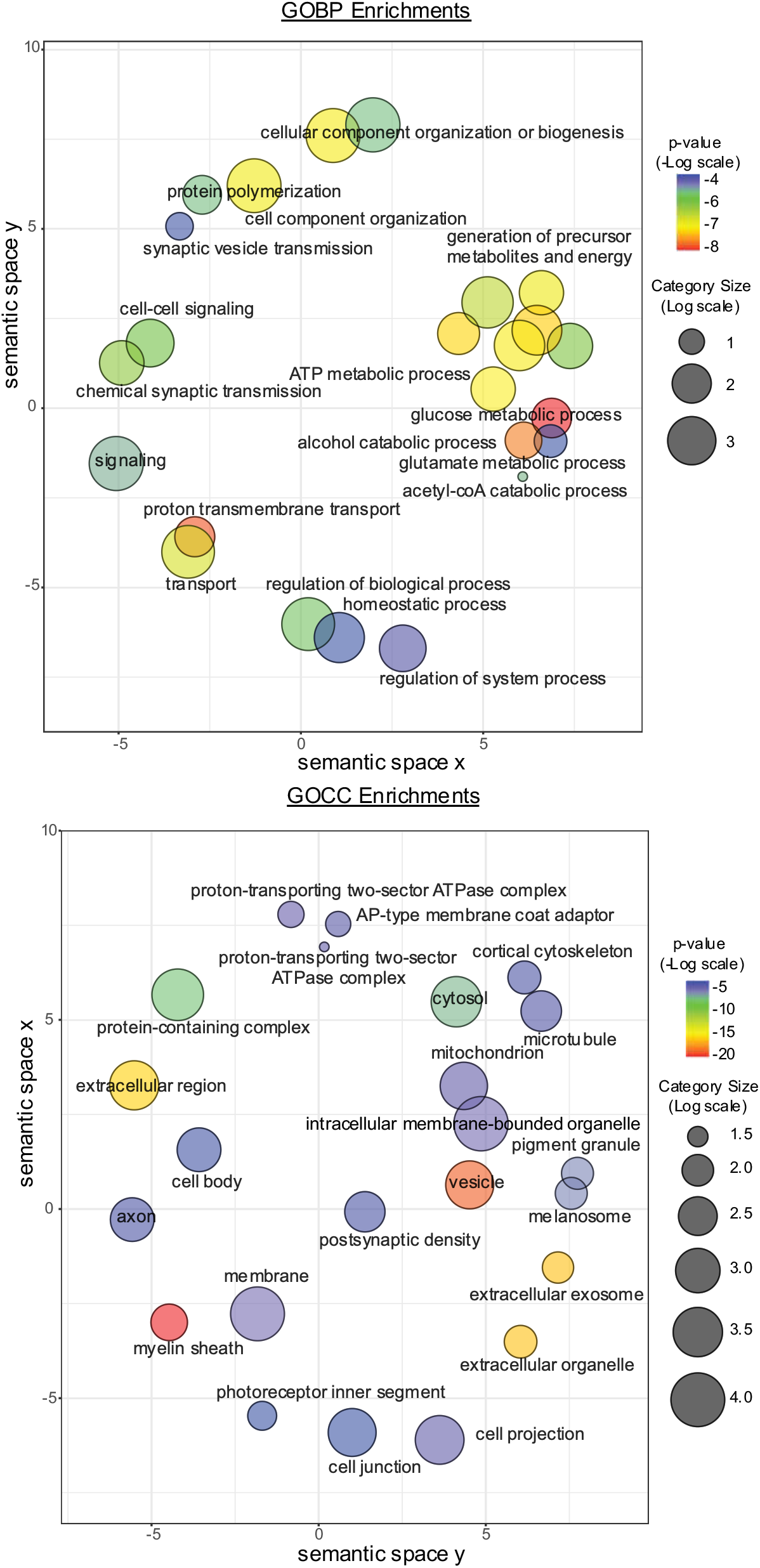
Gene ontology (GO) categories enriched for methionine oxidation in mouse brain cortices. Disc color indicates the Benjamini-Hochberg corrected p-value for enrichment in the set of peptides identified as having significant amounts of *in vivo* methionine oxidation. Size is proportional to log number of total genes in category. Enrichment is significant at *p-adjusted <= 0.05* (Fisher’s exact test). Spatial arrangement of discs approximately reflects a grouping of categories by semantic similarity. Visualized categories have been selected from a broader set (Supplementary Table 3) to eliminate redundancy and obsolete terms. Visualization was done using the REViGO tool available at http://revigo.irb.hr/.

It is worth noting that the observed enrichments for subcellular localization of oxidation-prone methionines strongly overlaps with the known role and localization of MICAL proteins, enzymatic writers of methionine oxidation.^10, 11^ For example, it has been well documented that the monooxygenase domains of MICALs are important to the biogenesis and trafficking of exocytic vesicles; as well as their fusion to the plasma membrane.^16, 18, 65^ We report here that vesicles, extracellular exosomes, the plasma membrane and cortical cytoskeleton are all enriched for oxidation-prone methionines, suggesting that the intersection between exocytotic pathways and *in vivo* methionine oxidation may be more widespread than previously appreciated. Furthermore, MICAL proteins contain a conserved LIM domain that results in their localization to the cortical cytoskeleton *in vivo*.^66^ We report here that not only is the cortical cytoskeleton enriched for oxidation prone methionines, but so are other components of the cell periphery including the plasma membrane, post-synaptic density, myelin-sheath, cell junctions and cell projections.

Taken together, these results suggest that extrinsic, biological factors are the strongest determinants of *in vivo* methionine oxidation, with subcellular localization playing a dominant role. The extracellular space, cell periphery and membrane-bounded vesicles are all strongly enriched for oxidation-prone methionines, consistent with the hypothesis that the *in vivo* localization and activity of MICAL proteins play a significant role in establishing the proteomic distribution of *in vivo* methionine oxidation. Although outside the scope of this study, future experiments that combine the use of MObB and reverse genetics can be used to directly measure the role of MICALs in establishing *in vivo* methionine oxidation levels under varying physiological conditions.

## Discussion

We report an updated proteomic workflow that allows for the precise and unbiased quantification of *in vivo* methionine oxidation stoichiometries (MOS). Our workflow is based on a previously described stable isotope labeling strategy that prevents the *in vitro* accumulation of methionine oxidation by converting all *in vivo* unoxidized methionines to heavy labeled methionine sulfoxides, near the time of cell lysis.^57, 60, 61^ We have demonstrated that our workflow, which we have named Methionine Oxidation by Blocking (MObB), is sufficient for the quantitative and statistical analysis of *in vivo* methionine oxidation stoichiometries even when present at low levels (<5%).

We have applied MObB to generate a quantitative description of the proteomic distribution of *in vivo* methionine oxidation in the brain cortices of young (6 m.o.) and old (20 m.o.) mice. We find that *in vivo* MOS values are generally low in both age groups (Figure 3). In addition, we find no evidence for significant age-dependent effects on the proteomic distribution of *in vivo* MOS values (Figure 5d, e). Our results agree with previous observations that the global content of methionine sulfoxide does not significantly increase in mouse tissues during murine aging.^50^ Our results build upon this observation by demonstrating that not only does the global content of methionine sulfoxide not significantly change during murine aging, but neither does the proteomic distribution of methionine oxidation. Furthermore, we find no evidence for a significant relationship between protein turnover kinetics and the accumulation of methionine oxidation during murine aging.

In addition, we demonstrate that, for a subset of the methionine-containing proteome, methionine and methionine sulfoxide exist in a quantifiable steady-state that is biologically reproducible; we have named these methionines oxidation-prone methionines (Figure 3, Figure 5A-C, Supplementary Table 2). As discussed above, the steady-state of oxidation-prone methionines are not only biologically reproducible among individuals, but they do not change significantly as a function of age. Our results are consistent with the notion that *in vivo* methionine oxidation is a regulated process in a manner that is similar to other posttranslational modification (PTMs).

The mechanisms by which organisms are able to maintain steady-state levels of methionine sulfoxides remain to be determined. However, our results provide some preliminary suggestions that hint at plausible mechanisms. For example, we observe no strong relationship between the intrinsic or physiochemical properties of individual methionines and their *in vivo* propensity to be oxidized (Figure 6). This suggests that passive mechanisms in which the *in vivo* steady state between methionine and methionine sulfoxide is driven solely by chemical determinants and intrinsic properties of the protein are unlikely. Our data instead support the notion that extrinsic or biological effects play a dominant role in establishing the *in vivo* levels of methionine oxidation for many proteins, with subcellular localization being the most significant factor (Figure 7, Supplementary Table 3).

Sequestering reactive metabolites from reactive amino acid sidechains is one of the primary mechanisms that cells employ to regulate nonenzymatic posttranslational modifications and subcellular localization has been previously described as a primary determinant of methionine oxidation under conditions of oxidative stress.^21, 30^ However, the specific gene ontology (GO) enrichments we observe for *in vivo* methionine oxidation in mice under unstressed conditions suggests a more nuanced mechanism. *In vivo* oxidation-prone methionines in the brain cortices of unstressed mice are enriched for terms related to (i) the extracellular space, (ii) the cell periphery and (iii) membrane bounded vesicles (Figure 7, Supplementary Table 3). These clusters of functionally related GO terms possibly hint at an important role for MICAL activity in the proteomic distribution of methionine oxidation in mouse brain tissues. For example, all three MICAL genes in the *Mus musculus* genome contain a conserved LIM domain, which localizes MICAL proteins to the cortical cytoskeleton.^66^ As discussed above, we not only see enrichments for methionine oxidation at the cortical cytoskeleton but, more generally, a cluster of enrichments for methionine oxidation at the cell periphery (Supplementary Table 3). In addition, MICAL activity has been shown to be important for the biogenesis and trafficking of secretory vesicles.^16, 18, 65^ We observe not only enrichments for terms related to the biogenesis of synaptic vesicles, but also strong enrichments for both cytoplasmic and extracellular vesicles (Figure 7, Supplementary Table 3). It should be noted that, within the *Mus musculus* genome, MICAL1 and MICAL3 contain a Rab binding domain (RBD) that allows them to interact with Rab-GTPases on vesicles surfaces.^66^ In this current study we observe significant methionine oxidation of several Rab proteins on vesicle surfaces (Supplementary Table 4).

Although the above observations suggest that the localization and function of MICAL proteins may play broader role in the proteomic distribution of oxidation-prone methionines than previously appreciated, this hypothesis remains to be tested directly. Future proteomic experiments that employ the methodologies described in this study to measure methionine oxidation levels in MICAL deficient mice will greatly expand our understanding of the role of MICALs in establishing *in vivo* methionine oxidation stoichiometries.

## Acknowledgement

We thank the members of the Ghaemmaghami lab at the University of Rochester for helpful discussions and suggestions.

## Data Availability

All raw and processed data are available in the included Supporting Information and at the ProteomeXchange Consortium via the PRIDE partner repository (accession number PXD030245).^67^ Currently, the data can be accessed with the username and password mEYyupdO.

## Author Information

### Corresponding Author

*. Phone 585-275-4829

## Author Contributions

The study concept was conceived by J.B. and S.G. Its detailed planning was performed with contribution from all authors. M.S. and A.K. assisted with animal handling, euthanasia and tissue extraction. All wet-lab experiments were conducted by J.B. Mass spectrometric analyses were conducted by K.W., J.H. and J.B. Data analysis was conducted by J.B. The manuscript was written by J.B. and S.G. All authors have given approval to the final version of the manuscript.

## Notes

The authors declare no competing financial interest.

## Supporting information

Supplementary information includes:

- Supplementary Figures 1-6
- Supplementary Materials and Methods
- Supplementary Results
- Supplementary Table 1 (Supp_Table1.xls) contains a quantitation of MOS values across all samples.
- Supplementary Table 2 (Supp_Table2.xls) contains the results of a statistical analysis used to identify oxidation-prone methionines.
- Supplementary Table 3 (Supp_Table3.xls) contains the results of a GOterm analysis on oxidation-prone methionines.
- Supplementary Table 4 (Supp_Table3.xls) contains GOterm annotations of oxidation-prone methionines.

## Abbreviations

ROS: reactive oxygen species
PTM: posttranslational modification
MICAL: ‘molecule interacting with CasL’ protein
BCA: bicinchoninic acid assay
HPLC: high performance liquid chromatography
RT: retention time
NRMSE: normalized root mean square error
SEM: standard error of mean
FDR: false discovery rate
MOS: methionine oxidation stoichiometry
SASA: solvent accessible surface area
GO: gene ontology

