## Supplementary Materials and Methods for "Accurate proteome-wide measurement of methionine oxidation in aging mouse brains"

##### *Feature Selection*

Due to the previously reported observation that $H_2O_2$ is efficient at labeling both cysteine and methionine residues, the data analyzed here is restricted to tryptic peptides that contain one methionine and no cysteines.[1] In addition, in order to limit the computational complexity of the search to be conducted, MS1 features are limited to only those that can be matched between technical replicates. MS1 features are matched based on retention time, mass to charge ratio and fraction.

For each MS1 peptide feature that was identified by MaxQuant, the associated retention time range identified by MaxQuant was used to subset a swath of the MS1 spectra predicted to contain the associated feature. The swath was then filtered for peaks with a centroided mass to charge ratio within a 10 ppm window centered on the predicted mass to charge ratios for the identified peptide feature. Predicted mass to charge ratios were restricted to five isotopologues, ranging from the monoisotopic $^{16}O$ (light)-labeled peak to the +2 $^{18}O$ (heavy)-labeled peak, for each peptide.

##### *Chromatographic peak modeling and isotope cluster assembly.*

A chromatographic peak model was then fit to each isotopologue ion chromatogram. Chromatographic peaks were modeled as asymmetric Gaussians, with retention time (RT) as the independent variable (x-axis) and intensity as the dependent variable (y-axis). A maximum of two asymmetric Gaussian peaks were allowed for each isotopologue. The MObB algorithm choses between one peak and two peak models based on the agreement (Pearson correlation) between predicted and RAW intensities.

Modeled peaks, representing isotopologues consecutive along the *m/z* axis, are only connected if the cosine similarity between them is equal to or greater than 0.6. For isotopologues represented by a mixture of two asymmetric gaussians, each gaussian is represented as a separate peak and all possible combinations of cosine similarities with the neighboring isotopologue are assayed. Asymmetric Gaussian peaks that cannot be connected to any peak in the neighboring isotopologue are filtered out of the final model.

The MObB algorithm fails to assemble a model for a portion of the MS1 features returned by MaxQuant. The MObB algorithm will reject any model it cannot assemble a complete isotope cluster for and will reject any isotopologue it cannot model as an asymmetric Gaussian or mixture of asymmetric Gaussians.

*Linear modeling for the estimation of L/(L+H) ratios, or MOS values.*

Following modeling and assembly of all MS1 features identified in a given sample, MS1 features were grouped by peptide sequences and assembled into one ‘super’-feature, that is a combination of all non-redundant MS1 data points representing a peptide sequence. Peptide sequences with missed cleavages are grouped with their properly cleaved sequence counterpart; provided one could be identified.

For each ‘super’-feature, intensities are deisotoped into ($^{16}O$) light and ($^{18}O$) heavy channels. Deisotoping is done on a scan-by-scan basis resulting in a retention-time array of intensities for $^{16}O$ (light)-labeled peptides and matched intensities for $^{18}O$ (heavy)-labeled peptides.

A linear model is used to estimate an average ratio, or slope, between $^{16}O$ (light)-labeled peptides and $^{18}O$ (heavy)-labeled peptides. The y-intercept in the linear models used is not forced to zero, under the assumption that light and heavy channels may have different baselines or noise levels.

The slope of each linear model is a peptide-specific L/H ratio. However due to the theoretical overlap in the isotopic envelopes of ($^{16}O$) light labeled-peptides and ($^{18}O$) heavy-labeled peptides a correction factor is applied to the experimentally observed L/H ratios. For each peptide sequence, a theoretical isotope cluster was generated using the atomic composition of the peptide and the known abundance of natural isotopes. Theoretical models were generated for each peptide with theoretical ratios between the light ($^{16}O$) labeled-peptides and ($^{18}O$) heavy-labeled peptides ranging between 0 and 1 with a step size of 0.01. Theoretical isotope models were then deisotoped into light and heavy channels. A peptide-specific calibration curve was generated by comparing the ratio of theoretical intensities in light and heavy channels to the theoretical ratio in light ($^{16}O$) labeled-peptides and ($^{18}O$) heavy-labeled peptides used to generate them. As a final step, calibrated L/H ratios are converted to L/(L+H) ratios, or MOS values.

In order to pass quality filters, the linear regression used to estimate MOS values must have a coefficient of determination greater than or equal to 0.8.

*Missing value imputations.*

Three algorithms were evaluated for their efficacy in imputing missing values for MObB generated data, KNNimpute, LLSimpute and SVDimpute.[2, 3] For a complete description of each algorithm see refs 2-3. The efficacy of each algorithm was evaluated by measuring a Pearson correlation between the known value of artificially generated missing values to the values imputed by each algorithm. In total, 30% of the originally known data were temporarily converted into artificial missing values. Only a small subset of the peptides (rows) had no missing values in any sample, therefore artificial missing values were straited for peptides (rows) with missing value densities that approximates the missing value density of the experimental array. For example, if 10% of the experimental array had a one missing value, then 10% of the artificial missing values were sampled from rows with one natural missing value. In the case of SVDimpute missing values were first converted to zeros, and then imputed using an expectation maximum (EM) method, as previously described.

KNNimpute was found to be the best performing and most stable algorithm for imputing missing values in the current study. For final imputations, fifteen similar peptides were used to impute missing values in the KNNimpute algorithm. All other options were set to default values. Missing value imputation was performed using an in-house R script, and is available as supplementary file.

*Data normalization.*

Normalization was done in two steps. First, a global titration response curve was generated using data from young and old animals separately. Titration response curves were generated using the following equation,

$$MOS_i = (1 - t_i)(MOS_{in\ vivo}) + (t_i)$$

where $MOS_i$ is the global median of measured L/(L+H) ratios in titration sample *i*, $MOS_{in\ vivo}$ is the global median of estimated MOS values that would be measured without any carrier proteome and $t_i$ is the relative ratio of the carrier proteome used to create sample *i*.

Global titration response curves from young and old animals did not significantly differ from one another (Supplementary Figure 1). Data from young and old animals were therefore grouped together to create an experiment-wide titration response curve (Supplementary Figure 1). The experiment-wide response curve was then used to normalize the data. A normalization factor was calculated for each quantitation channel by adjusting the global medians of each channel to agree with the predicted values returned by the experiment-wide response curve. Following normalization of all quantitation channels data from technical replicates were combined by taking the median of both samples.

*Peptide-specific titration response curves and estimating in vivo MOS values.*

Peptide-specific titration response curves were generated as described above. Titration response curves were generated using the following equation,

$$MOS_{ij} = (1 - t_i)\left(MOS_{in\ vivo,j}\right) + (t_i)$$

where $MOS_{ij}$ is the measured L/(L+H) ratio for peptide *j* in sample *i*, $MOS_{in\ vivo,j}$ is the estimated *in vivo* MOS value that would be measured for peptide *j* without any carrier proteome and $t_i$ is the relative ratio of the carrier proteome used to create sample *i*. The nonlinear regression algorithm used to fit titration response curves returns an estimated parameter for $MOS_{in\ vivo,j}$ , as well as an associated standard error of means (SEM). Peptide-specific titration curves were generated using either age-specific grouping of data or no age-specific grouping of data (inter-age), where indicated in text.

In order to pass quality filters peptide-specific response curves must have fit the data with a normalized root mean-squared error (NRMSE) of less than or equal to 0.2.

#### Supplementary Results

*Chromatographic peak modeling improves the quality of $^{16}O/^{18}O$-methionine labeled isotope clusters.*

Previous attempts at quantifying in-vivo methionine oxidation suggested that more precision was needed to quantify the naturally low abundance of in-vivo methionine sulfoxide in unstressed cells.[4] Manual inspection of RAW data suggested that instrumental noise and interference from neighboring peptides was a major source of analytical variation. In order to address this issue, we have developed the computational tools necessary to improve the assembly of $^{16}O/^{18}O$-methionine labeled isotope clusters (Figure 1b). Improved modeling of chromatographic peaks allows MObB to filter noise and disentangle the signal from closely spaced peptides, provided that they do not have identical elution times. However, one of the challenges in modeling the elution behavior of methionine sulfoxide containing peptides is the observation that methionine sulfoxide containing peptides may or may not have stereospecific elution times.[5]

As can be seen in Supplementary Figure 4 the novel chromatographic peak modeling functions of MObB (orange) have high agreement with RAW data (blue), while still allowing for the smoothening of instrumental noise or ‘wobble’ in the RAW data. Chromatographic peaks were modeled as asymmetric gaussians and MObB’s core computational pipeline accurately and automatically chooses between one-peak (non-stereospecific, Supplementary Figure 4a) and two-peak (stereospecific, Supplementary Figure 4b) models.

Furthermore, as can be seen in Supplementary Figure 4c the chromatographic peak modeling functions of MObB significantly improve the quality of $^{16}O/^{18}O$-methionine labeled isotope clusters, where the quality of isotope clusters is measured as the cosine similarity between the elution profiles of light and heavy labeled isotopologues. In addition, a subset of low-quality MS1 peptide features are rejected by MObB's chromatographic peak modeling functions, on the basis that no gaussian models could be fit to the RAW data. Therefore, the chromatographic peak modeling functions of MObB allow for the filtering of noise/interference to improve signal quality and also acts as a quality filter for signals that could not be improved.

*Linear models allow for the robust estimation of L/(L+H) ratios.*

Previous attempts at quantifying in-vivo methionine oxidation estimated L/(L+H) ratios using a model-dependent strategy that relied on the sum of intensities across the retention-time axis.[1] The resulting 2D isotope clusters were than compared to an array of theoretical isotope clusters, with varying L/(L+H) ratios, and the L/(L+H) ratio of the best matching theoretical isotope cluster was reported as the MOS value. This strategy, however, was sensitive to experimental stochasticity and did not offer the quantitative resolution needed for highly precise measurements.

In this current study we improve upon this strategy by using a linear model to quantify an average scan-by-scan L/(L+H) ratio. In such a model, for each scan, the sum of intensities for light and heavy labeled isotopologues are plotted against one another and a linear regression is used to measure a slope, or average L/H ratio (Supplementary

Figure 5a). Measured L/H ratios are then corrected by the theoretical overlap between light and heavy labeled peptide signals before being converted to a L/(L+H) ratio.

As can be seen in Supplementary Figure 5a, the use of linear models for estimating L/(L+H) ratios is robust against nonrandom error. In addition, the use of linear models is sufficient for the quantification of site-specific differences in L/(L+H) ratios, within a single sample. Although for most methionine sulfoxide containing peptides, we are able to measure a highly accurate linear model, for a subset of peptides we are not able to measure an accurate linear model ($R^2 < 0.8$). The signal from these peptides is presumably dominated by noise and are therefore rejected by our quality filters (Supplementary Figure 5b). The use of linear models for estimating L/(L+H) ratios is not only robust against experimental stochasticity, but also an effective means for filtering low quality data.

It should be noted that although this strategy is robust against experimental stochasticity, the theoretical overlap between light and heavy labeled peptides is still calculated using a deterministic model of isotope distributions, and future improvements to MObB may include previously defined stochastic models of isotope distributions.[6, 7]

*Theoretical titration response curves allow for the estimation of in-vivo methionine oxidation stoichiometries.*

It has been previously reported that the in-vivo abundance of methionine sulfoxide (light labeled peptides) in unstressed cells is, on average, below or near the limit of quantitation (LOQ).[1] We address this issue by taking advantage of 'carrier proteome' strategies recently developed for the analysis of low-intensity samples such as single-cell proteomic samples or unenriched posttranslational modifications.[8] In this strategy a carrier proteome is made to mimic experimental samples and is spiked-in to the background of low-intensity samples in order to ensure peptide identification and quantification.

Conveniently, a similar strategy was applied to validate the initial development of labeling protocols for the $^{18}$O-blocking methodology, described in **ref 3**, of which MObB is an updated extension of.[1] Briefly, carrier proteomes made to mimic proteomes with complete in-vivo (light ($^{16}$O)-labeled) methionine oxidation were spiked into the background of heavy ($^{18}$O)-labeled proteomes and the resulting ratio between light and heavy labeled peptides was quantified as a function of the known mixing ratios. The results of this validation strategy were largely successful on both a global and site-specific scale.

Herein, we take advantage of this observation and utilize theoretical titration response curves to extrapolate site-specific in-vivo MOS values (Supplementary Figure 5c). For each peptide an MOS value is measured for an array of titration samples, prepared by mixing light labeled carrier proteomes with endogenous, heavy labeled proteomes at varying, prespecified ratios (10% and 25%). The resulting spectra has light labeled carrier proteome intensities lying on top of endogenous (in-vivo) light labeled proteome intensities. Endogenous and carrier proteome intensities are disentangled by adding varying amounts of carrier proteome and measuring a titration response. Theoretical titration response curves are fit to the data as described in Materials and Methods. The y-intercept of the resulting theoretical response curves, or the MOS value we expect to measure without any carrier proteome, is the estimated in-vivo MOS value.

As can be seen in Supplementary Figure 5c, the use of carrier proteomes and theoretical titration response curves is sufficient for quantifying site-specific in-vivo MOS values. Furthermore, for most methionines, we are able to measure titration responses that closely approximate the theoretical response (Supplementary Figure 5d). For a subset of the methionine-proteome we are not able to measure titration responses that agree with what is theoretically expected. This subset of peptides are not quantitatively diagnostic and are therefore removed from the assay by our quality filters (NRMSE > 0.2). The use of carrier proteomes and theoretical titration response curves is therefore not only an effective means for boosting the signal of lowly abundant peptides above the LOQ, but also an effective means for filtering low quality data.

*Missing value imputation improves coverage*

MobB is based upon a data dependent acquisition (DDA) acquisition strategy and is accordingly prone to missing values when combining datasets from different samples. Missing value densities for the current study range between 11-30% with an average of 16% (Supplementary figure 6a). In order to improve final coverage, a typical MobB workflow includes novel missing value imputation protocols.

Three different algorithms commonly used for the imputation of missing values were evaluated in the current study, KNNimpute, LLSimpute and SVD impute.[2, 3] KNNimpute imputes missing values by selecting *k* peptides with similar MOS values across other samples in the experiment, and then calculates a weighted mean to estimate missing values. For example, if we consider peptide A with a missing value in sample 1, KNNimpute will select *k* peptides with similar MOS values in *N*-2 other samples (where

N is the total number of samples in the experiment) and then calculate a weighted mean of MOS values in experiment 1 for the subset of *k* similar peptides, weighted by similarity to peptide A. Similarity is measured using k-nearest neighbors.[2]

Similarly, LLSimpute will impute missing values by using a linear combination of *k* peptides with similar MOS values across other samples in the experiment. In the case of LLSimpute, similarity is calculated by Pearson correlation.[3] SVDimpute differs from both KNNimpute and LLSimpute by first decomposing the experimental array into a set of eigenvectors and then missing values are imputed by a linear combination of *k* most significant eigenvectors.[2] For a more complete description of the algorithms used for missing value imputations in this current study see **refs7-8**.

As can be seen in Supplementary Figure 6b, KNNimpute is the best performing algorithm for the imputation of missing values in MObB generated data, for this current study. Performance was evaluated by artificially masking experimentally measured values as missing values. The performance of each algorithm is defined as the Pearson correlation coefficient between imputed values for artificially masked missing values and their original, experimentally measured values. The number of *k* similar peptides (KNNimpute, LLSimpute) or k most significant eigenvectors (SVDimpute) was arrayed for values ranging between 1-20. Not only is KNNimpute the best performing, but it is also the most stable towards parameter selection. Missing values in the current study were imputed using the KNNimpute algorithm with an optimal parameter selection of k=15.

A

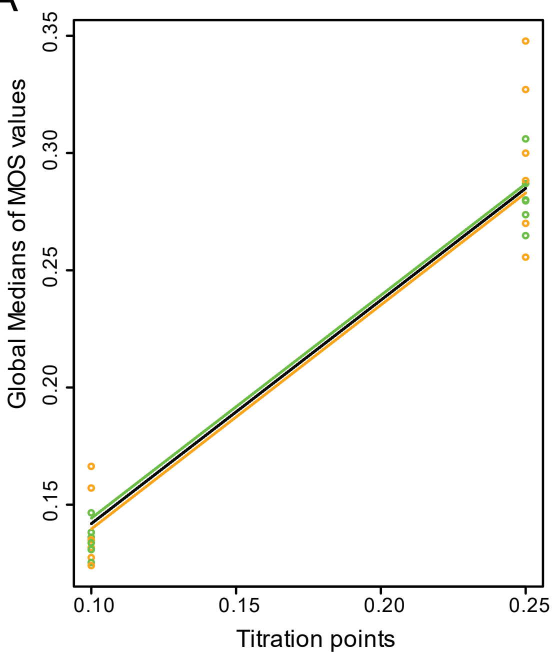


| | Young (6 m.o.) | Old (20 m.o.) |
| --- | --- | --- |
| MOS | 0.045 | 0.038 |
| SD | 0.010 | 0.010 |
| SE | 0.004 | 0.004 |
| N | 6 | 6 |

p-value = 0.316
N.S.

B

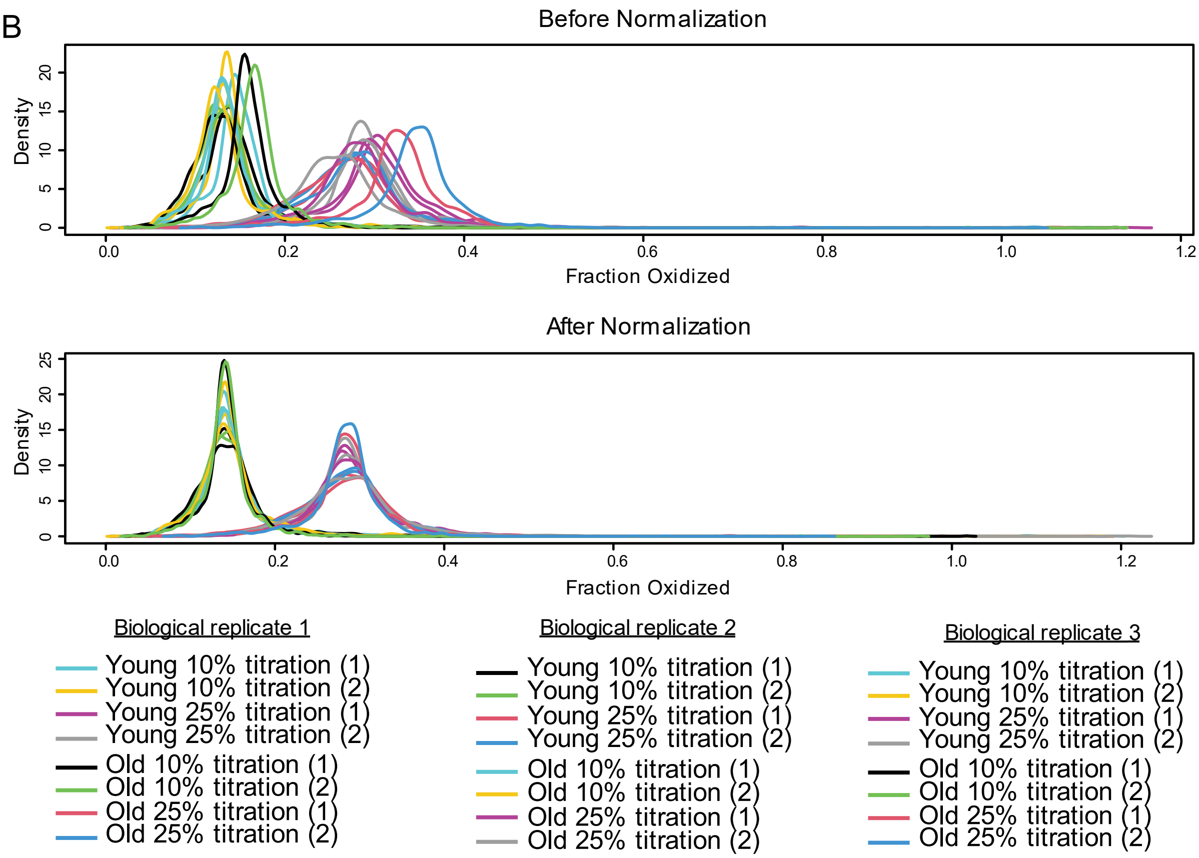

**Supplementary Figure 1 Normalization strategy used for MobB data**. (A) Global titration response curves for young (green) and old (orange) samples. On the x-axis is the titration value used for each sample and on y-axis is the global median of methionine oxidation stoichiometries measured in each sample. The titration responses of young (green) and old (orange) proteomes are not significantly different from one another (right). The black line represents the experiment-wide reference titration response curve used for data normalization. (B) Density plots of measured methionine oxidation stoichiometries in each sample before (top) and after (bottom) normalization.

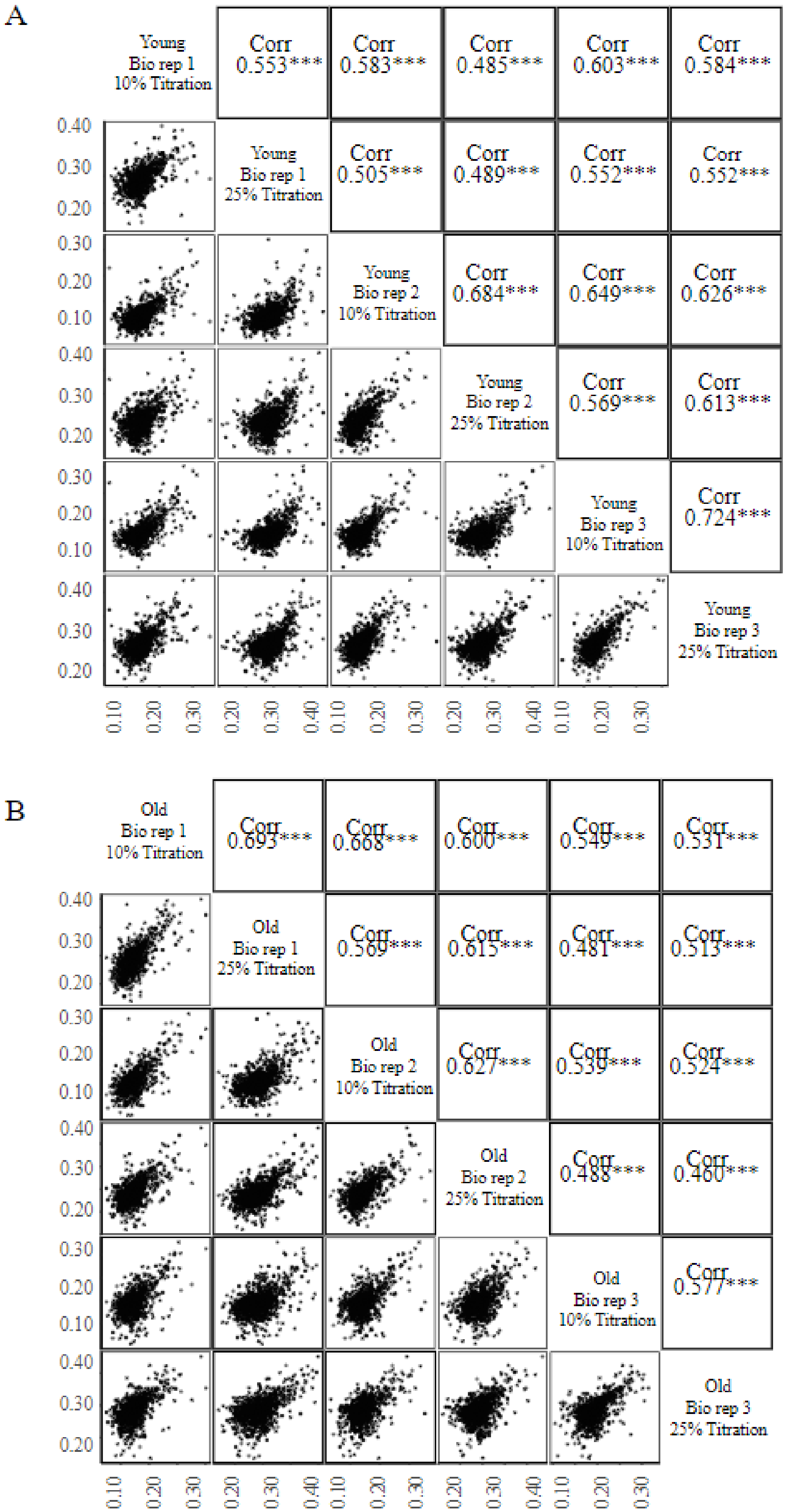
A
Young Bio rep 1 10% Titration
Corr 0.553***
Corr 0.583***
Corr 0.485***
Corr 0.603***
Corr 0.584***
Young Bio rep 1 25% Titration
Corr 0.505***
Corr 0.489***
Corr 0.552***
Corr 0.552***
Young Bio rep 2 10% Titration
Corr 0.684***
Corr 0.649***
Corr 0.626***
Young Bio rep 2 25% Titration
Corr 0.569***
Corr 0.613***
Young Bio rep 3 10% Titration
Corr 0.724***
Young Bio rep 3 25% Titration
B
Old Bio rep 1 10% Titration
Corr 0.693***
Corr 0.668***
Corr 0.600***
Corr 0.549***
Corr 0.531***
Old Bio rep 1 25% Titration
Corr 0.569***
Corr 0.615***
Corr 0.481***
Corr 0.513***
Old Bio rep 2 10% Titration
Corr 0.627***
Corr 0.539***
Corr 0.524***
Old Bio rep 2 25% Titration
Corr 0.488***
Corr 0.460***
Old Bio rep 3 10% Titration
Corr 0.577***
Old Bio rep 3 25% Titration

**Supplementary Figure 2. Quality control plots for all data collected without missing value imputation**. (A) A mulitscatter plot representing a complete series of pairwise comparisons of methionine oxidation stoichiometries measured between two samples from the young age group. Pearson correlation coefficients, along with significance levels are shown in the upper panels and the data visualized in the lower panels. Each point represents a unique methionine sulfoxide containing peptide. (B) A mulitscatter plot representing a complete series of pairwise comparisons of methionine oxidation stoichiometries measured between two samples from the old age group. Pearson correlation coefficients, along with significance levels are shown in the upper panels and the data visualized in the lower panels. Each point represents a unique methionine sulfoxide containing peptide.

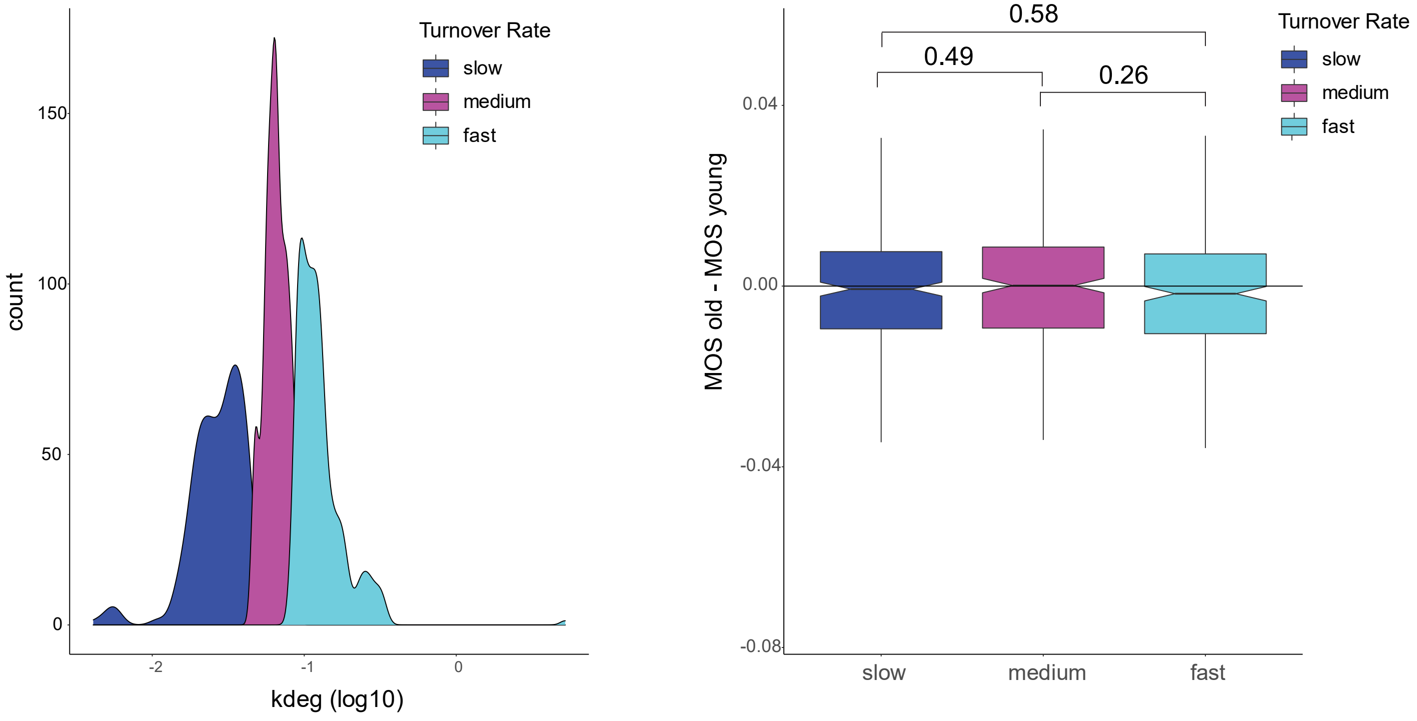

Turnover Rate
slow
medium
fast
count
150
100
50
0
-2
-1
0
kdeg (log10)
0.58
0.49
0.26
0.04
0.00
-0.04
-0.08
MOS old - MOS young
slow
medium
fast
Turnover Rate
slow
medium
fast

**Supplementary Figure 3. Protein turnover and the accumulation of methionine oxidation during mammalian aging**. (A) A density plot illustrating how protein turnover rates were grouped into three categories of approximately equal sizes, slow (blue), medium (magenta) and fast (cyan). (B) Boxplots comparing protein turnover rates to the inter-age difference in methionine oxidation stoichiometries. Pairwise Wilcoxon signed-rank tests were performed, and p-values associated with the difference in means between groups are shown as bars above each plot.

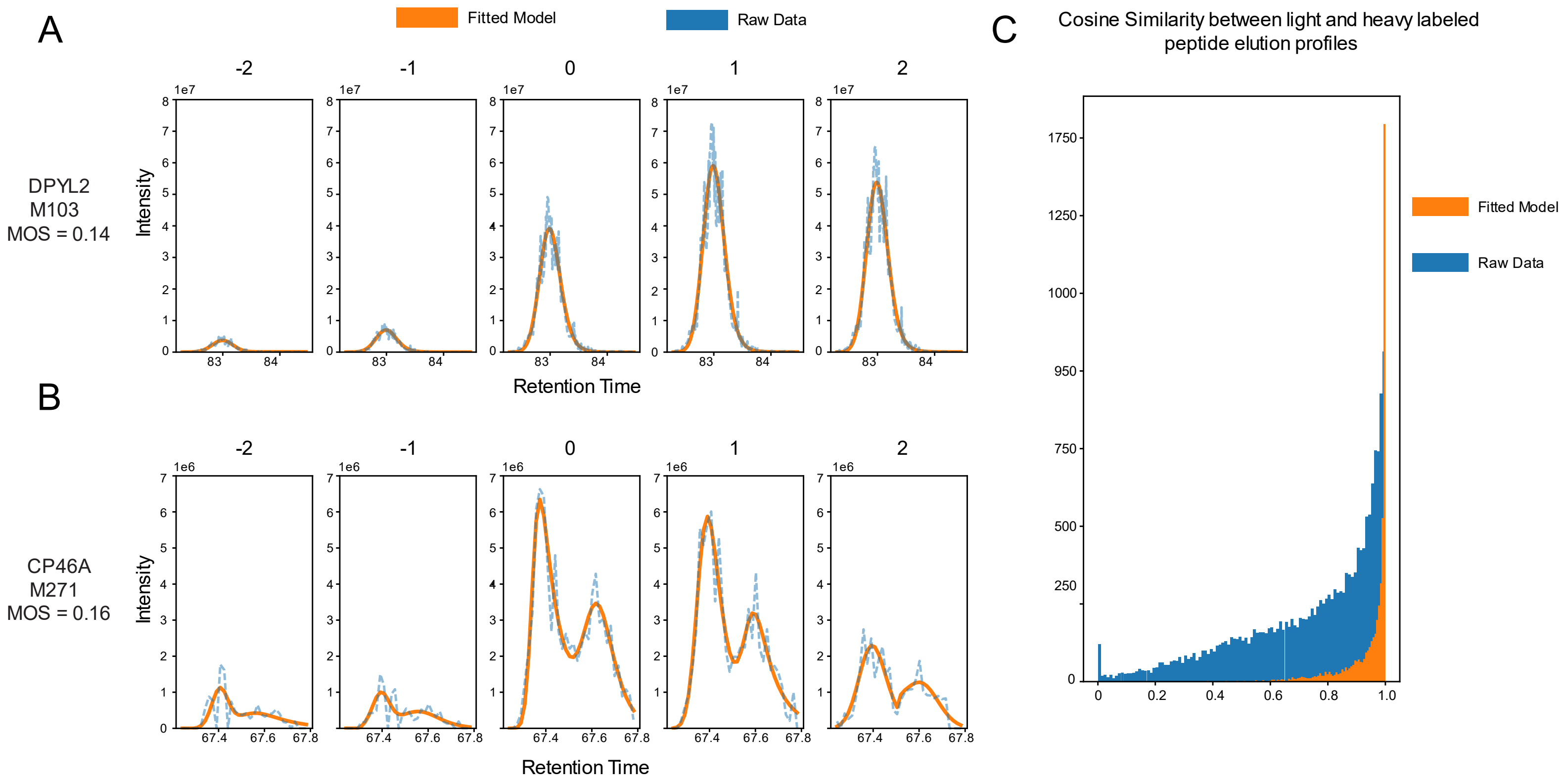

A
Fitted Model
Raw Data
-2
-1
0
1
2
1e7
DPYL2
M103
MOS = 0.14
Intensity
Retention Time
B
1e6
CP46A
M271
MOS = 0.16
Intensity
Retention Time
C
Cosine Similarity between light and heavy labeled peptide elution profiles
Fitted Model
Raw Data

**Supplementary Figure 4. Chromatographic peak modelling functions of a MobB workflow.** (A-B) The result of extracted ion chromatogram (XIC) modeling for a methionine sulfoxide containing peptide that does (B) or does not (A) have stereospecific retention times. Raw data is shown as a blue dotted line and the fitted model is shown as a solid orange line. Each graph is a unique isotopologue of the same peptide and isotope indexes, relative to the monoisotopic-heavy labeled peptide, are shown above each graph. All data was taken from a 10% titration point sample of a young mouse. (C) A histogram illustrating the improvement in isotope cluster assembly when using the XIC modelling functions of MobB. Cosine similarities between light and heavy labeled isotope pairs is used as the quality metric. Modeled data is shown in orange and unmodeled data is shown in blue. Data for all samples is visualized.

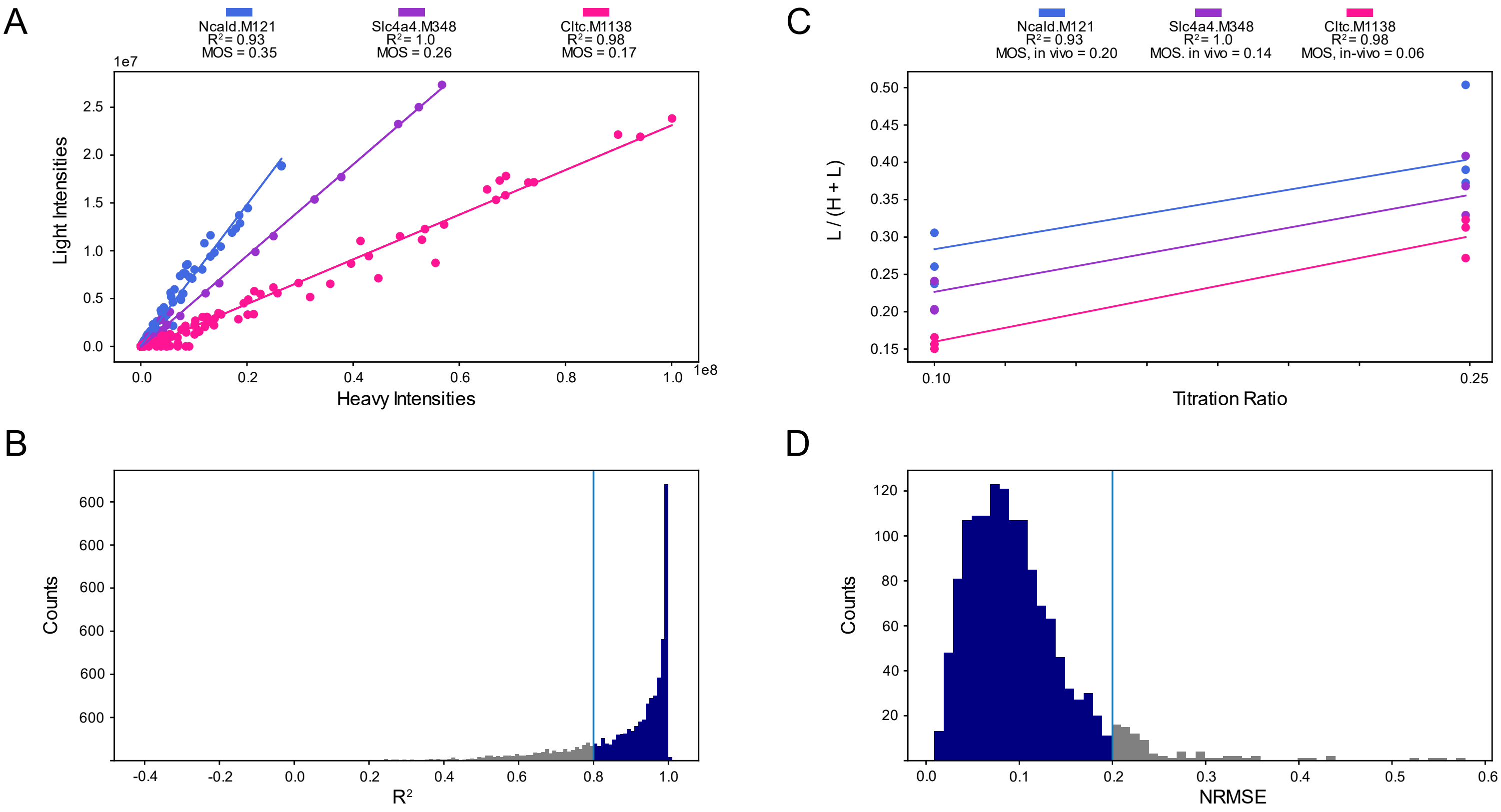

A
Ncald.M121
R² = 0.93
MOS = 0.35
Slc4a4.M348
R² = 1.0
MOS = 0.26
Cltc.M1138
R² = 0.98
MOS = 0.17
Light Intensities
Heavy Intensities
B
Counts
R²
C
Ncald.M121
R² = 0.93
MOS, in vivo = 0.20
Slc4a4.M348
R² = 1.0
MOS. in vivo = 0.14
Cltc.M1138
R² = 0.98
MOS, in-vivo = 0.06
L / (H + L)
Titration Ratio
D
Counts
NRMSE

**Supplementary Figure 5. Strategy for the estimation of in-vivo methionine oxidation stoichiometries (MOS).** (A) An example of the linear regression strategy used to estimate MOS values in each sample. Peptides with high (blue), moderate (purple) or low (pink) MOS values are shown as illustrative examples. All data was taken from a 10% titration point sample of a young mouse. (B) The global distribution of quality scores ($R^2$) for linear models used to measure MOS values. All data was taken from a 10% titration point sample of a young mouse. The blue horizontal line indicates the quality cutoff of $R^2 >= 0.8$. (C) An example of the titration response strategy used to estimate in-vivo methionine oxidation stoichiometries (MOS) for each peptide. Peptides with high (blue), moderate (purple) or low (pink) MOS values are shown as illustrative examples. All data shown was taken from the young age group. (D) The global distribution of quality scores (NRMSE) for peptides quantified in the young age group. The blue horizontal line indicates the quality cutoff of NRMSE <= 0.2.

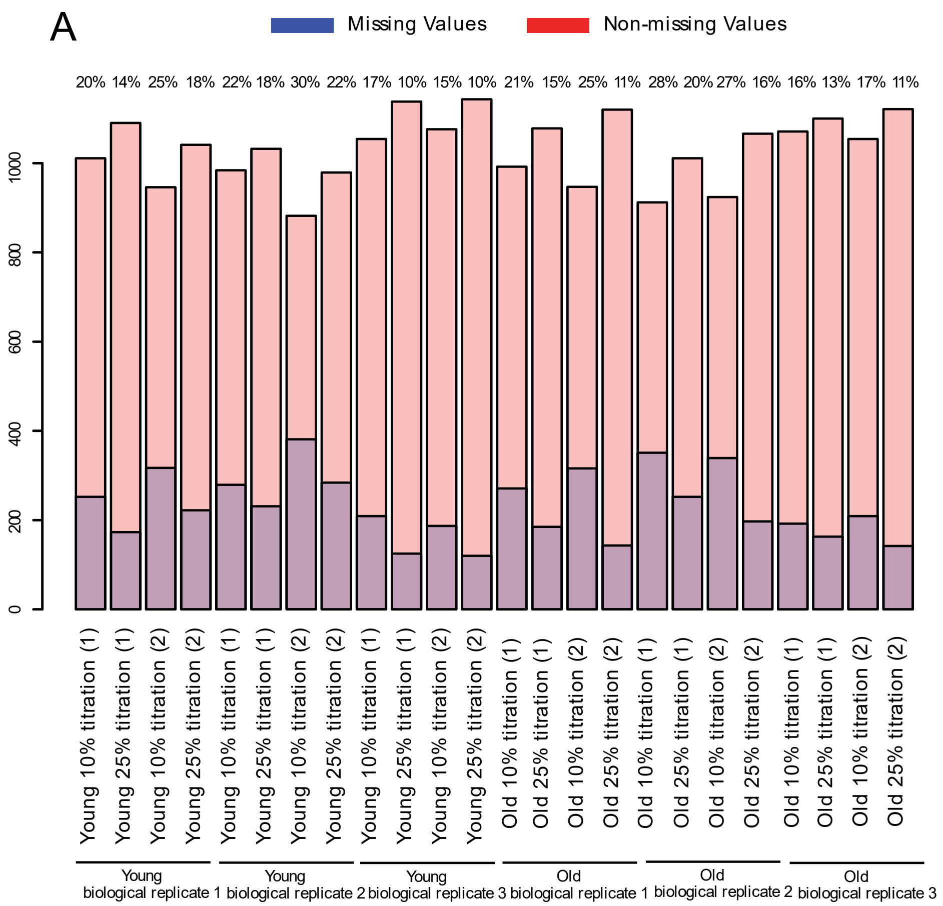

A
Missing Values
Non-missing Values
20% 14% 25% 18% 22% 18% 30% 22% 17% 10% 15% 10% 21% 15% 25% 11% 28% 20% 27% 16% 16% 13% 17% 11%
0
200
400
600
800
1000
Young 10% titration (1)
Young 25% titration (1)
Young 10% titration (2)
Young 25% titration (2)
Young 10% titration (1)
Young 25% titration (1)
Young 10% titration (2)
Young 25% titration (2)
Young 10% titration (1)
Young 25% titration (1)
Young 10% titration (2)
Young 25% titration (2)
Old 10% titration (1)
Old 25% titration (1)
Old 10% titration (2)
Old 25% titration (2)
Old 10% titration (1)
Old 25% titration (1)
Old 10% titration (2)
Old 25% titration (2)
Old 10% titration (1)
Old 25% titration (1)
Old 10% titration (2)
Old 25% titration (2)
Young biological replicate 1
Young biological replicate 2
Young biological replicate 3
Old biological replicate 1
Old biological replicate 2
Old biological replicate 3


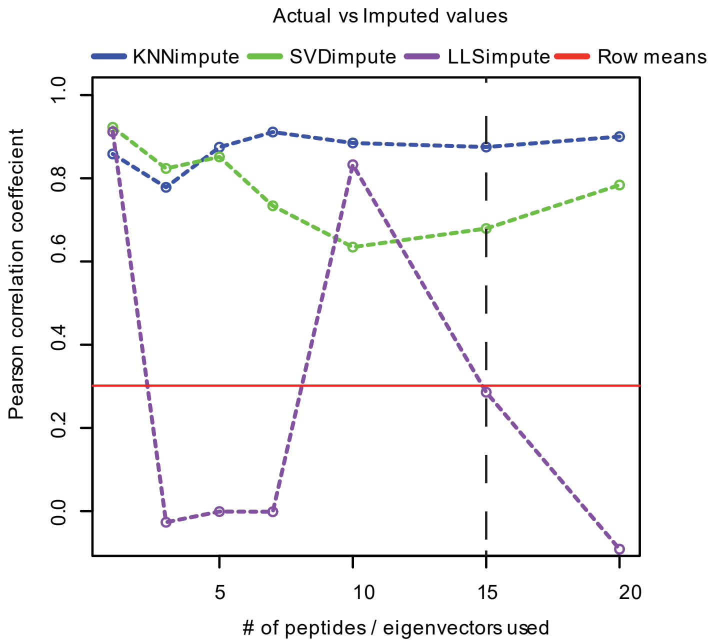

Actual vs Imputed values
KNNimpute
SVDimpute
LLSimpute
Row means
Pearson correlation coeffecient
0.0
0.2
0.4
0.6
0.8
1.0
5
10
15
20
### of peptides / eigenvectors used

**Supplementary Figure 6. A summary of missing value density and imputation.** (A) Bar chart illustrating the density of missing values (MV) in the final dataset for each sample. In order to be included in the final dataset a peptide must have been quantified in 7 out of 12 possible samples for both age groups. Percentage values for MV densities are shown at the top of each bar. MV densities range from 11%-30%, with an average of 16%. (B) A line plot illustrating the performance of three commonly used algorithms for imputing MVs in proteomic datasets. The Pearson correlation coefficient between the imputed values calculated for artificially created MVs and the actual observed experimental values for those peptides is shown on the y-axis. Parameter selection for each algorithm is shown on the x-axis. KNNimpute and LLSimpute each require a parameter specifying the number of similar peptides used when imputing MVs and SVDimpute requires a parameter specifying the number of eigenvectors to be used when imputing MVs. For reference the Pearson correlation coefficient returned when replacing missing values with row means is shown as a red line. The black dotted line shows the optimal parameter (KNNimpute, k=15) selected for imputing MVs in the final dataset.
