## Supplementary Code for "Accurate proteome-wide measurement of methionine oxidation in aging mouse brains": MObBcore_analysis.py

```
# -*- coding: utf-8 -*-
"""
Created on Thu Apr  1 07:57:32 2021

@author: jbett
"""
#import modules
import pandas as pd
from pyteomics import mzxml, auxiliary
import numpy as np
import math as math
import matplotlib as mp
import mpl_toolkits.mplot3d.art3d as art3d
import scipy as sp
from sklearn import metrics
import statistics as st
import matplotlib.patches as mpl_patches
import matplotlib.gridspec as gridspec
from matplotlib.lines import Line2D

from scipy.special import erf
from scipy.optimize import curve_fit
from scipy.signal import find_peaks, peak_prominences
from scipy.stats import pearsonr
from scipy import stats
from numpy import trapz
import pickle


def Createtheoretical(fn,theodist):
    dist1=np.pad(theodist, (0, 2), 'constant')
    dist2=np.pad(theodist, (2, 0), 'constant')
    dist3=(dist1*fn) + (dist2*(1 - fn));
    return dist3

def RTanalysis(modelin,timesarray):
    RTlength=max(timesarray)-min(timesarray)
    XIClight=modelin[0]+modelin[1]
    for n in range(2,len(modelin)):
        if n == 2:
            XICheavy=modelin[n]
        else:
            XICheavy=XICheavy+modelin[n]
            
    RTcos = 1-sp.spatial.distance.cosine(XIClight,XICheavy)
    weightedscansl = sum((timesarray * XIClight))/sum(XIClight)
    weightedscansh = sum((timesarray * XICheavy))/sum(XICheavy)
    RTdiff = abs(weightedscansl-weightedscansh)/RTlength
    return RTcos,RTdiff


def asym_peak(t, pars):
    'from Anal. Chem. 1994, 66, 1294-1301'
    a0 = pars[0]  # peak area
    a1 = pars[1]  # elution time
    a2 = pars[2]  # width of gaussian
    a3 = pars[3]  # exponential damping term
    f = (a0/2/a3*np.exp(a2**2/2.0/a3**2 + (a1 - t)/a3)
         *(erf((t-a1)/(np.sqrt(2.0)*a2) - a2/np.sqrt(2.0)/a3) + 1.0))
    return f

#To get two peaks, we simply add two peaks together.

def two_peaks(t, *pars):    
    'function of two overlapping peaks'
    a10 = pars[0]  # peak area
    a11 = pars[1]  # elution time
    a12 = pars[2]  # width of gaussian
    a13 = pars[3]  # exponential damping term
    a20 = pars[4]  # peak area
    a21 = pars[5]  # elution time
    a22 = pars[6]  # width of gaussian
    a23 = pars[7]  # exponential damping term   
    p1 = asym_peak(t, [a10, a11, a12, a13])
    p2 = asym_peak(t, [a20, a21, a22, a23])
    return p1 + p2


def asym_peakx(t, *pars):
    'from Anal. Chem. 1994, 66, 1294-1301'
    a0 = pars[0]  # peak area
    a1 = pars[1]  # elution time
    a2 = pars[2]  # width of gaussian
    a3 = pars[3]  # exponential damping term
    f = (a0/2/a3*np.exp(a2**2/2.0/a3**2 + (a1 - t)/a3)
         *(erf((t-a1)/(np.sqrt(2.0)*a2) - a2/np.sqrt(2.0)/a3) + 1.0))
    return f


def modelpeaks(feat):
    intsfitlist=[]
    intslist=[]
    areas=[]
    maxpeaks=[]
    intsarray=[]
    cosarray=[]
    feat=feat
    areas=[]
    maxiso=feat[:,4].max()
    maxiso=maxiso.astype(np.int64)+1
    r2list=[]
    r22list=[]
    featmodel = {}
    for n in range(-2,3):
        featsub=feat[:,4]==n
        featsubs=feat[featsub,:]
        times=featsubs[:,3]
        ints=featsubs[:,1]
        peaks,_=find_peaks(ints)
        prominences = peak_prominences(ints, peaks)[0]
        centers=pd.DataFrame({'peaks':peaks,'prominences':prominences})
        centers=centers.sort_values(by='prominences')
        centers=centers.reset_index()
        if len(centers)>0:
            RT1=centers['peaks'][len(centers)-1]
        else:
            break
        if len(centers) > 1:
            RT2=centers['peaks'][len(centers)-2]
        prominences.sort()
        

        
        parguess = (np.trapz(featsubs[:,1],times), featsubs[RT1,3], 0.05, 0.05)
        try:
            popt, pcov = curve_fit(asym_peakx, times, ints, parguess)
        except RuntimeError:
            popt=[]
            pcov=[]
        
        
        if len(popt)==0:
            intsfit = np.nan
            cor = 0
        if len(popt)>0:
            intsfit=asym_peakx(times,*popt)
            cor,_=pearsonr(ints,intsfit)
            if cor >= 0.9 or len(centers) <= 1:
                peak1 = asym_peakx(times, *popt)
                featmodel["{}".format(n)] = {}
                featmodel["{}".format(n)]['peak1'] = peak1
                featmodel["{}".format(n)]['peak2'] = np.zeros(len(times))
                
                    
        if cor < 0.9 and len(centers) > 1:
            parguess2 = (np.trapz(featsubs[:,1],times)/2, featsubs[RT1,3], 0.05, 0.05,
                    np.trapz(featsubs[:,1],times)/2, featsubs[RT2,3], 0.05, 0.05)
            
            try:
                popt2, pcov = curve_fit(two_peaks, times, ints, parguess2)
            except:
                print("No model could be fit")
                popt2=[]
                pcov=[]
                pass
                
            intsfit2=two_peaks(times,*popt2)
            cor2,_=pearsonr(ints,intsfit2)
            if cor2 > cor:
                intsfit=intsfit2
                popt=popt2
                
                peak1pars = popt[0:4]
                peak1 = asym_peakx(times,*peak1pars)
                
                peak2pars = popt[4:8]
                peak2 = asym_peakx(times,*peak2pars)
                
                
                featmodel["{}".format(n)] = {}
                featmodel["{}".format(n)]['peak1'] = peak1
                featmodel["{}".format(n)]['peak2'] = peak2
    

    path1=[]
    path2=[]
    xx=-1
    for z in range(-2,3):
        isoin = featmodel['{}'.format(z)]
        in1=isoin['peak1']
        in2=isoin['peak2']
        if z == -2:
            path1.append(in1)
            path2.append(in2)
        else:
            adj1=path1[xx]
            adj2=path2[xx]
            if 1-sp.spatial.distance.cosine(in1,adj1)>=0.6 or 1-sp.spatial.distance.cosine(in2,adj1)>=0.6:
                if 1-sp.spatial.distance.cosine(in1,adj1)>=0.6 and 1-sp.spatial.distance.cosine(in2,adj1)>=0.6:
                    path1.append(in1+in2)
                else:
                    if 1-sp.spatial.distance.cosine(in1,adj1)>=0.6:
                        path1.append(in1)
                    if 1-sp.spatial.distance.cosine(in2,adj1)>=0.6:
                        path1.append(in2)
            else:
                path1.append(np.zeros(len(times)))
                
            if 1-sp.spatial.distance.cosine(in1,adj2)>=0.6 or 1-sp.spatial.distance.cosine(in2,adj2)>=0.6:
                if 1-sp.spatial.distance.cosine(in1,adj1)>=0.6 and 1-sp.spatial.distance.cosine(in2,adj1)>=0.6:
                    path1.append(in1+in2)
                else:               
                    if 1-sp.spatial.distance.cosine(in1,adj2)>=0.6:
                        path2.append(in1)
                    if 1-sp.spatial.distance.cosine(in2,adj2)>=0.6:
                        path2.append(in2)
            else:
                path2.append(np.zeros(len(times)))
        xx=xx+1


    graph=[]
    graphsub=[]
    for g in range(0,len(path1)):
        if sum(path1[g])>0:
            graphsub.append(path1[g])
    if len(graphsub)>4:
        graph.append(graphsub)
    
    graphsub=[]
    for g in range(0,len(path2)):
        if sum(path2[g])>0:
            graphsub.append(path2[g])
    if len(graphsub)>4:
        graph.append(graphsub)
    
    finalmodel=[]
    peakmaxs=[]
    if len(graph) > 1:
        for x in range(0,len(graph[0])):
                finalmodelx = graph[0][x]+graph[1][x]
                finalmodel.append(finalmodelx)
                peakmaxs.append(max(graph[0][x])+max(graph[1][x]))
    else:
        for x in range(0,len(graph[0])):
                finalmodelx = graph[0][x]
                finalmodel.append(finalmodelx)
                peakmaxs.append(max(graph[0][x]))
            

        
    return finalmodel, times,peakmaxs

#Create methods

#range with non-integer step size
def frange(start, stop, step):
     i = start
     while i < stop:
         yield i
         i += step
#mean
def mean(lst): 
    return sum(lst) / len(lst)

#dot product (Cosine Similarity)
def dotproduct(v1, v2):
  return sum((a*b) for a, b in zip(v1, v2))

# length (Cosine Similarity)
def length(v):
  return math.sqrt(dotproduct(v, v))

#angle (Cosine Similarity)
def angle(v1, v2):
  return math.acos(dotproduct(v1, v2) / (length(v1) * length(v2)))


#find closest value (K) in list (lst) (not currently used)
def closest(lst, K): 
      return min(range(lst.idxmin(),lst.idxmax()), key = lambda i: abs(lst[i]-K))

#Get atomic composition
def getatomcomp(seq,AAcompmap):
    atomcomp=list()
    for q in range(0,len(seq)):
        Carbon=AAcompmap[seq[q]][0]
        Hydrogen=AAcompmap[seq[q]][1]
        Nitrogen=AAcompmap[seq[q]][2]
        Oxygen=AAcompmap[seq[q]][3]
        Sulfur=AAcompmap[seq[q]][4]
        atomcomp.append([Carbon,Hydrogen,Nitrogen,Oxygen,Sulfur])
    
    atomcomp=[ sum(row[i] for row in atomcomp) for i in range(len(atomcomp[0])) ]
    return atomcomp

#Grab swath
def getswath(RTS,RTE,spectrumhandle):
    swath = spectrumhandle.time[RTS:RTE]    
    
    return swath

#filter swath for mz values within tolerance
def filterswath(swath,mzs,cutoff):
    feat=[]
    for g in mzs:
        isoindex=np.flatnonzero(g==mzs)[0]-2
    
        for m in range(0,len(swath)):
            scan=[]
            ms1mzarray=swath[m]['m/z array']
            ms1intsarray=swath[m]['intensity array']
            for j in range(0,len(ms1mzarray)):
                if (ms1mzarray[j]>g-(g*cutoff/1000000))&(ms1mzarray[j]<g+(g*cutoff/1000000)):
                    mz=ms1mzarray[j]
                    intensity=ms1intsarray[j]
                    scannum=int(swath[m]['num'])
                    RT=float(swath[m]['retentionTime'])
                    scan=[mz,intensity,scannum,RT,isoindex]
                
                if not scan:
                    mz=g
                    intensity = 0
                    scannum=int(swath[m]['num'])
                    RT=float(swath[m]['retentionTime'])
                    scan=[mz,intensity,scannum,RT,isoindex]
            
            feat.append(scan)
            
    if not feat:
        feature=None
    else:
        feature=np.vstack(feat)
        return feature


#filter consecutive scans based on cossine similiarity (not currently used)
def scanfilter(feat,cutoff):
    monoiso=feat[feat[:,4]==0]
    featweightedt=monoiso[:,2]*monoiso[:,1]
    centerscan=sum(featweightedt)/sum(monoiso[:,1])
    nearestscan=[index for index,value in enumerate(feat[:,2]) if abs(value-centerscan) ==min(abs(feat[:,2]-centerscan))][0]  
    feat=pd.DataFrame(feat)
    featgb = feat.groupby(2)    
    featframe=[featgb.get_group(x) for x in featgb.groups]
    keeper=[]
    for n in range(nearestscan+1,len(featframe)):
        scanstart=featframe[nearestscan]
        scanin=featframe[n]
        patternstart=scanstart[1]/sum(scanstart[1])
        patternin=scanin[1]/sum(scanin[1])
        sim=angle(patternstart,patternin)
        if sim<=cutoff:
            keeper.append(featframe[n])
        else:
            break
    for n in range(nearestscan-1,0,-1):
        scanstart=featframe[nearestscan]
        scanin=featframe[n]
        patternstart=scanstart[1]/sum(scanstart[1])
        patternin=scanin[1]/sum(scanin[1])
        sim=angle(patternstart,patternin)
        if sim<=cutoff:
            keeper.append(featframe[n])
        else:
            break
        
    keeper.append(featframe[nearestscan])
    
    
    featreturn=np.vstack(keeper)
    return featreturn

#get the theoretical isotope distribution
def gettheodist(atomcomp,cutoff):
    form=[[12.0,13.0035,0.99,atomcomp[0]],[1.007825,2.014102,0.99985,atomcomp[1]],[14.003074,15.000109,0.99634,atomcomp[2]],[15.9949,17.999,0.99762,atomcomp[3]],[31.972,33.9679,0.9502,atomcomp[4]]];
    temp3=[]
    for i in range(0,len(form)):
        atomset=form[i]
        mainmass=atomset[0]
        altmass=atomset[1]
        abundancemainmass=atomset[2]
        abundancealtmass=1-abundancemainmass
        atoms=atomset[3]
        temp2=[]
        
        for j in range(0,atoms):
            temp1=[];
            combs= math.factorial(atoms)/(math.factorial(j)*math.factorial(atoms-j));
            prob= (abundancemainmass**(atoms-j))*(abundancealtmass**j);
            odds=combs*prob;
            mass = (altmass*j)+(mainmass*(atoms-j));
            if odds > cutoff:
                temp1=[j,mass,odds];
                temp2.append(temp1);
                
        temp3.append(temp2)
            
    temp4=[]
    
    for ci in range(0,len(temp3[0])):
        cset=temp3[0][ci];
        cmass=cset[1];
        codds=cset[2];
        for hi in range(0,len(temp3[1])):
            hset=temp3[1][hi];
            hmass=hset[1];
            hodds=hset[2];
            for ni in range(0,len(temp3[2])):
                nset=temp3[2][ni];
                nmass=nset[1];
                nodds=nset[2];
                for oi in range(0,len(temp3[3])):
                    oset=temp3[3][oi];
                    omass=oset[1];
                    oodds=oset[2];
                    for si in range(0,len(temp3[4])):
                        sset=temp3[4][si];
                        smass=sset[1];
                        sodds=sset[2];
                        tmass= round(cmass+hmass+nmass+omass+smass);
                        todds=codds*hodds*nodds*oodds*sodds;
                        temp4.append([tmass,todds]);
                        
    temp4=pd.DataFrame(temp4)
    temp4=temp4.groupby([0])[1].sum()
    theodist=pd.DataFrame({'Mass':temp4.index,'odds':temp4})
    theodist=theodist.reset_index()
    theodist=theodist.drop(columns=0)
    theodist=theodist[theodist['odds']>=cutoff]
    return theodist

#get theoretical m/z values for 16O/18O isotope pairs
def getMZs(theodist,mz0,indexoffset):
   mzs=[]
   if indexoffset==0:
       for y in range(-2,len(theodist)):
           mzn=mz0+(y/charge)
           mzs.append(mzn)
   else:
        for y in range(0,len(theodist)+2):
            mzn=mz0+(y/charge)
            mzs.append(mzn)
   return mzs        

#measure the distance between XIC centers
def RTshift(feat):
    RTlength=max(feat[:,3])-min(feat[:,3])
    featweightedt=feat[:,3]*feat[:,1]
    if sum(featweightedt)!=0:
        featweightedt=sum(featweightedt)/sum(feat[:,1])
        feat=pd.DataFrame(feat)
        featgb = feat.groupby(4)    
        featframe=[featgb.get_group(x) for x in featgb.groups]
        
        light=featframe[0:2]
        lights=pd.DataFrame()
        for n in range(0,len(light)):
            isoin=light[n]
            lights=lights.append(isoin)
        isoweighted=sum(lights[3]*lights[1])
        if sum(lights[1])!=0:
            isocenterlight=isoweighted/sum(lights[1])
        else:
            return 0
        
        heavy=featframe[2:len(featframe)]
        heavys=pd.DataFrame()
        for n in range(0,len(heavy)):
            isoin=heavy[n]
            heavys=heavys.append(isoin)
        isoweighted=sum(heavys[3]*heavys[1])
        if sum(heavys[1])!=0:
            isocenterheavy=isoweighted/sum(heavys[1])
        else:
            return np.nan
        
        if sum(lights[1])!=0 and sum(heavys[1])!=0:
           RTshift=abs(isocenterheavy-isocenterlight)/RTlength
           return RTshift

#measure cosine similarity in XIC
def RTcosx(feat):
    feat=pd.DataFrame(feat)
    featgb = feat.groupby(4)    
    featframe=[featgb.get_group(x) for x in featgb.groups]
            
    light=featframe[0:2]
    lights=pd.DataFrame()
    for n in range(0,len(light)):
        isoin=light[n]
        lights=lights.append(isoin)
        
    
    heavy=featframe[2:len(featframe)]
    heavys=pd.DataFrame()
    for n in range(0,len(heavy)):
        isoin=heavy[n]
        heavys=heavys.append(isoin)
        
    
    XIClight = pd.DataFrame(lights)\
          .groupby(2, as_index=False)\
          .sum()\
          .reset_index()
    
   
          
    XICheavy = pd.DataFrame(heavys)\
          .groupby(2, as_index=False)\
          .sum()\
          .reset_index() 
          
    
    cossim=1-sp.spatial.distance.cosine(XIClight[1],XICheavy[1])
    heavyscans = XICheavy[2]
    lightscans = XIClight[2]
    heavyints = XICheavy[1]
    lightints = XIClight[1]
    XICheavy = pd.concat([heavyscans,heavyints],axis=1)
    XIClight = pd.concat([lightscans,lightints],axis=1)
    return [cossim,XIClight,XICheavy]
         
#Connect isoprotomers if they improve fit to theoretical
def filterisotopesMZ(feat,theomodel):
    if sum(feat[:,1])!=0:
        featweightedt=feat[:,3]*feat[:,1]
        featweightedt=sum(featweightedt)/sum(feat[:,1])
        feat=pd.DataFrame(feat)
        featgb = feat.groupby(4)    
        featframe=[featgb.get_group(x) for x in featgb.groups]
        isolist=[]
        mzlist=[]
        
        spectra2d=[]
        spectra2dt=[]
        for i in range(0,len(featframe)):
            isoin=featframe[i]
            isoin=isoin.reset_index()
            isolist.append(sum(isoin[1]))
            mzlist.append(isoin[4][0])
        if sum(isolist)!=0:
            feat=[]
            for isoprot in range(0,len(mzlist)):
                if mzlist[isoprot]<0:
                    feat.append(featframe[isoprot])
                    spectra2d.append(isolist[isoprot])
                else:
                    spectra2dt.append(isolist[isoprot])
                    theomodelv=theomodel['odds']/sum(theomodel['odds'])
                    if sum(np.array(spectra2dt)) != 0:
                        isolistv=np.array(spectra2dt)/sum(np.array(spectra2dt))
                        isolistvcopy=np.pad(isolistv, (0, (len(theomodel)-len(isolistv))), 'constant')
                        isolistv=isolistvcopy
                    else:
                        isolistv=np.array(spectra2dt)
                        isolistvcopy=np.pad(isolistv, (0, (len(theomodel)-len(isolistv))), 'constant')
                        isolistv=isolistvcopy
                    ls=metrics.mean_squared_error(theomodelv,isolistv)
                    
                    if mzlist[isoprot]==0:
                        lskp=ls
                        feat.append(featframe[isoprot])
                    if mzlist[isoprot]!=0:
                        if ls<lskp:
                            lskp=ls
                            feat.append(featframe[isoprot])
            if not feat:
                feat=None
                return feat
            else:
                feat=np.vstack(feat)
                return feat

#get fraction oxidized
def EstimateFractionOxidized(spectra2d,theodist):
    theocopy=theodist/sum(theodist)
    spectra2d=spectra2d[0:len(theocopy)]
    intscopy=spectra2d/sum(spectra2d)
    
    fnkp=0
    lskp=100
    for n in range(0,1000):
        lslist=[]
        fn=n/1000
        dist1=np.pad(theocopy, (0, 2), 'constant')
        dist2=np.pad(theocopy, (2, 0), 'constant')
        dist3=(dist1*fn) + (dist2*(1 - fn));
        dist3copy=dist3[0:len(intscopy)]
        dist3=dist3copy/sum(dist3)
        for idx in range(0,len(intscopy)):
            lslist.append((dist3[idx]-intscopy[idx])**2)
            
        ls=sum(lslist);
        if fn==0:
            lskp = ls;
            fnkp = fn;
            theocopykeep = dist3;
            observed = intscopy;
               
        if ls < lskp:
            lskp = ls;
            fnkp = fn;
            theocopykeep = dist3;
            observed = intscopy;
    return [fnkp,lskp,theocopykeep,observed]

#create data report for peptide features          
def printanalysis(n):
    model=featarray[n]
    xich=XIChs[n]
    xicl=XICls[n]
    observed=spectra2ds[n]
    theoretical=theoreticals[n]
    fig = mp.figure()
    gs = fig.add_gridspec(2, 2)
    ax = fig.add_subplot(gs[0,:], projection='3d')
    for xi, yi, zi in zip(model[:,4], model[:,2], model[:,1]):        
        line=art3d.Line3D(*zip((xi, yi, 0), (xi, yi, zi)), marker='o', markevery=(1, 1))
        ax.add_line(line)
    ax.set_xlim3d(min(model[:,4]), max(model[:,4]))
    ax.set_ylim3d(min(model[:,2]), max(model[:,2]))   
    ax.set_zlim3d(0, max(model[:,1])) 
    ax.view_init(azim=-80)
    
    handles = [mpl_patches.Rectangle((0, 0), 1, 1, fc="white", ec="white", 
                                 lw=0, alpha=0)] * 4
    labels = []
    labels.append('Sequence = '+df['Sequence'][n])
    labels.append('Protein = '+df['Proteins'][n])
    labels.append('Rawfile = '+rawfile+'.mzXML')
    labels.append('Fraction Oxidized = '+str(fracoxlist[n]))

    ax.legend(handles, labels, loc='upper right', fontsize='small', 
          fancybox=True, framealpha=0.7, 
          handlelength=0, handletextpad=0)
    
    ax = fig.add_subplot(gs[1,0])
    mp.plot(xich[2],xich[1])
    mp.plot(xicl[2],xicl[1])
    handles = [mpl_patches.Rectangle((0, 0), 1, 1, fc="white", ec="white", 
                                 lw=0, alpha=0)] * 4
    labels = []
    labels.append('Sequence = '+df['Sequence'][n])
    labels.append('Protein = '+df['Proteins'][n])
    labels.append('RT correlation = '+str(RTcoss[n]))
    labels.append('RT shift = '+str(RTshifts[n]))
    legend1 = mp.legend(handles, labels, loc='upper left', fontsize='small', 
          fancybox=True, framealpha=0.7, 
          handlelength=0, handletextpad=0)
    custom_lines = [Line2D([0], [0], color='orange', lw=4),
                    Line2D([0], [0], color='blue', lw=4)]
    legend2 = ax.legend(custom_lines, ['Light', 'Heavy'],loc='upper right')
    ax.add_artist(legend1)
    ax.add_artist(legend2)
    
    ax = fig.add_subplot(gs[1,1])
    x1=range(-2,len(observed)-2)
    y1=observed
    y2=theoretical
    handles = [mpl_patches.Rectangle((0, 0), 1, 1, fc="white", ec="white", 
                                 lw=0, alpha=0)] * 3
    labels = []
    labels.append('Sequence = '+df['Sequence'][n])
    labels.append('Protein = '+df['Proteins'][n])
    labels.append('RSS = '+str(fitlist[n]))   
    
    legend1 = mp.legend(handles, labels, fontsize='small', 
          fancybox=True, framealpha=0.7, 
          handlelength=0, handletextpad=0,loc='upper center')
    custom_lines = [Line2D([0], [0], color='blue', lw=4),
                Line2D([0], [0], color='green', lw=4)]
    legend2 = ax.legend(custom_lines, ['Observed data', 'Theoretical data'],loc='upper right')
    ax.add_artist(legend1)
    ax.add_artist(legend2)
    mp.stem(x1,y1,'b', markerfmt='bo', label='observed data')
    mp.stem(x1,y2,'g', markerfmt='go', label='theoretical')
    mp.show()

#plot XIC for peptide features
def plotXIC(xicl,xich):
    mp.plot(xich[2],xich[1])
    mp.plot(xicl[2],xicl[1])
    mp.show()
                                                  


# dictionary data class with function to add
class my_dictionary(dict):  
  
    # __init__ function  
    def __init__(self):  
        self = dict()  
          
    # Function to add key:value  
    def add(self, key, value):  
        self[key] = value 
        
#amino acid to atomic composition map
A= [3,7,1,2,0]
R= [6,14,4,2,0]
N= [4,8,2,3,0]
D= [4,7,1,4,0]
C= [3,7,1,2,1]
Q= [5,10,2,3,0]
E= [5,9,1,4,0]
G= [2,5,1,2,0]
H= [6,9,3,2,0]
I= [6,13,1,2,0]
L= [6,13,1,2,0]
K= [6,14,2,2,0]
M= [5,11,1,2,1]
F =[9,11,1,2,0]
P= [5,9,1,2,0]
S= [3,7,1,3,0]
T= [4,9,1,3,0]
W= [11,12,2,2,0]
Y= [9,11,1,3,0]
V= [5,11,1,2,0]
aakeys=['A','R','N','D','C','Q','E','G','H','I','L','K','M','F','P','S','T','W','Y','V']
aaempiricalvalues =[A,R,N,D,C,Q,E,G,H,I,L,K,M,F,P,S,T,W,Y,V]
AAcompmap=dict(zip(aakeys,aaempiricalvalues))


    
#read in evidence data
fields = ['Sequence','Proteins','Charge','Acetyl (Protein N-term)','O18(M)','Oxidation (M)','PEP','m/z','Calibrated retention time start','Calibrated retention time finish','Intensity','MS/MS scan number','Retention time','Raw file','Missed cleavages','Retention time calibration','Mass error [ppm]']
df = pd.read_table('C://Users/jbett/Desktop/mouseMOBB/newresults/evidenceYM1_1_10.txt')

#subset evidence data for peptides of interest
mets = df['Sequence'].str.count("M")
df['mets']=mets
df=df[df['mets']==1]
df = df[~df['Sequence'].str.contains("C")]
df = df[~df['Proteins'].str.contains("CON",na=True)]
df = df[~df['Proteins'].str.contains("REV",na=True)]
df['OX']=df['O18(M)']+df['Oxidation (M)']
df.sort_values(by='Raw file',inplace=True)
df=df.reset_index()

dfcopy=df[df['Missed cleavages']==0]
dfcopy=df
uniqueseqs=np.unique(dfcopy['Sequence'])


#create empty arrays
fracoxlist=[]
atomlib=my_dictionary()
featarray=[]
fitlist=[]
RTshifts=[]
RTcoss=[]
XIChs=[]
XICls=[]
LightInt=[]
theoreticals=[]
spectra2ds=[]
theolist=[]
openspec='none'

for n in range(0,len(df)):
    #for each peptide read in data used to build search
    seq=df['Sequence'][n]
    protein = df['Proteins'][n]
    charge=df['Charge'][n]
    RTS=df['Calibrated retention time start'][n]-df['Retention time calibration'][n]
    RTE=df['Calibrated retention time finish'][n]-df['Retention time calibration'][n]
    rtmid=(RTS+RTE)/2
    rawfile=df['Raw file'][n]
    mz0=df['m/z'][n]
    feat=[]
    
    #open rawfile and convert spectra to readable format
    if openspec != rawfile:
        f = mzxml.read('C://Users/jbett/Desktop/yeastMOBB/'+rawfile+'.mzXML',use_index=True)
        openspec=rawfile
    
    #for each peptide get the atomic composition    
    atomcomp=getatomcomp(seq,AAcompmap) 
    
    #build atomic composition library as it goes (saves speed)
    if tuple(atomcomp) not in atomlib:
        theodist=gettheodist(atomcomp,0.0001)
        atomlib.add(tuple(atomcomp),theodist)
        theolist.append(theodist)
        
    if tuple(atomcomp) in atomlib:
        theodist =  atomlib[tuple(atomcomp)]
        theolist.append(theodist)
        
    
    #get theoretical m/z values for 16O/18O isotope pairs
    indexoffset=df['Oxidation (M)'][n]*(-2)
    mzs=getMZs(theodist,mz0,indexoffset)
    
    #subset raw spectra based on RT bounds
    swath=getswath(RTS,RTE,f)
    
    #subset raw spectra based on theoretical m/z values (default tolerence 5ppm)
    feat=filterswath(swath,mzs,5)
    if feat is None:
        featarray.append(feat)
        fracoxlist.append(np.nan)
        fitlist.append(np.nan)
        continue
    
    #connect successve isotope peaks only if they improve fit to theoretical isotope distribution
    feat=filterisotopesMZ(feat,theodist)
    if feat is None:
        featarray.append(feat)
        fracoxlist.append(np.nan)
        fitlist.append(np.nan)
        continue
    
    featarray.append(feat)
    print(n)

       
models=[]
peaks=[]
timesarray=[]    
for q in range(0,len(featarray)):
    finalmodel=[]
    timesx=[]
    try:
        finalmodel, timesx,peaksx = modelpeaks(featarray[q])
        models.append(finalmodel)
        timesarray.append(timesx)
        peaks.append(peaksx)
        print(q)
    except:
        models.append(np.nan)
        peaks.append(np.nan)
        timesarray.append(np.nan)
        print(q)
                
spectra2ds=[]        
for q in range(0,len(models)):
    intsarray=[]
    try:
        for x in range(0,len(models[q])):
            intensity = sum(models[q][x])
            intsarray.append(intensity)
        spectra2ds.append(intsarray)
    except:
        spectra2ds.append(np.nan)
        
fracoxlist=[]
fitlist=[]
for q in range(0,len(spectra2ds)):
    try:
       fn,ls,theo,obsrved= EstimateFractionOxidized(spectra2ds[q],theolist[q])
       fracoxlist.append(fn)
       fitlist.append(ls)
    except:
       fracoxlist.append(np.nan)
       fitlist.append(np.nan) 


theolist=[]        
for q in range(0,len(df)):
    seq=df['Sequence'][q]
    atomcomp=getatomcomp(seq,AAcompmap)
    theodist=gettheodist(atomcomp,0.0001)
    theolist.append(np.asarray(theodist['odds'][0:5]/sum(theodist['odds'][0:5])))
    print(q)
       
RTcoss=[] 
RTdiffs=[]
for q in range(0,len(models)):
    try:
        RTcos,RTdiff = RTanalysis(models[q],timesarray[q])
        RTcoss.append(RTcos)
        RTdiffs.append(RTdiff)
    except:
        RTcoss.append(np.nan)
        RTdiffs.append(np.nan)
    print(q)        

df['RTcor']=RTcoss
df['RTshift']=RTdiffs


Spectras=[]
slopes=[]
r2s=[]
p_values=[]
std_errs=[]
df['feature']=models
for n in range(0,len(uniqueseqs)):
    Sequence=uniqueseqs[n]
    target=df[df['Sequence'].str.contains(Sequence)]
    target=target.reset_index()
    spectra=[]
    for j in range(0,5):
        isoins=[]
        for i in range(0,len(target)):
            try:
                isoin=target['feature'][i][j]
                isoins=np.hstack((isoins,isoin))
            except:
                continue
        spectra.append(isoins)
        
    Spectras.append(spectra)
    print(n)


#get fraction oxidized
slopes=[]
r2s=[]
p_values=[]
std_errs=[]
totalints=[]
for n in range(0,len(Spectras)):
    try:
        Y=Spectras[n][2]+Spectras[n][3]+Spectras[n][4]
        X=Spectras[n][0]+Spectras[n][1]
        slope, intercept, r_value, p_value, std_err = stats.linregress(Y,X)
        slopes.append(slope)
        r2s.append(r_value)
        p_values.append(p_value)
        std_errs.append(std_err)
        totalints.append(sum(X)+sum(Y))
    except:
        slopes.append(np.nan)
        r2s.append(np.nan)
        p_values.append(np.nan)
        std_errs.append(np.nan)
        totalints.append(np.nan)
    print(n)
    
for n in range(0,len(slopes)):
    ins=slopes[n]
    oxnew=slopes[n]/(1+slopes[n])
    slopes[n]=oxnew     
    
#correct fraction oxidized
slopecorrections=[]
for q in range(0,len(uniqueseqs)):
    atomcomp = atomcomp=getatomcomp(uniqueseqs[q],AAcompmap)
    theoretical = gettheodist(atomcomp,0.0001)
    actuals=[]
    observed=[]
    for n in range(0,1000):
        fn=n/1000
        z=Createtheoretical(fn, theoretical)
        actuals.append(fn)
        observed.append(sum(z[:,1][0:2]))
    slope, intercept, r_value, p_value, std_err = stats.linregress(observed,actuals)
    slopecorrections.append([slope,intercept])
    print(q)

slopesnew=[]    
for q in range(0,len(slopes)):
    try:
       corrected = (slopes[q]*slopecorrections[q][0])+slopecorrections[q][1]
       slopesnew.append(corrected)
    except:
        slopesnew.append(np.nan)
    print(q)
    
    

output = pd.DataFrame(
    {'Sequence': uniqueseqs,
     'FractionOxidized':slopesnew,
     'Fit':r2s,
     'totalints':totalints
    })

#output.to_csv('C://Users/jbett/Desktop/yeastMOBB/WT3results.txt')

#with open("C://Users/jbett/Desktop/mouseMOBB/newresults/results/modelsWT3.txt", "wb") as mx:
    pickle.dump(models,mx)
    
#with open("C://Users/jbett/Desktop/mouseMOBB/newresults/results/pepmodelsWT3.txt", "wb") as sx:
    pickle.dump(Spectras,sx)
```
