## Supplementary Code for "Accurate proteome-wide measurement of methionine oxidation in aging mouse brains": MObBpost_analysis.R

```
library(broom)
library(splitstackshape)
library(DMwR2)
library(modeest)
library(pcaMethods)
standard_error <- function(x) sd(x) / sqrt(length(x))
nrmse_func <-  function(obs, pred, type = "sd") {
  
  squared_sums <- sum((obs - pred)^2)
  mse <- squared_sums/length(obs)
  rmse <- sqrt(mse)
  if (type == "sd") nrmse <- rmse/sd(obs)
  if (type == "mean") nrmse <- rmse/mean(obs)
  if (type == "maxmin") nrmse <- rmse/ (max(obs) - min(obs))
  if (type == "iq") nrmse <- rmse/ (quantile(obs, 0.75) - quantile(obs, 0.25))
  if (!type %in% c("mean", "sd", "maxmin", "iq")) message("Wrong type!")
  nrmse <- round(nrmse, 3)
  return(nrmse)
  
}


################################################################################
################################################################################
#############Read in and Format old data######################################
################################################################################
################################################################################
################################################################################
################################################################################

OM1_1_10<-read.csv("C://Users/jbett/Desktop/mouseMOBB/newresults/results/OM1_1_10.txt",header=TRUE)
OM1_1_25<-read.csv("C://Users/jbett/Desktop/mouseMOBB/newresults/results/OM1_1_25.txt",header=TRUE)
OM1_2_10<-read.csv("C://Users/jbett/Desktop/mouseMOBB/newresults/results/OM1_2_10.txt",header=TRUE)
OM1_2_25<-read.csv("C://Users/jbett/Desktop/mouseMOBB/newresults/results/OM1_2_25.txt",header=TRUE)
OM2_1_10<-read.csv("C://Users/jbett/Desktop/mouseMOBB/newresults/results/OM2_1_10.txt",header=TRUE)
OM2_1_25<-read.csv("C://Users/jbett/Desktop/mouseMOBB/newresults/results/OM2_1_25.txt",header=TRUE)
OM2_2_10<-read.csv("C://Users/jbett/Desktop/mouseMOBB/newresults/results/OM2_2_10.txt",header=TRUE)
OM2_2_25<-read.csv("C://Users/jbett/Desktop/mouseMOBB/newresults/results/OM2_2_25.txt",header=TRUE)
OM3_1_10<-read.csv("C://Users/jbett/Desktop/mouseMOBB/newresults/results/OM3_1_10.txt",header=TRUE)
OM3_1_25<-read.csv("C://Users/jbett/Desktop/mouseMOBB/newresults/results/OM3_1_25.txt",header=TRUE)
OM3_2_10<-read.csv("C://Users/jbett/Desktop/mouseMOBB/newresults/results/OM3_2_10.txt",header=TRUE)
OM3_2_25<-read.csv("C://Users/jbett/Desktop/mouseMOBB/newresults/results/OM3_2_25.txt",header=TRUE)

OM1_1_10$X<-NULL
OM1_1_25$X<-NULL
OM1_2_10$X<-NULL
OM1_2_25$X<-NULL
OM2_1_10$X<-NULL
OM2_1_25$X<-NULL
OM2_2_10$X<-NULL
OM2_2_25$X<-NULL
OM3_1_10$X<-NULL
OM3_1_25$X<-NULL
OM3_2_10$X<-NULL
OM3_2_25$X<-NULL

OM1_1_10<-OM1_1_10[!is.na(OM1_1_10$FractionOxidized),]
OM1_1_10<-OM1_1_10[OM1_1_10$Fit>=0.8,]
OM1_1_10_evi<-read.table("C://Users/jbett/Desktop/mouseMOBB/newresults/evidenceOM1_1_10.txt",sep="\t",header = TRUE)
OM1_1_10_evi_proteins<-OM1_1_10_evi$Proteins
OM1_1_10_evi_seqs<-OM1_1_10_evi$Sequence

Tt<-as.data.frame(cbind(OM1_1_10_evi_seqs,OM1_1_10_evi_proteins))
pepannot<-data.frame(sequence=character(),proteins=character())

for (n in 1:nrow(OM1_1_10_evi)) {
  keyin<-OM1_1_10_evi$Sequence[n]
  Ttsub<-Tt[Tt$OM1_1_10_evi_seqs==keyin,]
  prots<-Ttsub$OM1_1_10_evi_proteins
  protssublist<-c()
  for (q in 1:length(prots)) {
    protssub<-prots[q]
    protssublist<-c(protssublist,unlist(strsplit(protssub, ";")))
    
    
  }
  protslist<-unique(protssublist)
  pepannot[n,]<-c(as.character(keyin),paste(protslist,sep = "",collapse = ";"))
  
  
}
pepannot<-unique(pepannot)


for (n in 1:nrow(OM1_1_10)) {
  keyin<-OM1_1_10$Sequence[n]
  protsin<-pepannot$proteins[pepannot$sequence==keyin]
  OM1_1_10$Proteins[n]<-protsin
  
}


rm(Tt,pepannot,keyin,Ttsub,prots,protssublist,protslist)

OM1_1_25<-OM1_1_25[!is.na(OM1_1_25$FractionOxidized),]
OM1_1_25<-OM1_1_25[OM1_1_25$Fit>=0.8,]
OM1_1_25_evi<-read.table("C://Users/jbett/Desktop/mouseMOBB/newresults/evidenceOM1_1_25.txt",sep="\t",header = TRUE)
OM1_1_25_evi_proteins<-OM1_1_25_evi$Proteins
OM1_1_25_evi_seqs<-OM1_1_25_evi$Sequence

Tt<-as.data.frame(cbind(OM1_1_25_evi_seqs,OM1_1_25_evi_proteins))
pepannot<-data.frame(sequence=character(),proteins=character())

for (n in 1:nrow(OM1_1_25_evi)) {
  keyin<-OM1_1_25_evi$Sequence[n]
  Ttsub<-Tt[Tt$OM1_1_25_evi_seqs==keyin,]
  prots<-Ttsub$OM1_1_25_evi_proteins
  protssublist<-c()
  for (q in 1:length(prots)) {
    protssub<-prots[q]
    protssublist<-c(protssublist,unlist(strsplit(protssub, ";")))
    
    
  }
  protslist<-unique(protssublist)
  pepannot[n,]<-c(as.character(keyin),paste(protslist,sep = "",collapse = ";"))
  
  
}
pepannot<-unique(pepannot)


for (n in 1:nrow(OM1_1_25)) {
  keyin<-OM1_1_25$Sequence[n]
  protsin<-pepannot$proteins[pepannot$sequence==keyin]
  OM1_1_25$Proteins[n]<-protsin
  
}


rm(Tt,pepannot,keyin,Ttsub,prots,protssublist,protslist)


OM1_2_10<-OM1_2_10[!is.na(OM1_2_10$FractionOxidized),]
OM1_2_10<-OM1_2_10[OM1_2_10$Fit>=0.8,]
OM1_2_10_evi<-read.table("C://Users/jbett/Desktop/mouseMOBB/newresults/evidenceOM1_2_10.txt",sep="\t",header = TRUE)
OM1_2_10_evi_proteins<-OM1_2_10_evi$Proteins
OM1_2_10_evi_seqs<-OM1_2_10_evi$Sequence

Tt<-as.data.frame(cbind(OM1_2_10_evi_seqs,OM1_2_10_evi_proteins))
pepannot<-data.frame(sequence=character(),proteins=character())

for (n in 1:nrow(OM1_2_10_evi)) {
  keyin<-OM1_2_10_evi$Sequence[n]
  Ttsub<-Tt[Tt$OM1_2_10_evi_seqs==keyin,]
  prots<-Ttsub$OM1_2_10_evi_proteins
  protssublist<-c()
  for (q in 1:length(prots)) {
    protssub<-prots[q]
    protssublist<-c(protssublist,unlist(strsplit(protssub, ";")))
    
    
  }
  protslist<-unique(protssublist)
  pepannot[n,]<-c(as.character(keyin),paste(protslist,sep = "",collapse = ";"))
  
  
}
pepannot<-unique(pepannot)


for (n in 1:nrow(OM1_2_10)) {
  keyin<-OM1_2_10$Sequence[n]
  protsin<-pepannot$proteins[pepannot$sequence==keyin]
  OM1_2_10$Proteins[n]<-protsin
  
}


rm(Tt,pepannot,keyin,Ttsub,prots,protssublist,protslist)

OM1_2_25<-OM1_2_25[!is.na(OM1_2_25$FractionOxidized),]
OM1_2_25<-OM1_2_25[OM1_2_25$Fit>=0.8,]
OM1_2_25_evi<-read.table("C://Users/jbett/Desktop/mouseMOBB/newresults/evidenceOM1_2_25.txt",sep="\t",header = TRUE)
OM1_2_25_evi_proteins<-OM1_2_25_evi$Proteins
OM1_2_25_evi_seqs<-OM1_2_25_evi$Sequence

Tt<-as.data.frame(cbind(OM1_2_25_evi_seqs,OM1_2_25_evi_proteins))
pepannot<-data.frame(sequence=character(),proteins=character())

for (n in 1:nrow(OM1_2_25_evi)) {
  keyin<-OM1_2_25_evi$Sequence[n]
  Ttsub<-Tt[Tt$OM1_2_25_evi_seqs==keyin,]
  prots<-Ttsub$OM1_2_25_evi_proteins
  protssublist<-c()
  for (q in 1:length(prots)) {
    protssub<-prots[q]
    protssublist<-c(protssublist,unlist(strsplit(protssub, ";")))
    
    
  }
  protslist<-unique(protssublist)
  pepannot[n,]<-c(as.character(keyin),paste(protslist,sep = "",collapse = ";"))
  
  
}
pepannot<-unique(pepannot)


for (n in 1:nrow(OM1_2_25)) {
  keyin<-OM1_2_25$Sequence[n]
  protsin<-pepannot$proteins[pepannot$sequence==keyin]
  OM1_2_25$Proteins[n]<-protsin
  
}


rm(Tt,pepannot,keyin,Ttsub,prots,protssublist,protslist)

OM2_1_10<-OM2_1_10[!is.na(OM2_1_10$FractionOxidized),]
OM2_1_10<-OM2_1_10[OM2_1_10$Fit>=0.8,]
OM2_1_10_evi<-read.table("C://Users/jbett/Desktop/mouseMOBB/newresults/evidenceOM2_1_10.txt",sep="\t",header = TRUE)
OM2_1_10_evi_proteins<-OM2_1_10_evi$Proteins
OM2_1_10_evi_seqs<-OM2_1_10_evi$Sequence

Tt<-as.data.frame(cbind(OM2_1_10_evi_seqs,OM2_1_10_evi_proteins))
pepannot<-data.frame(sequence=character(),proteins=character())

for (n in 1:nrow(OM2_1_10_evi)) {
  keyin<-OM2_1_10_evi$Sequence[n]
  Ttsub<-Tt[Tt$OM2_1_10_evi_seqs==keyin,]
  prots<-Ttsub$OM2_1_10_evi_proteins
  protssublist<-c()
  for (q in 1:length(prots)) {
    protssub<-prots[q]
    protssublist<-c(protssublist,unlist(strsplit(protssub, ";")))
    
    
  }
  protslist<-unique(protssublist)
  pepannot[n,]<-c(as.character(keyin),paste(protslist,sep = "",collapse = ";"))
  
  
}
pepannot<-unique(pepannot)


for (n in 1:nrow(OM2_1_10)) {
  keyin<-OM2_1_10$Sequence[n]
  protsin<-pepannot$proteins[pepannot$sequence==keyin]
  OM2_1_10$Proteins[n]<-protsin
  
}


rm(Tt,pepannot,keyin,Ttsub,prots,protssublist,protslist)


OM2_1_25<-OM2_1_25[!is.na(OM2_1_25$FractionOxidized),]
OM2_1_25<-OM2_1_25[OM2_1_25$Fit>=0.8,]
OM2_1_25_evi<-read.table("C://Users/jbett/Desktop/mouseMOBB/newresults/evidenceOM2_1_25.txt",sep="\t",header = TRUE)
OM2_1_25_evi_proteins<-OM2_1_25_evi$Proteins
OM2_1_25_evi_seqs<-OM2_1_25_evi$Sequence

Tt<-as.data.frame(cbind(OM2_1_25_evi_seqs,OM2_1_25_evi_proteins))
pepannot<-data.frame(sequence=character(),proteins=character())

for (n in 1:nrow(OM2_1_25_evi)) {
  keyin<-OM2_1_25_evi$Sequence[n]
  Ttsub<-Tt[Tt$OM2_1_25_evi_seqs==keyin,]
  prots<-Ttsub$OM2_1_25_evi_proteins
  protssublist<-c()
  for (q in 1:length(prots)) {
    protssub<-prots[q]
    protssublist<-c(protssublist,unlist(strsplit(protssub, ";")))
    
    
  }
  protslist<-unique(protssublist)
  pepannot[n,]<-c(as.character(keyin),paste(protslist,sep = "",collapse = ";"))
  
  
}
pepannot<-unique(pepannot)


for (n in 1:nrow(OM2_1_25)) {
  keyin<-OM2_1_25$Sequence[n]
  protsin<-pepannot$proteins[pepannot$sequence==keyin]
  OM2_1_25$Proteins[n]<-protsin
  
}


rm(Tt,pepannot,keyin,Ttsub,prots,protssublist,protslist)


OM2_2_10<-OM2_2_10[!is.na(OM2_2_10$FractionOxidized),]
OM2_2_10<-OM2_2_10[OM2_2_10$Fit>=0.8,]
OM2_2_10_evi<-read.table("C://Users/jbett/Desktop/mouseMOBB/newresults/evidenceOM2_2_10.txt",sep="\t",header = TRUE)
OM2_2_10_evi_proteins<-OM2_2_10_evi$Proteins
OM2_2_10_evi_seqs<-OM2_2_10_evi$Sequence

Tt<-as.data.frame(cbind(OM2_2_10_evi_seqs,OM2_2_10_evi_proteins))
pepannot<-data.frame(sequence=character(),proteins=character())

for (n in 1:nrow(OM2_2_10_evi)) {
  keyin<-OM2_2_10_evi$Sequence[n]
  Ttsub<-Tt[Tt$OM2_2_10_evi_seqs==keyin,]
  prots<-Ttsub$OM2_2_10_evi_proteins
  protssublist<-c()
  for (q in 1:length(prots)) {
    protssub<-prots[q]
    protssublist<-c(protssublist,unlist(strsplit(protssub, ";")))
    
    
  }
  protslist<-unique(protssublist)
  pepannot[n,]<-c(as.character(keyin),paste(protslist,sep = "",collapse = ";"))
  
  
}
pepannot<-unique(pepannot)


for (n in 1:nrow(OM2_2_10)) {
  keyin<-OM2_2_10$Sequence[n]
  protsin<-pepannot$proteins[pepannot$sequence==keyin]
  OM2_2_10$Proteins[n]<-protsin
  
}


rm(Tt,pepannot,keyin,Ttsub,prots,protssublist,protslist)


OM2_2_25<-OM2_2_25[!is.na(OM2_2_25$FractionOxidized),]
OM2_2_25<-OM2_2_25[OM2_2_25$Fit>=0.8,]
OM2_2_25_evi<-read.table("C://Users/jbett/Desktop/mouseMOBB/newresults/evidenceOM2_2_25.txt",sep="\t",header = TRUE)
OM2_2_25_evi_proteins<-OM2_2_25_evi$Proteins
OM2_2_25_evi_seqs<-OM2_2_25_evi$Sequence

Tt<-as.data.frame(cbind(OM2_2_25_evi_seqs,OM2_2_25_evi_proteins))
pepannot<-data.frame(sequence=character(),proteins=character())

for (n in 1:nrow(OM2_2_25_evi)) {
  keyin<-OM2_2_25_evi$Sequence[n]
  Ttsub<-Tt[Tt$OM2_2_25_evi_seqs==keyin,]
  prots<-Ttsub$OM2_2_25_evi_proteins
  protssublist<-c()
  for (q in 1:length(prots)) {
    protssub<-prots[q]
    protssublist<-c(protssublist,unlist(strsplit(protssub, ";")))
    
    
  }
  protslist<-unique(protssublist)
  pepannot[n,]<-c(as.character(keyin),paste(protslist,sep = "",collapse = ";"))
  
  
}
pepannot<-unique(pepannot)


for (n in 1:nrow(OM2_2_25)) {
  keyin<-OM2_2_25$Sequence[n]
  protsin<-pepannot$proteins[pepannot$sequence==keyin]
  OM2_2_25$Proteins[n]<-protsin
  
}


rm(Tt,pepannot,keyin,Ttsub,prots,protssublist,protslist)


OM3_1_10<-OM3_1_10[!is.na(OM3_1_10$FractionOxidized),]
OM3_1_10<-OM3_1_10[OM3_1_10$Fit>=0.8,]
OM3_1_10_evi<-read.table("C://Users/jbett/Desktop/mouseMOBB/newresults/evidenceOM3_1_10.txt",sep="\t",header = TRUE)
OM3_1_10_evi_proteins<-OM3_1_10_evi$Proteins
OM3_1_10_evi_seqs<-OM3_1_10_evi$Sequence

Tt<-as.data.frame(cbind(OM3_1_10_evi_seqs,OM3_1_10_evi_proteins))
pepannot<-data.frame(sequence=character(),proteins=character())

for (n in 1:nrow(OM3_1_10_evi)) {
  keyin<-OM3_1_10_evi$Sequence[n]
  Ttsub<-Tt[Tt$OM3_1_10_evi_seqs==keyin,]
  prots<-Ttsub$OM3_1_10_evi_proteins
  protssublist<-c()
  for (q in 1:length(prots)) {
    protssub<-prots[q]
    protssublist<-c(protssublist,unlist(strsplit(protssub, ";")))
    
    
  }
  protslist<-unique(protssublist)
  pepannot[n,]<-c(as.character(keyin),paste(protslist,sep = "",collapse = ";"))
  
  
}
pepannot<-unique(pepannot)


for (n in 1:nrow(OM3_1_10)) {
  keyin<-OM3_1_10$Sequence[n]
  protsin<-pepannot$proteins[pepannot$sequence==keyin]
  OM3_1_10$Proteins[n]<-protsin
  
}


rm(Tt,pepannot,keyin,Ttsub,prots,protssublist,protslist)


OM3_1_25<-OM3_1_25[!is.na(OM3_1_25$FractionOxidized),]
OM3_1_25<-OM3_1_25[OM3_1_25$Fit>=0.8,]
OM3_1_25_evi<-read.table("C://Users/jbett/Desktop/mouseMOBB/newresults/evidenceOM3_1_25.txt",sep="\t",header = TRUE)
OM3_1_25_evi_proteins<-OM3_1_25_evi$Proteins
OM3_1_25_evi_seqs<-OM3_1_25_evi$Sequence

Tt<-as.data.frame(cbind(OM3_1_25_evi_seqs,OM3_1_25_evi_proteins))
pepannot<-data.frame(sequence=character(),proteins=character())

for (n in 1:nrow(OM3_1_25_evi)) {
  keyin<-OM3_1_25_evi$Sequence[n]
  Ttsub<-Tt[Tt$OM3_1_25_evi_seqs==keyin,]
  prots<-Ttsub$OM3_1_25_evi_proteins
  protssublist<-c()
  for (q in 1:length(prots)) {
    protssub<-prots[q]
    protssublist<-c(protssublist,unlist(strsplit(protssub, ";")))
    
    
  }
  protslist<-unique(protssublist)
  pepannot[n,]<-c(as.character(keyin),paste(protslist,sep = "",collapse = ";"))
  
  
}
pepannot<-unique(pepannot)


for (n in 1:nrow(OM3_1_25)) {
  keyin<-OM3_1_25$Sequence[n]
  protsin<-pepannot$proteins[pepannot$sequence==keyin]
  OM3_1_25$Proteins[n]<-protsin
  
}


rm(Tt,pepannot,keyin,Ttsub,prots,protssublist,protslist)


OM3_2_10<-OM3_2_10[!is.na(OM3_2_10$FractionOxidized),]
OM3_2_10<-OM3_2_10[OM3_2_10$Fit>=0.8,]
OM3_2_10_evi<-read.table("C://Users/jbett/Desktop/mouseMOBB/newresults/evidenceOM3_2_10.txt",sep="\t",header = TRUE)
OM3_2_10_evi_proteins<-OM3_2_10_evi$Proteins
OM3_2_10_evi_seqs<-OM3_2_10_evi$Sequence

Tt<-as.data.frame(cbind(OM3_2_10_evi_seqs,OM3_2_10_evi_proteins))
pepannot<-data.frame(sequence=character(),proteins=character())

for (n in 1:nrow(OM3_2_10_evi)) {
  keyin<-OM3_2_10_evi$Sequence[n]
  Ttsub<-Tt[Tt$OM3_2_10_evi_seqs==keyin,]
  prots<-Ttsub$OM3_2_10_evi_proteins
  protssublist<-c()
  for (q in 1:length(prots)) {
    protssub<-prots[q]
    protssublist<-c(protssublist,unlist(strsplit(protssub, ";")))
    
    
  }
  protslist<-unique(protssublist)
  pepannot[n,]<-c(as.character(keyin),paste(protslist,sep = "",collapse = ";"))
  
  
}
pepannot<-unique(pepannot)


for (n in 1:nrow(OM3_2_10)) {
  keyin<-OM3_2_10$Sequence[n]
  protsin<-pepannot$proteins[pepannot$sequence==keyin]
  OM3_2_10$Proteins[n]<-protsin
  
}


rm(Tt,pepannot,keyin,Ttsub,prots,protssublist,protslist)


OM3_2_25<-OM3_2_25[!is.na(OM3_2_25$FractionOxidized),]
OM3_2_25<-OM3_2_25[OM3_2_25$Fit>=0.8,]
OM3_2_25_evi<-read.table("C://Users/jbett/Desktop/mouseMOBB/newresults/evidenceOM3_2_25.txt",sep="\t",header = TRUE)
OM3_2_25_evi_proteins<-OM3_2_25_evi$Proteins
OM3_2_25_evi_seqs<-OM3_2_25_evi$Sequence

Tt<-as.data.frame(cbind(OM3_2_25_evi_seqs,OM3_2_25_evi_proteins))
pepannot<-data.frame(sequence=character(),proteins=character())

for (n in 1:nrow(OM3_2_25_evi)) {
  keyin<-OM3_2_25_evi$Sequence[n]
  Ttsub<-Tt[Tt$OM3_2_25_evi_seqs==keyin,]
  prots<-Ttsub$OM3_2_25_evi_proteins
  protssublist<-c()
  for (q in 1:length(prots)) {
    protssub<-prots[q]
    protssublist<-c(protssublist,unlist(strsplit(protssub, ";")))
    
    
  }
  protslist<-unique(protssublist)
  pepannot[n,]<-c(as.character(keyin),paste(protslist,sep = "",collapse = ";"))
  
  
}
pepannot<-unique(pepannot)


for (n in 1:nrow(OM3_2_25)) {
  keyin<-OM3_2_25$Sequence[n]
  protsin<-pepannot$proteins[pepannot$sequence==keyin]
  OM3_2_25$Proteins[n]<-protsin
  
}


rm(Tt,pepannot,keyin,Ttsub,prots,protssublist,protslist)


OM1_1_10$Fit<-NULL
OM1_1_25$Fit<-NULL
OM1_2_10$Fit<-NULL
OM1_2_25$Fit<-NULL
OM2_1_10$Fit<-NULL
OM2_1_25$Fit<-NULL
OM2_2_10$Fit<-NULL
OM2_2_25$Fit<-NULL
OM3_1_10$Fit<-NULL
OM3_1_25$Fit<-NULL
OM3_2_10$Fit<-NULL
OM3_2_25$Fit<-NULL


df_OM1_1<-merge(OM1_1_10,OM1_1_25,by=c("Sequence"),all=TRUE)
for (n in 1:nrow(df_OM1_1)) {
  ifelse(is.na(df_OM1_1$Proteins.x[n]),df_OM1_1$Proteins.x[n]<-df_OM1_1$Proteins.y[n],
         df_OM1_1$Proteins.x[n]<-df_OM1_1$Proteins.x[n])
  
}
df_OM1_1$Proteins.y<-NULL
colnames(df_OM1_1)<-c("Sequence","Fraction_0xidized_OM1_1_10","Proteins","Fraction_0xidized_OM1_1_25")
#write.csv(df_OM1_1,"C://Users/jbett/Desktop/mouseMOBB/newresults/results/mergedtitrationpoints/OM1_1.csv",row.names = FALSE,col.names = TRUE)

df_OM1_2<-merge(OM1_2_10,OM1_2_25,by=c("Sequence"),all=TRUE)
for (n in 1:nrow(df_OM1_2)) {
  ifelse(is.na(df_OM1_2$Proteins.x[n]),df_OM1_2$Proteins.x[n]<-df_OM1_2$Proteins.y[n],
         df_OM1_2$Proteins.x[n]<-df_OM1_2$Proteins.x[n])
  
}
df_OM1_2$Proteins.y<-NULL
colnames(df_OM1_2)<-c("Sequence","Fraction_0xidized_OM1_2_10","Proteins","Fraction_0xidized_OM1_2_25")
#write.csv(df_OM1_2,"C://Users/jbett/Desktop/mouseMOBB/newresults/results/mergedtitrationpoints/OM1_2.csv",row.names = FALSE,col.names = TRUE)


df_OM2_1<-merge(OM2_1_10,OM2_1_25,by=c("Sequence"),all=TRUE)
for (n in 1:nrow(df_OM2_1)) {
  ifelse(is.na(df_OM2_1$Proteins.x[n]),df_OM2_1$Proteins.x[n]<-df_OM2_1$Proteins.y[n],
         df_OM2_1$Proteins.x[n]<-df_OM2_1$Proteins.x[n])
  
}
df_OM2_1$Proteins.y<-NULL
colnames(df_OM2_1)<-c("Sequence","Fraction_0xidized_OM2_1_10","Proteins","Fraction_0xidized_OM2_1_25")
#write.csv(df_OM2_1,"C://Users/jbett/Desktop/mouseMOBB/newresults/results/mergedtitrationpoints/OM2_1.csv",row.names = FALSE,col.names = TRUE)

df_OM2_2<-merge(OM2_2_10,OM2_2_25,by=c("Sequence"),all=TRUE)
for (n in 1:nrow(df_OM2_2)) {
  ifelse(is.na(df_OM2_2$Proteins.x[n]),df_OM2_2$Proteins.x[n]<-df_OM2_2$Proteins.y[n],
         df_OM2_2$Proteins.x[n]<-df_OM2_2$Proteins.x[n])
  
}
df_OM2_2$Proteins.y<-NULL
colnames(df_OM2_2)<-c("Sequence","Fraction_0xidized_OM2_2_10","Proteins","Fraction_0xidized_OM2_2_25")
#write.csv(df_OM2_2,"C://Users/jbett/Desktop/mouseMOBB/newresults/results/mergedtitrationpoints/OM2_2.csv",row.names = FALSE,col.names = TRUE)

df_OM3_1<-merge(OM3_1_10,OM3_1_25,by=c("Sequence"),all=TRUE)
for (n in 1:nrow(df_OM3_1)) {
  ifelse(is.na(df_OM3_1$Proteins.x[n]),df_OM3_1$Proteins.x[n]<-df_OM3_1$Proteins.y[n],
         df_OM3_1$Proteins.x[n]<-df_OM3_1$Proteins.x[n])
  
}
df_OM3_1$Proteins.y<-NULL
colnames(df_OM3_1)<-c("Sequence","Fraction_0xidized_OM3_1_10","Proteins","Fraction_0xidized_OM3_1_25")
#write.csv(df_OM3_1,"C://Users/jbett/Desktop/mouseMOBB/newresults/results/mergedtitrationpoints/OM3_1.csv",row.names = FALSE,col.names = TRUE)

df_OM3_2<-merge(OM3_2_10,OM3_2_25,by=c("Sequence"),all=TRUE)
for (n in 1:nrow(df_OM3_2)) {
  ifelse(is.na(df_OM3_2$Proteins.x[n]),df_OM3_2$Proteins.x[n]<-df_OM3_2$Proteins.y[n],
         df_OM3_2$Proteins.x[n]<-df_OM3_2$Proteins.x[n])
  
}
df_OM3_2$Proteins.y<-NULL
colnames(df_OM3_2)<-c("Sequence","Fraction_0xidized_OM3_2_10","Proteins","Fraction_0xidized_OM3_2_25")
#write.csv(df_OM3_2,"C://Users/jbett/Desktop/mouseMOBB/newresults/results/mergedtitrationpoints/OM3_2.csv",row.names = FALSE,col.names = TRUE)

dfold<-merge(df_OM1_1,df_OM1_2,by=c("Sequence","Proteins"),all=TRUE)
dfold<-merge(dfold,df_OM2_1,by=c("Sequence","Proteins"),all=TRUE)
dfold<-merge(dfold,df_OM2_2,by=c("Sequence","Proteins"),all=TRUE)
dfold<-merge(dfold,df_OM3_1,by=c("Sequence","Proteins"),all=TRUE)
dfold<-merge(dfold,df_OM3_2,by=c("Sequence","Proteins"),all=TRUE)
dfold<-dfold[!is.na(dfold$Sequence),]
dfold$validvalues<-12-rowSums(is.na(dfold[3:14]))
dfold<-dfold[dfold$validvalues>=7,]

################################################################################
################################################################################
#############Read in and Format young data######################################
################################################################################
################################################################################
################################################################################
################################################################################


YM1_1_10<-read.csv("C://Users/jbett/Desktop/mouseMOBB/newresults/results/YM1_1_10.txt",header=TRUE)
YM1_1_25<-read.csv("C://Users/jbett/Desktop/mouseMOBB/newresults/results/YM1_1_25.txt",header=TRUE)
YM1_2_10<-read.csv("C://Users/jbett/Desktop/mouseMOBB/newresults/results/YM1_2_10.txt",header=TRUE)
YM1_2_25<-read.csv("C://Users/jbett/Desktop/mouseMOBB/newresults/results/YM1_2_25.txt",header=TRUE)
YM2_1_10<-read.csv("C://Users/jbett/Desktop/mouseMOBB/newresults/results/YM2_1_10.txt",header=TRUE)
YM2_1_25<-read.csv("C://Users/jbett/Desktop/mouseMOBB/newresults/results/YM2_1_25.txt",header=TRUE)
YM2_2_10<-read.csv("C://Users/jbett/Desktop/mouseMOBB/newresults/results/YM2_2_10.txt",header=TRUE)
YM2_2_25<-read.csv("C://Users/jbett/Desktop/mouseMOBB/newresults/results/YM2_2_25.txt",header=TRUE)
YM3_1_10<-read.csv("C://Users/jbett/Desktop/mouseMOBB/newresults/results/YM3_1_10.txt",header=TRUE)
YM3_1_25<-read.csv("C://Users/jbett/Desktop/mouseMOBB/newresults/results/YM3_1_25.txt",header=TRUE)
YM3_2_10<-read.csv("C://Users/jbett/Desktop/mouseMOBB/newresults/results/YM3_2_10.txt",header=TRUE)
YM3_2_25<-read.csv("C://Users/jbett/Desktop/mouseMOBB/newresults/results/YM3_2_25.txt",header=TRUE)

YM1_1_10$X<-NULL
YM1_1_25$X<-NULL
YM1_2_10$X<-NULL
YM1_2_25$X<-NULL
YM2_1_10$X<-NULL
YM2_1_25$X<-NULL
YM2_2_10$X<-NULL
YM2_2_25$X<-NULL
YM3_1_10$X<-NULL
YM3_1_25$X<-NULL
YM3_2_10$X<-NULL
YM3_2_25$X<-NULL

YM1_1_10<-YM1_1_10[!is.na(YM1_1_10$FractionOxidized),]
YM1_1_10<-YM1_1_10[YM1_1_10$Fit>=0.8,]
YM1_1_10_evi<-read.table("C://Users/jbett/Desktop/mouseMOBB/newresults/evidenceYM1_1_10.txt",sep="\t",header = TRUE)
YM1_1_10_evi_proteins<-YM1_1_10_evi$Proteins
YM1_1_10_evi_seqs<-YM1_1_10_evi$Sequence

Tt<-as.data.frame(cbind(YM1_1_10_evi_seqs,YM1_1_10_evi_proteins))
pepannot<-data.frame(sequence=character(),proteins=character())

for (n in 1:nrow(YM1_1_10_evi)) {
  keyin<-YM1_1_10_evi$Sequence[n]
  Ttsub<-Tt[Tt$YM1_1_10_evi_seqs==keyin,]
  prots<-Ttsub$YM1_1_10_evi_proteins
  protssublist<-c()
  for (q in 1:length(prots)) {
    protssub<-prots[q]
    protssublist<-c(protssublist,unlist(strsplit(protssub, ";")))
    
    
  }
  protslist<-unique(protssublist)
  pepannot[n,]<-c(as.character(keyin),paste(protslist,sep = "",collapse = ";"))
  
  
}
pepannot<-unique(pepannot)


for (n in 1:nrow(YM1_1_10)) {
  keyin<-YM1_1_10$Sequence[n]
  protsin<-pepannot$proteins[pepannot$sequence==keyin]
  YM1_1_10$Proteins[n]<-protsin
  
}


rm(Tt,pepannot,keyin,Ttsub,prots,protssublist,protslist)

YM1_1_25<-YM1_1_25[!is.na(YM1_1_25$FractionOxidized),]
YM1_1_25<-YM1_1_25[YM1_1_25$Fit>=0.8,]
YM1_1_25_evi<-read.table("C://Users/jbett/Desktop/mouseMOBB/newresults/evidenceYM1_1_25.txt",sep="\t",header = TRUE)
YM1_1_25_evi_proteins<-YM1_1_25_evi$Proteins
YM1_1_25_evi_seqs<-YM1_1_25_evi$Sequence

Tt<-as.data.frame(cbind(YM1_1_25_evi_seqs,YM1_1_25_evi_proteins))
pepannot<-data.frame(sequence=character(),proteins=character())

for (n in 1:nrow(YM1_1_25_evi)) {
  keyin<-YM1_1_25_evi$Sequence[n]
  Ttsub<-Tt[Tt$YM1_1_25_evi_seqs==keyin,]
  prots<-Ttsub$YM1_1_25_evi_proteins
  protssublist<-c()
  for (q in 1:length(prots)) {
    protssub<-prots[q]
    protssublist<-c(protssublist,unlist(strsplit(protssub, ";")))
    
    
  }
  protslist<-unique(protssublist)
  pepannot[n,]<-c(as.character(keyin),paste(protslist,sep = "",collapse = ";"))
  
  
}
pepannot<-unique(pepannot)


for (n in 1:nrow(YM1_1_25)) {
  keyin<-YM1_1_25$Sequence[n]
  protsin<-pepannot$proteins[pepannot$sequence==keyin]
  YM1_1_25$Proteins[n]<-protsin
  
}


rm(Tt,pepannot,keyin,Ttsub,prots,protssublist,protslist)


YM1_2_10<-YM1_2_10[!is.na(YM1_2_10$FractionOxidized),]
YM1_2_10<-YM1_2_10[YM1_2_10$Fit>=0.8,]
YM1_2_10_evi<-read.table("C://Users/jbett/Desktop/mouseMOBB/newresults/evidenceYM1_2_10.txt",sep="\t",header = TRUE)
YM1_2_10_evi_proteins<-YM1_2_10_evi$Proteins
YM1_2_10_evi_seqs<-YM1_2_10_evi$Sequence

Tt<-as.data.frame(cbind(YM1_2_10_evi_seqs,YM1_2_10_evi_proteins))
pepannot<-data.frame(sequence=character(),proteins=character())

for (n in 1:nrow(YM1_2_10_evi)) {
  keyin<-YM1_2_10_evi$Sequence[n]
  Ttsub<-Tt[Tt$YM1_2_10_evi_seqs==keyin,]
  prots<-Ttsub$YM1_2_10_evi_proteins
  protssublist<-c()
  for (q in 1:length(prots)) {
    protssub<-prots[q]
    protssublist<-c(protssublist,unlist(strsplit(protssub, ";")))
    
    
  }
  protslist<-unique(protssublist)
  pepannot[n,]<-c(as.character(keyin),paste(protslist,sep = "",collapse = ";"))
  
  
}
pepannot<-unique(pepannot)


for (n in 1:nrow(YM1_2_10)) {
  keyin<-YM1_2_10$Sequence[n]
  protsin<-pepannot$proteins[pepannot$sequence==keyin]
  YM1_2_10$Proteins[n]<-protsin
  
}


rm(Tt,pepannot,keyin,Ttsub,prots,protssublist,protslist)

YM1_2_25<-YM1_2_25[!is.na(YM1_2_25$FractionOxidized),]
YM1_2_25<-YM1_2_25[YM1_2_25$Fit>=0.8,]
YM1_2_25_evi<-read.table("C://Users/jbett/Desktop/mouseMOBB/newresults/evidenceYM1_2_25.txt",sep="\t",header = TRUE)
YM1_2_25_evi_proteins<-YM1_2_25_evi$Proteins
YM1_2_25_evi_seqs<-YM1_2_25_evi$Sequence

Tt<-as.data.frame(cbind(YM1_2_25_evi_seqs,YM1_2_25_evi_proteins))
pepannot<-data.frame(sequence=character(),proteins=character())

for (n in 1:nrow(YM1_2_25_evi)) {
  keyin<-YM1_2_25_evi$Sequence[n]
  Ttsub<-Tt[Tt$YM1_2_25_evi_seqs==keyin,]
  prots<-Ttsub$YM1_2_25_evi_proteins
  protssublist<-c()
  for (q in 1:length(prots)) {
    protssub<-prots[q]
    protssublist<-c(protssublist,unlist(strsplit(protssub, ";")))
    
    
  }
  protslist<-unique(protssublist)
  pepannot[n,]<-c(as.character(keyin),paste(protslist,sep = "",collapse = ";"))
  
  
}
pepannot<-unique(pepannot)


for (n in 1:nrow(YM1_2_25)) {
  keyin<-YM1_2_25$Sequence[n]
  protsin<-pepannot$proteins[pepannot$sequence==keyin]
  YM1_2_25$Proteins[n]<-protsin
  
}


rm(Tt,pepannot,keyin,Ttsub,prots,protssublist,protslist)

YM2_1_10<-YM2_1_10[!is.na(YM2_1_10$FractionOxidized),]
YM2_1_10<-YM2_1_10[YM2_1_10$Fit>=0.8,]
YM2_1_10_evi<-read.table("C://Users/jbett/Desktop/mouseMOBB/newresults/evidenceYM2_1_10.txt",sep="\t",header = TRUE)
YM2_1_10_evi_proteins<-YM2_1_10_evi$Proteins
YM2_1_10_evi_seqs<-YM2_1_10_evi$Sequence

Tt<-as.data.frame(cbind(YM2_1_10_evi_seqs,YM2_1_10_evi_proteins))
pepannot<-data.frame(sequence=character(),proteins=character())

for (n in 1:nrow(YM2_1_10_evi)) {
  keyin<-YM2_1_10_evi$Sequence[n]
  Ttsub<-Tt[Tt$YM2_1_10_evi_seqs==keyin,]
  prots<-Ttsub$YM2_1_10_evi_proteins
  protssublist<-c()
  for (q in 1:length(prots)) {
    protssub<-prots[q]
    protssublist<-c(protssublist,unlist(strsplit(protssub, ";")))
    
    
  }
  protslist<-unique(protssublist)
  pepannot[n,]<-c(as.character(keyin),paste(protslist,sep = "",collapse = ";"))
  
  
}
pepannot<-unique(pepannot)


for (n in 1:nrow(YM2_1_10)) {
  keyin<-YM2_1_10$Sequence[n]
  protsin<-pepannot$proteins[pepannot$sequence==keyin]
  YM2_1_10$Proteins[n]<-protsin
  
}


rm(Tt,pepannot,keyin,Ttsub,prots,protssublist,protslist)


YM2_1_25<-YM2_1_25[!is.na(YM2_1_25$FractionOxidized),]
YM2_1_25<-YM2_1_25[YM2_1_25$Fit>=0.8,]
YM2_1_25_evi<-read.table("C://Users/jbett/Desktop/mouseMOBB/newresults/evidenceYM2_1_25.txt",sep="\t",header = TRUE)
YM2_1_25_evi_proteins<-YM2_1_25_evi$Proteins
YM2_1_25_evi_seqs<-YM2_1_25_evi$Sequence

Tt<-as.data.frame(cbind(YM2_1_25_evi_seqs,YM2_1_25_evi_proteins))
pepannot<-data.frame(sequence=character(),proteins=character())

for (n in 1:nrow(YM2_1_25_evi)) {
  keyin<-YM2_1_25_evi$Sequence[n]
  Ttsub<-Tt[Tt$YM2_1_25_evi_seqs==keyin,]
  prots<-Ttsub$YM2_1_25_evi_proteins
  protssublist<-c()
  for (q in 1:length(prots)) {
    protssub<-prots[q]
    protssublist<-c(protssublist,unlist(strsplit(protssub, ";")))
    
    
  }
  protslist<-unique(protssublist)
  pepannot[n,]<-c(as.character(keyin),paste(protslist,sep = "",collapse = ";"))
  
  
}
pepannot<-unique(pepannot)


for (n in 1:nrow(YM2_1_25)) {
  keyin<-YM2_1_25$Sequence[n]
  protsin<-pepannot$proteins[pepannot$sequence==keyin]
  YM2_1_25$Proteins[n]<-protsin
  
}


rm(Tt,pepannot,keyin,Ttsub,prots,protssublist,protslist)


YM2_2_10<-YM2_2_10[!is.na(YM2_2_10$FractionOxidized),]
YM2_2_10<-YM2_2_10[YM2_2_10$Fit>=0.8,]
YM2_2_10_evi<-read.table("C://Users/jbett/Desktop/mouseMOBB/newresults/evidenceYM2_2_10.txt",sep="\t",header = TRUE)
YM2_2_10_evi_proteins<-YM2_2_10_evi$Proteins
YM2_2_10_evi_seqs<-YM2_2_10_evi$Sequence

Tt<-as.data.frame(cbind(YM2_2_10_evi_seqs,YM2_2_10_evi_proteins))
pepannot<-data.frame(sequence=character(),proteins=character())

for (n in 1:nrow(YM2_2_10_evi)) {
  keyin<-YM2_2_10_evi$Sequence[n]
  Ttsub<-Tt[Tt$YM2_2_10_evi_seqs==keyin,]
  prots<-Ttsub$YM2_2_10_evi_proteins
  protssublist<-c()
  for (q in 1:length(prots)) {
    protssub<-prots[q]
    protssublist<-c(protssublist,unlist(strsplit(protssub, ";")))
    
    
  }
  protslist<-unique(protssublist)
  pepannot[n,]<-c(as.character(keyin),paste(protslist,sep = "",collapse = ";"))
  
  
}
pepannot<-unique(pepannot)


for (n in 1:nrow(YM2_2_10)) {
  keyin<-YM2_2_10$Sequence[n]
  protsin<-pepannot$proteins[pepannot$sequence==keyin]
  YM2_2_10$Proteins[n]<-protsin
  
}


rm(Tt,pepannot,keyin,Ttsub,prots,protssublist,protslist)


YM2_2_25<-YM2_2_25[!is.na(YM2_2_25$FractionOxidized),]
YM2_2_25<-YM2_2_25[YM2_2_25$Fit>=0.8,]
YM2_2_25_evi<-read.table("C://Users/jbett/Desktop/mouseMOBB/newresults/evidenceYM2_2_25.txt",sep="\t",header = TRUE)
YM2_2_25_evi_proteins<-YM2_2_25_evi$Proteins
YM2_2_25_evi_seqs<-YM2_2_25_evi$Sequence

Tt<-as.data.frame(cbind(YM2_2_25_evi_seqs,YM2_2_25_evi_proteins))
pepannot<-data.frame(sequence=character(),proteins=character())

for (n in 1:nrow(YM2_2_25_evi)) {
  keyin<-YM2_2_25_evi$Sequence[n]
  Ttsub<-Tt[Tt$YM2_2_25_evi_seqs==keyin,]
  prots<-Ttsub$YM2_2_25_evi_proteins
  protssublist<-c()
  for (q in 1:length(prots)) {
    protssub<-prots[q]
    protssublist<-c(protssublist,unlist(strsplit(protssub, ";")))
    
    
  }
  protslist<-unique(protssublist)
  pepannot[n,]<-c(as.character(keyin),paste(protslist,sep = "",collapse = ";"))
  
  
}
pepannot<-unique(pepannot)


for (n in 1:nrow(YM2_2_25)) {
  keyin<-YM2_2_25$Sequence[n]
  protsin<-pepannot$proteins[pepannot$sequence==keyin]
  YM2_2_25$Proteins[n]<-protsin
  
}


rm(Tt,pepannot,keyin,Ttsub,prots,protssublist,protslist)


YM3_1_10<-YM3_1_10[!is.na(YM3_1_10$FractionOxidized),]
YM3_1_10<-YM3_1_10[YM3_1_10$Fit>=0.8,]
YM3_1_10_evi<-read.table("C://Users/jbett/Desktop/mouseMOBB/newresults/evidenceYM3_1_10.txt",sep="\t",header = TRUE)
YM3_1_10_evi_proteins<-YM3_1_10_evi$Proteins
YM3_1_10_evi_seqs<-YM3_1_10_evi$Sequence

Tt<-as.data.frame(cbind(YM3_1_10_evi_seqs,YM3_1_10_evi_proteins))
pepannot<-data.frame(sequence=character(),proteins=character())

for (n in 1:nrow(YM3_1_10_evi)) {
  keyin<-YM3_1_10_evi$Sequence[n]
  Ttsub<-Tt[Tt$YM3_1_10_evi_seqs==keyin,]
  prots<-Ttsub$YM3_1_10_evi_proteins
  protssublist<-c()
  for (q in 1:length(prots)) {
    protssub<-prots[q]
    protssublist<-c(protssublist,unlist(strsplit(protssub, ";")))
    
    
  }
  protslist<-unique(protssublist)
  pepannot[n,]<-c(as.character(keyin),paste(protslist,sep = "",collapse = ";"))
  
  
}
pepannot<-unique(pepannot)


for (n in 1:nrow(YM3_1_10)) {
  keyin<-YM3_1_10$Sequence[n]
  protsin<-pepannot$proteins[pepannot$sequence==keyin]
  YM3_1_10$Proteins[n]<-protsin
  
}


rm(Tt,pepannot,keyin,Ttsub,prots,protssublist,protslist)


YM3_1_25<-YM3_1_25[!is.na(YM3_1_25$FractionOxidized),]
YM3_1_25<-YM3_1_25[YM3_1_25$Fit>=0.8,]
YM3_1_25_evi<-read.table("C://Users/jbett/Desktop/mouseMOBB/newresults/evidenceYM3_1_25.txt",sep="\t",header = TRUE)
YM3_1_25_evi_proteins<-YM3_1_25_evi$Proteins
YM3_1_25_evi_seqs<-YM3_1_25_evi$Sequence

Tt<-as.data.frame(cbind(YM3_1_25_evi_seqs,YM3_1_25_evi_proteins))
pepannot<-data.frame(sequence=character(),proteins=character())

for (n in 1:nrow(YM3_1_25_evi)) {
  keyin<-YM3_1_25_evi$Sequence[n]
  Ttsub<-Tt[Tt$YM3_1_25_evi_seqs==keyin,]
  prots<-Ttsub$YM3_1_25_evi_proteins
  protssublist<-c()
  for (q in 1:length(prots)) {
    protssub<-prots[q]
    protssublist<-c(protssublist,unlist(strsplit(protssub, ";")))
    
    
  }
  protslist<-unique(protssublist)
  pepannot[n,]<-c(as.character(keyin),paste(protslist,sep = "",collapse = ";"))
  
  
}
pepannot<-unique(pepannot)


for (n in 1:nrow(YM3_1_25)) {
  keyin<-YM3_1_25$Sequence[n]
  protsin<-pepannot$proteins[pepannot$sequence==keyin]
  YM3_1_25$Proteins[n]<-protsin
  
}


rm(Tt,pepannot,keyin,Ttsub,prots,protssublist,protslist)


YM3_2_10<-YM3_2_10[!is.na(YM3_2_10$FractionOxidized),]
YM3_2_10<-YM3_2_10[YM3_2_10$Fit>=0.8,]
YM3_2_10_evi<-read.table("C://Users/jbett/Desktop/mouseMOBB/newresults/evidenceYM3_2_10.txt",sep="\t",header = TRUE)
YM3_2_10_evi_proteins<-YM3_2_10_evi$Proteins
YM3_2_10_evi_seqs<-YM3_2_10_evi$Sequence

Tt<-as.data.frame(cbind(YM3_2_10_evi_seqs,YM3_2_10_evi_proteins))
pepannot<-data.frame(sequence=character(),proteins=character())

for (n in 1:nrow(YM3_2_10_evi)) {
  keyin<-YM3_2_10_evi$Sequence[n]
  Ttsub<-Tt[Tt$YM3_2_10_evi_seqs==keyin,]
  prots<-Ttsub$YM3_2_10_evi_proteins
  protssublist<-c()
  for (q in 1:length(prots)) {
    protssub<-prots[q]
    protssublist<-c(protssublist,unlist(strsplit(protssub, ";")))
    
    
  }
  protslist<-unique(protssublist)
  pepannot[n,]<-c(as.character(keyin),paste(protslist,sep = "",collapse = ";"))
  
  
}
pepannot<-unique(pepannot)


for (n in 1:nrow(YM3_2_10)) {
  keyin<-YM3_2_10$Sequence[n]
  protsin<-pepannot$proteins[pepannot$sequence==keyin]
  YM3_2_10$Proteins[n]<-protsin
  
}


rm(Tt,pepannot,keyin,Ttsub,prots,protssublist,protslist)


YM3_2_25<-YM3_2_25[!is.na(YM3_2_25$FractionOxidized),]
YM3_2_25<-YM3_2_25[YM3_2_25$Fit>=0.8,]
YM3_2_25_evi<-read.table("C://Users/jbett/Desktop/mouseMOBB/newresults/evidenceYM3_2_25.txt",sep="\t",header = TRUE)
YM3_2_25_evi_proteins<-YM3_2_25_evi$Proteins
YM3_2_25_evi_seqs<-YM3_2_25_evi$Sequence

Tt<-as.data.frame(cbind(YM3_2_25_evi_seqs,YM3_2_25_evi_proteins))
pepannot<-data.frame(sequence=character(),proteins=character())

for (n in 1:nrow(YM3_2_25_evi)) {
  keyin<-YM3_2_25_evi$Sequence[n]
  Ttsub<-Tt[Tt$YM3_2_25_evi_seqs==keyin,]
  prots<-Ttsub$YM3_2_25_evi_proteins
  protssublist<-c()
  for (q in 1:length(prots)) {
    protssub<-prots[q]
    protssublist<-c(protssublist,unlist(strsplit(protssub, ";")))
    
    
  }
  protslist<-unique(protssublist)
  pepannot[n,]<-c(as.character(keyin),paste(protslist,sep = "",collapse = ";"))
  
  
}
pepannot<-unique(pepannot)


for (n in 1:nrow(YM3_2_25)) {
  keyin<-YM3_2_25$Sequence[n]
  protsin<-pepannot$proteins[pepannot$sequence==keyin]
  YM3_2_25$Proteins[n]<-protsin
  
}


rm(Tt,pepannot,keyin,Ttsub,prots,protssublist,protslist)


YM1_1_10$Fit<-NULL
YM1_1_25$Fit<-NULL
YM1_2_10$Fit<-NULL
YM1_2_25$Fit<-NULL
YM2_1_10$Fit<-NULL
YM2_1_25$Fit<-NULL
YM2_2_10$Fit<-NULL
YM2_2_25$Fit<-NULL
YM3_1_10$Fit<-NULL
YM3_1_25$Fit<-NULL
YM3_2_10$Fit<-NULL
YM3_2_25$Fit<-NULL


df_YM1_1<-merge(YM1_1_10,YM1_1_25,by=c("Sequence"),all=TRUE)
for (n in 1:nrow(df_YM1_1)) {
  ifelse(is.na(df_YM1_1$Proteins.x[n]),df_YM1_1$Proteins.x[n]<-df_YM1_1$Proteins.y[n],
         df_YM1_1$Proteins.x[n]<-df_YM1_1$Proteins.x[n])
  
}
df_YM1_1$Proteins.y<-NULL
colnames(df_YM1_1)<-c("Sequence","Fraction_0xidized_YM1_1_10","Proteins","Fraction_0xidized_YM1_1_25")
#write.csv(df_YM1_1,"C://Users/jbett/Desktop/mouseMOBB/newresults/results/mergedtitrationpoints/YM1_1.csv",row.names = FALSE,col.names = TRUE)

df_YM1_2<-merge(YM1_2_10,YM1_2_25,by=c("Sequence"),all=TRUE)
for (n in 1:nrow(df_YM1_2)) {
  ifelse(is.na(df_YM1_2$Proteins.x[n]),df_YM1_2$Proteins.x[n]<-df_YM1_2$Proteins.y[n],
         df_YM1_2$Proteins.x[n]<-df_YM1_2$Proteins.x[n])
  
}
df_YM1_2$Proteins.y<-NULL
colnames(df_YM1_2)<-c("Sequence","Fraction_0xidized_YM1_2_10","Proteins","Fraction_0xidized_YM1_2_25")
#write.csv(df_YM1_2,"C://Users/jbett/Desktop/mouseMOBB/newresults/results/mergedtitrationpoints/YM1_2.csv",row.names = FALSE,col.names = TRUE)


df_YM2_1<-merge(YM2_1_10,YM2_1_25,by=c("Sequence"),all=TRUE)
for (n in 1:nrow(df_YM2_1)) {
  ifelse(is.na(df_YM2_1$Proteins.x[n]),df_YM2_1$Proteins.x[n]<-df_YM2_1$Proteins.y[n],
         df_YM2_1$Proteins.x[n]<-df_YM2_1$Proteins.x[n])
  
}
df_YM2_1$Proteins.y<-NULL
colnames(df_YM2_1)<-c("Sequence","Fraction_0xidized_YM2_1_10","Proteins","Fraction_0xidized_YM2_1_25")
#write.csv(df_YM2_1,"C://Users/jbett/Desktop/mouseMOBB/newresults/results/mergedtitrationpoints/YM2_1.csv",row.names = FALSE,col.names = TRUE)

df_YM2_2<-merge(YM2_2_10,YM2_2_25,by=c("Sequence"),all=TRUE)
for (n in 1:nrow(df_YM2_2)) {
  ifelse(is.na(df_YM2_2$Proteins.x[n]),df_YM2_2$Proteins.x[n]<-df_YM2_2$Proteins.y[n],
         df_YM2_2$Proteins.x[n]<-df_YM2_2$Proteins.x[n])
  
}
df_YM2_2$Proteins.y<-NULL
colnames(df_YM2_2)<-c("Sequence","Fraction_0xidized_YM2_2_10","Proteins","Fraction_0xidized_YM2_2_25")
#write.csv(df_YM2_2,"C://Users/jbett/Desktop/mouseMOBB/newresults/results/mergedtitrationpoints/YM2_2.csv",row.names = FALSE,col.names = TRUE)

df_YM3_1<-merge(YM3_1_10,YM3_1_25,by=c("Sequence"),all=TRUE)
for (n in 1:nrow(df_YM3_1)) {
  ifelse(is.na(df_YM3_1$Proteins.x[n]),df_YM3_1$Proteins.x[n]<-df_YM3_1$Proteins.y[n],
         df_YM3_1$Proteins.x[n]<-df_YM3_1$Proteins.x[n])
  
}
df_YM3_1$Proteins.y<-NULL
colnames(df_YM3_1)<-c("Sequence","Fraction_0xidized_YM3_1_10","Proteins","Fraction_0xidized_YM3_1_25")
#write.csv(df_YM3_1,"C://Users/jbett/Desktop/mouseMOBB/newresults/results/mergedtitrationpoints/YM3_1.csv",row.names = FALSE,col.names = TRUE)

df_YM3_2<-merge(YM3_2_10,YM3_2_25,by=c("Sequence"),all=TRUE)
for (n in 1:nrow(df_YM3_2)) {
  ifelse(is.na(df_YM3_2$Proteins.x[n]),df_YM3_2$Proteins.x[n]<-df_YM3_2$Proteins.y[n],
         df_YM3_2$Proteins.x[n]<-df_YM3_2$Proteins.x[n])
  
}
df_YM3_2$Proteins.y<-NULL
colnames(df_YM3_2)<-c("Sequence","Fraction_0xidized_YM3_2_10","Proteins","Fraction_0xidized_YM3_2_25")
#write.csv(df_YM3_2,"C://Users/jbett/Desktop/mouseMOBB/newresults/results/mergedtitrationpoints/YM3_2.csv",row.names = FALSE,col.names = TRUE)

dfyoung<-merge(df_YM1_1,df_YM1_2,by=c("Sequence","Proteins"),all=TRUE)
dfyoung<-merge(dfyoung,df_YM2_1,by=c("Sequence","Proteins"),all=TRUE)
dfyoung<-merge(dfyoung,df_YM2_2,by=c("Sequence","Proteins"),all=TRUE)
dfyoung<-merge(dfyoung,df_YM3_1,by=c("Sequence","Proteins"),all=TRUE)
dfyoung<-merge(dfyoung,df_YM3_2,by=c("Sequence","Proteins"),all=TRUE)
dfyoung<-dfyoung[!is.na(dfyoung$Sequence),]
dfyoung$validvalues<-12-rowSums(is.na(dfyoung[3:14]))
dfyoung<-dfyoung[dfyoung$validvalues>=7,]


################################################################################
################################################################################
############# Merge Young and Old Data #########################################
################################################################################
################################################################################
################################################################################
################################################################################


df<-merge(dfyoung,dfold,by=c("Sequence"))
for (n in 1:nrow(df)) {
  if(df$Proteins.x[n]!=df$Proteins.y[n]){
    xprots<-unlist(strsplit(df$Proteins.x[n], ";"))
    yprots<-unlist(strsplit(df$Proteins.y[57], ";"))
    ifelse(length(yprots)<length(xprots),df$Proteins.x[n]<-df$Proteins.y[n],df$Proteins.x[n]<-df$Proteins.x[n])
  }
  
}
df$Proteins.y<-NULL
validvalues_old<-df$validvalues.y
validvalues_young<-df$validvalues.x
df$validvalues.x<-NULL
df$validvalues.y<-NULL
df$validvalues_young<-validvalues_young
df$validvalues_old<-validvalues_old

colnames(df)[2]<-"Proteins"
df$validvalues<-df$validvalues_old+df$validvalues_young


################################################################################
################################################################################
############# Impute Missing Values  ###########################################
################################################################################
################################################################################
################################################################################
################################################################################
ks<-list(1,3,5,7,10,15,20)
kserrors_knn<-list()
kserrors_SVD<-list()
kserrors_lls<-list()
z=1

for (w in ks) {
  
  testdata<-stratified(df,"validvalues",0.5)
  nontestdata<-df[!(df$Sequence%in%testdata$Sequence),]
  
  test<-data.frame(matrix(nrow = nrow(testdata),ncol = 5))
  colnames(test)<-c("n","i","realvalue","imputedvalue","validvalues")
  
  
  for (n in 1:nrow(testdata)) {
    invector<-as.numeric(testdata[n,3:26])
    invector2<-invector[!is.na(invector)]
    r<-sample(invector2,1)
    i<-which(invector==r)
    test$realvalue[n]<-invector[i]
    testdata[n,i+2]<-NA
    test$n[n]<-testdata$Sequence[n]
    test$i[n]<-i+2
    test$validvalues[n]<-testdata$validvalues[n]
    
    
  }
  
  dft_knn<-as.data.frame(rbind(testdata,nontestdata))
  dft_knn[,3:26]<-knnImputation(dft_knn[,3:26],k=w)
  
  dft_SVD<-as.data.frame(rbind(testdata,nontestdata))
  datmat<-dft_SVD[,3:26]
  datmat<-pca(datmat, method="svdImpute", nPcs = w, center = TRUE)
  datmat<-completeObs(datmat)
  dft_SVD[,3:26]<-datmat
  rm(datmat)
  
  dft_lls<-as.data.frame(rbind(testdata,nontestdata))
  datmat<-dft_lls[,3:26]
  impute.lls<-llsImpute(t(datmat),k=w,allVariables = TRUE,center = TRUE)
  datmat<-t(completeObs(impute.lls))
  dft_lls[,3:26]<-datmat
  
  
  for (q in 1:nrow(test)) {
    
    rowin_knn<-dft_knn[dft_knn$Sequence==test$n[q],]
    test$imputedvalue_knn[q]<-as.numeric(rowin_knn[test$i[q]])
    
    rowin_SVD<-dft_SVD[dft_SVD$Sequence==test$n[q],]
    test$imputedvalue_SVD[q]<-as.numeric(rowin_SVD[test$i[q]])
    
    rowin_lls<-dft_lls[dft_lls$Sequence==test$n[q],]
    test$imputedvalue_lls[q]<-as.numeric(rowin_lls[test$i[q]])
    
  }
  
  kserrors_knn[z]<-cor(test$realvalue,test$imputedvalue_knn)
  kserrors_SVD[z]<-cor(test$realvalue,test$imputedvalue_SVD)
  kserrors_lls[z]<-cor(test$realvalue,test$imputedvalue_lls)
  z=z+1
}


#row means#
test$imputedvalue<-0
dft<-as.data.frame(rbind(testdata,nontestdata))
datmat<-dft[,3:26]
k <- which(is.na(datmat), arr.ind=TRUE)
datmat[k] <- rowMeans(datmat, na.rm=TRUE)[k[,1]]
dft[,3:26]<-datmat
for (q in 1:nrow(test)) {
  inrow<-dft[dft$Sequence==test$n[q],]
  test$imputedvalue[q]<-as.numeric(inrow[test$i[q]])
  
}
rowmeanserror<-cor(test$realvalue,test$imputedvalue)

plot(ks,kserrors_knn,ylim=c(-0.065,1),col="blue")
lines(ks,kserrors_knn,col="blue",lty=2)
points(ks,kserrors_SVD,col="green")
lines(ks,kserrors_SVD,col="green",lty=2)
points(ks,kserrors_lls,col="purple")
lines(ks,kserrors_lls,col="purple",lty=2)
abline(h=rowmeanserror,col="red")


df[,3:26]<-knnImputation(df[,3:26],k=15)


################################################################################
################################################################################
############# Normalize data #########################################
################################################################################
################################################################################
################################################################################
################################################################################

points<-c(0.1,0.25,0.1,0.25,0.1,0.25,0.1,0.25,0.1,0.25,0.1,0.25,
          0.1,0.25,0.1,0.25,0.1,0.25,0.1,0.25,0.1,0.25,0.1,0.25)

y_geomeans<-c()
for (n in c(3:26)) {
  y_geomeans<-append(y_geomeans,geoMean(df[,n]),after = length(y_geomeans))
    
}

y_means<-c()
for (n in c(3:26)) {
  y_means<-append(y_means,mean(df[,n]),after = length(y_means))
  
}

y_median<-c()
for (n in c(3:26)) {
  y_median<-append(y_median,median(df[,n]),after = length(y_median))
  
}

y_mode<-c()
for (n in c(3:26)) {
  y_mode<-append(y_mode,mlv(df[,n],method = "naive"),after = length(y_mode))
  
}

par(mfrow=c(4,6))

q=1
for (n in c(3:26)) {
  hist(df[,n],30,col="grey",xlim=c(0,0.4))
  abline(v=y_geomeans[q],col="cyan")
  abline(v=y_means[q],col="green")
  abline(v=y_median[q],col="red")
  abline(v=y_mode[q],col="purple")
  q=q+1
  
  
}

########### Seperate Young from old ####################
young<-df[,3:14]
old<-df[,15:26]
points<-c(0.1,0.25,0.1,0.25,0.1,0.25,0.1,0.25,0.1,0.25,0.1,0.25)
dat_young<-as.data.frame(cbind(points,y_geomeans[1:6]))
dat_old<-as.data.frame(cbind(points,y_geomeans[7:12]))


model.nls_young<-nls(V2~ ((1-points)*(f))+points, data=dat_young,start=list(f=0))
model.nls_old<-nls(V2~ ((1-points)*(f))+points, data=dat_old,start=list(f=0))
avevalue_young<-tidy(model.nls_young)$estimate
avevalue_old<-tidy(model.nls_old)$estimate
globalerror_young<-tidy(model.nls_young)$std.error
globalerror_old<-tidy(model.nls_old)$std.error
NormFacs<-predict(model.nls,x=dat$points)
NormFacs<-y_geomeans/NormFacs
df[,3:26] <- sweep(df[3:26], 2, NormFacs, FUN = "/")

plot(dat_old$points,dat_old$V2,col=rgb(1,0,0,0.3),pch=16,xlim=c(0,0.25),ylim=c(0,0.3))
points(dat_young$points,dat_young$V2,col=rgb(0,0,1,0.3),pch=16)
lines(c(0,dat_young$points),c(avevalue_young,predict(model.nls_young)),col="blue")
lines(c(0,dat_old$points),c(avevalue_old,predict(model.nls_old)),col="red")
abline(v=0)
abline(h=0)

#### Diff between young and old is nonsignificant so normaliza together ############
points<-c(0.1,0.25,0.1,0.25,0.1,0.25,0.1,0.25,0.1,0.25,0.1,0.25,
          0.1,0.25,0.1,0.25,0.1,0.25,0.1,0.25,0.1,0.25,0.1,0.25)

dat<-as.data.frame(cbind(points,y_geomeans))
model.nls<-nls(y_geomeans~ ((1-points)*(f))+points, data=dat,start=list(f=0))
NormFacs<-predict(model.nls,x=dat$points)
NormFacs<-y_geomeans/NormFacs
dft<-df
dft[,3:26] <- sweep(df[3:26], 2, NormFacs, FUN = "/")
boxplot(df[,3:26],outline=FALSE,notch=TRUE,col=rep(c("red","blue"),12),las=2)
abline(h=0.139012,col="red",lty=2)
abline(h=0.282510,col="blue",lty=2)
boxplot(dft[,3:26],outline=FALSE,notch=TRUE,col=rep(c("red","blue"),12),las=2)
abline(h=0.139012,col="red",lty=2)
abline(h=0.282510,col="blue",lty=2)

# Make stat matrix#
df<-read.table("C://Users/jbett/Desktop/mousemobbnormalized_technicalcombined.txt",sep="\t",header = TRUE)
nc<-nls.control(minFactor = 1/4096,warnOnly = TRUE)

points<-c(0.1,0.25,0.1,0.25,0.1,0.25)
for (n in 1:nrow(df)) {
  
  y_young<-as.numeric(df[n,1:6])
  dat_young<-as.data.frame(cbind(points,y_young))
  dat_young$y_young<-unlist(dat_young$y_young)
  dat_young$points<-unlist(dat_young$points)
  model.nls_young<-nls(y_young ~ ((1-points)*(f))+points, data=dat_young,start=list(f=0),control = nc)
  df$value_young[n]<-tidy(model.nls_young)$estimate
  df$stderror_young[n]<-tidy(model.nls_young)$std.error
  df$cor_young[n]<-nrmse_func(y_young,predict(model.nls_young,x=points),type="mean")
  
  y_old<-as.numeric(df[n,7:12])
  dat_old<-as.data.frame(cbind(points,y_old))
  dat_old$y_old<-unlist(dat_old$y_old)
  dat_old$points<-unlist(dat_old$points)
  model.nls_old<-nls(y_old~ ((1-points)*(f))+points, data=dat_old,start=list(f=0),control = nc)
  df$value_old[n]<-tidy(model.nls_old)$estimate
  df$stderror_old[n]<-tidy(model.nls_old)$std.error
  df$cor_old[n]<-nrmse_func(y_old,predict(model.nls_old,x=points),type="mean")
  
  
}


df<-df[df$cor_old<=0.2,]
df<-df[df$cor_young<=0.2,]


df$teststatistic_young<-(df$value_young-median(df$value_young))/df$stderror_young
df$pvalue_young<-pt(-abs(df$teststatistic_young),df =5)

hm_young<-ash((df$value_young-median(df$value_young)),df$stderror_young,df = 5,mixcompdist = "uniform")
df$p_adjusted_ash_uniform_young<-hm_young[["result"]][["qvalue"]]

hm_young<-ash((df$value_young-median(df$value_young)),df$stderror_young,df = 5,mixcompdist = "+uniform")
df$p_adjusted_ash_halfuniform_young<-hm_young[["result"]][["qvalue"]]


df$p_adjusted_young_BH<-p.adjust(df$pvalue_young,method="BH")
df$p_adjusted_young_BY<-p.adjust(df$pvalue_young,method="BY")


df$teststatistic_old<-(df$value_old-median(df$value_old))/df$stderror_old
df$pvalue_old<-pt(-abs(df$teststatistic_old),df =5)

hm_old<-ash((df$value_old-median(df$value_old)),df$stderror_old,df = 5,mixcompdist = "uniform")
df$p_adjusted_ash_uniform_old<-hm_old[["result"]][["qvalue"]]

hm_old<-ash((df$value_old-median(df$value_old)),df$stderror_old,df = 5,mixcompdist = "+uniform")
df$p_adjusted_ash_halfuniform_old<-hm_old[["result"]][["qvalue"]]

df$p_adjusted_old_BH<-p.adjust(df$pvalue_old,method="BH")
df$p_adjusted_old_BY<-p.adjust(df$pvalue_old,method="BY")

df$sd_young<-df$stderror_young*sqrt(6)
df$sd_old<-df$stderror_old*sqrt(6)


df$pooled_variance<-(((6-1)*df$sd_old)+((6-1)*df$sd_young))/(6+6-2)
df$t_twosample<-(df$value_old-df$value_young)/(df$pooled_variance*sqrt((1/6)+(1/6)))
df$pvalue_twosample<-pt(-abs(df$t_twosample),df =10)

hm_two<-ash(df$value_old-df$value_young,df$pooled_variance,df = 10,mixcompdist = "uniform")
df$p_adjusted_two_BH<-p.adjust(df$pvalue_twosample,method="BH")


for (n in 1:nrow(df)) {
  ifelse(df$p_adjusted_ash_uniform_old[n]<=0.05,df$sig_ashuniform_old[n]<-TRUE,df$sig_ashuniform_old[n]<-FALSE)
}

for (n in 1:nrow(df)) {
  ifelse(df$p_adjusted_ash_uniform_young[n]<=0.05,df$sig_ashuniform_young[n]<-TRUE,df$sig_ashuniform_young[n]<-FALSE)
}

########### interage ################################
nc<-nls.control(minFactor = 1/4096,warnOnly = TRUE)

points<-c(0.1,0.25,0.1,0.25,0.1,0.25,0.1,0.25,0.1,0.25,0.1,0.25)
for (n in 1:nrow(df)) {
  
  y_all<-as.numeric(df[n,1:12])
  dat_all<-as.data.frame(cbind(points,y_all))
  dat_all$y_all<-unlist(dat_all$y_all)
  dat_all$points<-unlist(dat_all$points)
  model.nls_all<-nls(y_all ~ ((1-points)*(f))+points, data=dat_all,start=list(f=0),control = nc)
  df$value_all[n]<-tidy(model.nls_all)$estimate
  df$stderror_all[n]<-tidy(model.nls_all)$std.error
  df$cor_all[n]<-nrmse_func(y_all,predict(model.nls_all,x=points),type="mean")
}


df$teststatistic_all<-(df$value_all-median(df$value_all))/df$stderror_all
df$pvalue_all<-pt(-abs(df$teststatistic_all),df =11)

hm_all<-ash((df$value_all-median(df$value_all)),df$stderror_all,df = 5,mixcompdist = "uniform")
df$p_adjusted_ash_uniform_all<-hm_all[["result"]][["qvalue"]]

hm_all<-ash((df$value_all-median(df$value_all)),df$stderror_all,df = 5,mixcompdist = "+uniform")
df$p_adjusted_ash_halfuniform_all<-hm_all[["result"]][["qvalue"]]


for (n in 1:nrow(df)) {
  ifelse(df$p_adjusted_ash_uniform_all[n]<=0.05,df$sig_ashuniform_all[n]<-TRUE,df$sig_ashuniform_all[n]<-FALSE)
}


ggscatter(data=df,x="value_young",y="value_old",color  =ifelse(df$sig_ashhalf_all==TRUE,"red","grey"))+
  geom_abline(intercept = 0,slope = 1,lty=2,col="green")+
  geom_hline(yintercept = median(df$value_old),col="slateblue4",lty=2)+
  geom_vline(xintercept = median(df$value_young),col="slateblue4",lty=2)


for (n in 1:nrow(df)) {
  tensinyoung<-as.vector(unlist(df[n,c(1,3,5)]))
  twentyfivesinyoung<-as.vector(unlist(df[n,c(2,4,6)]))
  
  tensinold<-as.vector(unlist(df[n,c(7,9,11)]))
  twentyfivesinold<-as.vector(unlist(df[n,c(8,10,12)]))
  
  df$stderror_young_10s[n]<-standard_error(tensinyoung)
  df$stderror_young_25s[n]<-standard_error(twentyfivesinyoung)
  
  df$stderror_old_10s[n]<-standard_error(tensinold)
  df$stderror_old_25s[n]<-standard_error(twentyfivesinold)
}


errormat<-df[,c(16,19,36,37,38,39)]
```
